# Functional Differentiation of GH172 Arabinofuranosidases Through Divergent Quaternary Structures

**DOI:** 10.64898/2026.08.13.744632

**Authors:** Jennifer Ross, Moescha J. Hoopman, Florian Küllmer, Omar Al-Jourani, Augustinas Silale, Morwan M. Osman, Zongjia Chen, Hannah R. Bridges, Kaspar J. Garnham, Carl Morland, Jean-Lou Reyre, Abigail J. Layton, Johan Turkenburg, Sam Hart, Alexandra Solovyova, Andrew Porter, Arnaud Baslé, Jeroen D. C. Codée, Spencer J. Williams, Patrick J. Moynihan, Herman S. Overkleeft, James N. Blaza, Elisabeth C. Lowe

**Author notes:** To whom correspondence should be addressed: Elisabeth Lowe, Patrick Moynihan, James Blaza, Herman Overkleeft.

## Abstract

Mycobacteria synthesise the unusual glycan ᴅ-arabinan as a major component of the cell wall glycoconjugates arabinogalactan (AG) and lipoarabinomannan (LAM). We previously identified *Dysgonomonas gadei*, a member of the Bacteroidota, as capable of complete ᴅ-arabinan degradation through the concerted action of endo- and exo-acting enzymes. Among these are three glycoside hydrolase family 172 (GH172) enzymes with exo-α-ᴅ-arabinofuranosidase activity against AG and LAM, although their linkage specificities were unknown. Here, using defined synthetic substrates, we show that the three enzymes possess distinct linkage preferences. We also develop α-ᴅ-arabinofuranosyl cyclophellitol aziridines as covalent inhibitors and activity-based probes for GH172 enzymes. X-ray crystallography and cryo-EM to reveal strikingly different quaternary assemblies across the three homologues, while a 1.5 Å cryo-EM structure of dodecameric Dg67 covalently modified by an aziridine inhibitor identifies the catalytic nucleophile and provides direct structural support for a retaining mechanism. A BODIPY-tagged aziridine probe selectively labelled the three GH172 enzymes in *D. gadei* cell lysates. Together, these findings define functional and structural diversity within GH172 and establish chemical probes for profiling α-ᴅ-arabinofuranosidase activity in complex biological samples.

## Introduction

The cell wall of acid-fast bacteria, including *Mycobacterium tuberculosis*, contains complex glycans built in part from the unusual monosaccharide ᴅ-arabinofuranose (ᴅ-Ara*f*). ᴅ-Ara*f* is a prominent component of the essential mycobacterial glycans and lipoglycans, arabinogalactan (AG) and lipoarabinomannan (LAM) (**Figure 1A**).^1^ Despite the abundance of mycobacterial ᴅ-Ara*f*-containing glycans, the first examples of enzymes able to degrade these species were only reported in the past five years.^2,3^ Members of glycoside hydrolase (GH) families GH183 and GH172 from several bacterial species were shown to possess endo-D-arabinanase and exo-D-arabinofuranosidase activity, respectively.^2–4^ Unexpectedly, we found that the gut Bacteroidota species, *Dysgonomonas gadei*, can degrade LAM and AG. We identified a polysaccharide utilisation locus (PUL) in this species encoding three GH172 enzymes (Dg67, Dg79, and Dg71; **Figure 1B**) and presented evidence that these proteins assemble into distinct oligomeric states.^2^

**Figure 1:**
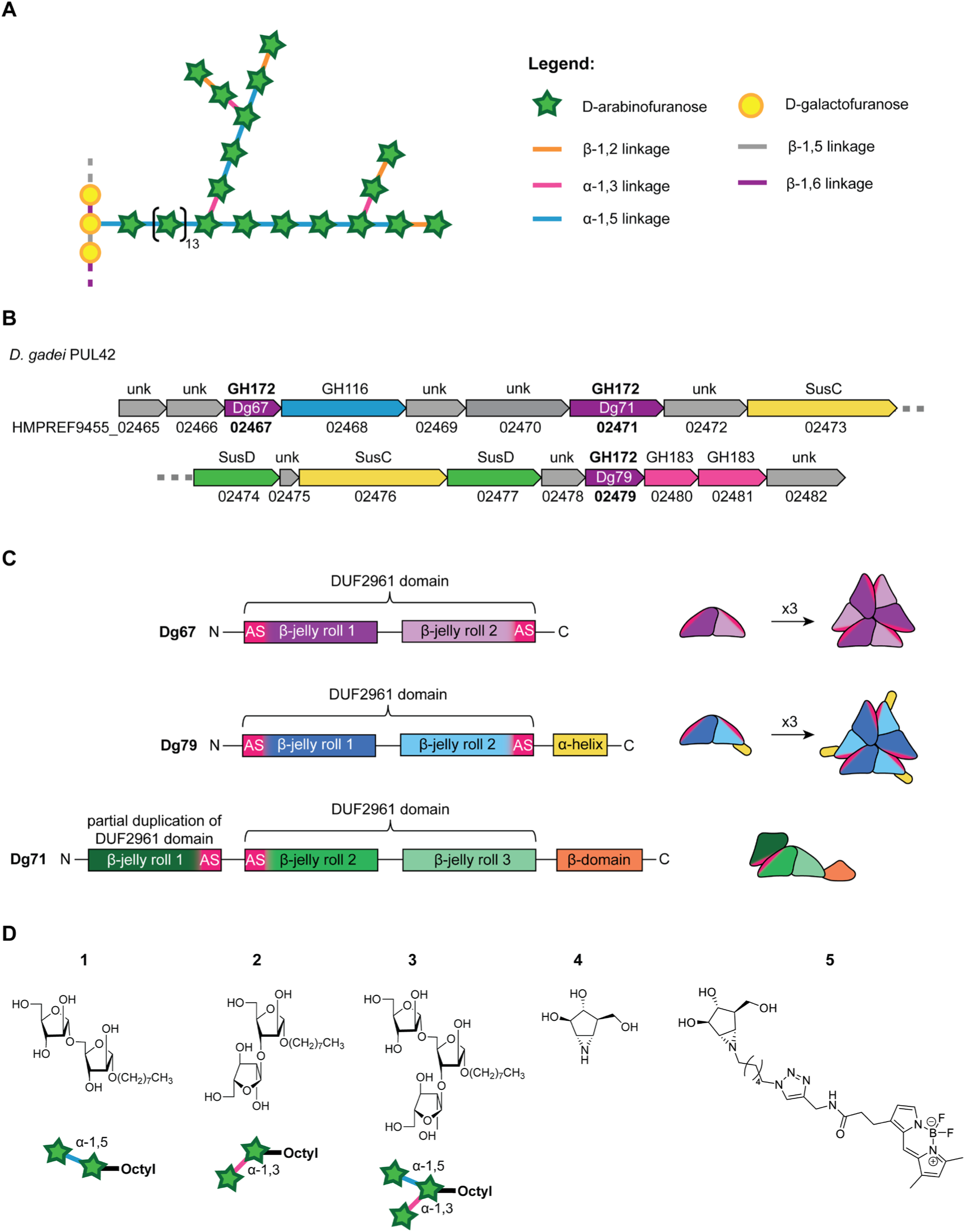
GH172 enzymes and their substrates. **A**: Simplified representation of the ᴅ-arabinan branches of mycobacterial AG. **B**: Schematic representation of the arabinan-degrading PUL42 from *D. gadei*. **C**: Domain architectures of Dg67, Dg79, and Dg71, with jelly roll domains shown in shades of purple, blue and green respectively, and regions contributing active site residues (AS) shown in pink. Cartoon representations to the right illustrate how oligomerisation of jelly-roll folds (purples, blues, and greens) completes the active sites at protomer interfaces. **D**: Chemical structures of the synthetic α-ᴅ-arabinofuranosyl di- and trisaccharide substrates **1**–**3** and α-ᴅ-arabinofuranosyl cyclophellitol aziridines **4** and **5**.

Members of family GH172 (PFAM DUF2961) are unique among glycoside hydrolase families in adopting a double β-jelly-roll (DJR) fold and are evolutionarily related to the DJR major capsid proteins of viruses in the kingdom *Bamfordvirae.*^5^ We have identified two catalytically competent DJR architectures in GH172 enzymes. In the first DJR configuration, exemplified by Dg67 and Dg79, each protomer contains a single DJR domain, and the active site is formed at the interface between neighbouring protomers, making oligomerization essential for activity. The splitting of active site residues across different protomers is also observed in a GH129 enzyme where two protomers form a homodimer and make two active sites.^6^ The GH172 catalytic residues are distributed across the N- and C-terminal ends of each DJR domain, and because of geometric constraints, at least three protomers are required to assemble a complete set of productive active sites. As such, the trimer represents the minimal functional catalytic unit (**Figure 1C**) as a dimer of the DJR cannot complete both active sites. These prerequisite trimers can associate into higher-order assemblies, including hexamers, formed as dimers of trimers, and dodecamers, formed as tetramers of trimers. In the second DJR architecture, exemplified by Dg71, partial duplication of the DUF2961 domain within a single gene gives rise to an additional β-jelly-roll domain. In this arrangement, the active site is completed within a single polypeptide, and therefore oligomerization is not required for catalytic activity (**Figure 1C**, lower schematic). 3D structures of first DJR configuration have been determined for three GH172 members to date, two hexamers (PDB: 4KQ7, unpublished; PDB: 7V1V, a difructose dianhydride I synthase/hydrolase known as αFFase1) and a dodecameric exo-ᴅ-arabinofuranosidase (PDB: 8IC8).^3,4^ The 3D structure of the second type of DJR architecture, corresponding to Dg71 remains unknown. Moreover, owing to a lack of substrate bound structures, it is unclear how the two GH172 DJR architectures organise their interfacial active sites and catalytic machinery.

Here, we define the functional and structural diversity of three GH172 enzymes, Dg67, Dg79 and Dg71, encoded within a polysaccharide utilisation locus of *D. gadei*. Using synthetic ᴅ-arabinofuranosyl oligosaccharides with defined glycosidic linkages representing substructures of AG and LAM (compounds **1–3**, **Figure 1D**), we show that the three enzymes possess distinct linkage preferences despite sharing exo-α-ᴅ-arabinofuranosidase activity and a conserved retaining catalytic apparatus. We also prepared α-ᴅ-arabinofuranosyl cyclophellitol aziridines **4** and **5** (**Figure 1D**) as covalent inhibitors and activity-based probes for profiling GH172 enzymes *in vitro* and in cell lysates. Structural analysis by X-ray crystallography and cryo-EM revealed markedly different oligomeric assemblies and active-site architectures built around a conserved double-jelly-roll catalytic framework. Finally, a 1.5 Å cryo-EM structure of dodecameric Dg67 covalently modified by **4** identifies the catalytic nucleophile, resolves the opened aziridine and its interactions with the catalytic acid/base, and provides direct structural support for the retaining mechanism of GH172 enzymes. Together, these studies show how a conserved GH172 catalytic framework can accommodate substantial variation in quaternary structure and substrate specificity, while establishing α-ᴅ-arabinofuranosyl cyclophellitol aziridines as chemical tools for investigating this unusual family of glycosidases.

## Results

In our initial characterization of the three *D. gadei* GH172 enzymes it was observed that the three enzymes differed in size, solution-phase oligomerisation, and activities. Dg67 and Dg79 were more active than Dg71 on mycobacterial ᴅ-arabinans, suggesting differing substrate selectivity.^2^ Furthermore, size-exclusion chromatography-light scattering (SEC-LS) identified Dg71 as a dimer, Dg79 as a hexamer, and Dg67 as a dodecamer. To assess whether these differences in oligomerisation states may influence substrate selectivity and function we set out to determine their 3D structures and to systematically study their substrate activities using defined substrates and inhibitors.

Dg67 shares 15.1% and 27.7% sequence identity with Dg71 and Dg79 respectively. Dg79 and Dg71 share 15.7% sequence identity with one another. Despite this low overall sequence identity, the catalytic residues are strictly conserved in these three GH172 enzymes (**Figure S1**). Multiple homologues belonging to the same GH family may evolve and be retained provided they confer a fitness benefit, such as different functional roles in terms of substrate or cellular location. ^2,7^ Models for PUL-mediated breakdown of complex glycans typically invoke the initial action of endo-active enzymes on the cell surface, which create oligosaccharides from larger polysaccharides. These smaller oligosaccharides are transported into the periplasm by TonB-dependent transport systems where they undergo further degradation by exo-glycosidases, or in some cases additional endo-glycosidases. Proteomics of ᴅ-arabinan grown *D. gadei* detected all three GH172 enzymes, suggesting they are not functionally redundant and may act on related substrates whilst differing in substrate preference or biological role.^2^ Furthermore, SignalP 6.0 analysis of the sequences of the three *D. gadei* GH172 members predicts that all three are located within the periplasm, suggesting a possible role in oligosaccharide degradation.^8^

### Linkage preferences of the three *D. gadei* GH172 enzymes

The differing activities of Dg67, Dg71, and Dg79 on mycobacterial AG suggested that the enzymes may recognize distinct structural features within arabinan. However, the complex architecture of AG, together with the heterogeneity introduced during cell-wall extraction, prevents the glycosidic linkages cleaved by each enzyme from being assigned using the native polymer alone (**Figure 1A**). We therefore examined their activities using defined substrates to establish their linkage preferences. Such differences are plausible given the substrate diversity already reported with family GH172. For example, the family member αFFase1 acts on α-ᴅ-Fru*f*-1,2′:2,1′-β-ᴅ-Fru*f* (difructose dianhydride I [DFA I]), catalyzing a reversible reaction that establishes an equilibrium between DFA I and inulobiose.^4^ We therefore tested DFA I as a substrate for the three *D. gadei* GH172 proteins, but none showed detectable activity (**Figure S2**). A second source of substrates with defined linkages is the pili of *Pseudomonas aeruginosa* PA7, which are adorned with α-1,5-linked ᴅ-Ara*f* residues. In our previous study, Dg67 and Dg79, but not Dg71, degraded these oligosaccharides to ᴅ-arabinose.^2,9^ This difference suggested that Dg71 differs in linkage preference from Dg67 and Dg79. To test this directly, we synthesized three oligosaccharides containing defined glycosidic linkages (compounds **1**–**3**, **Figure 1D**)^10,11^.

Dg67 completely degraded compounds **1**, **2**, and **3** to ᴅ-arabinose (**Table S1**), demonstrating that it can hydrolyse both α-1,5 and α-1,3 linkages, including within a branched substrate (**Figure 2**, purple traces). The ᴅ-Ara*f*-octyl reducing-end moiety of each substrate is not detected by ion chromatography-pulsed amperometric detection (IC-PAD); consequently, the measured ᴅ-Ara*f* concentrations and IC-PAD peak areas do not account for this portion of the cleaved substrate. By contrast, Dg79 completely degraded compound **1**, which contains an α-1,5 linkage, but showed only weak activity towards compound **2**, which contains an α-1,3 linkage, indicating a preference for α-1,5-linked ᴅ-Ara*f*. Dg79 also partially degraded the branched trisaccharide **3** (**Figure 2**, blue traces). IC-PAD analysis revealed formation of a new product that co-eluted with compound **2**, consistent with cleavage of the 1,5-linkage while leaving the 1,3 linkage intact (**Figure S3**). Accordingly, Dg79 released a similar amount of ᴅ-Ara*f* to Dg67 from compound **1**, but substantially less from compounds **2** and **3** (**Table S1**). Dg71 showed only trace activity towards compounds **2** and **3** and no detectable activity against **1** (**Figure 2**, green traces; **Table S1**). Its lack of activity against compound **1** is consistent with our previous observation that Dg71 does not degrade the α-1,5-ᴅ-Ara*f* oligosaccharides decorating the pili of *P. aeruginosa* PA7.

**Figure 2:**
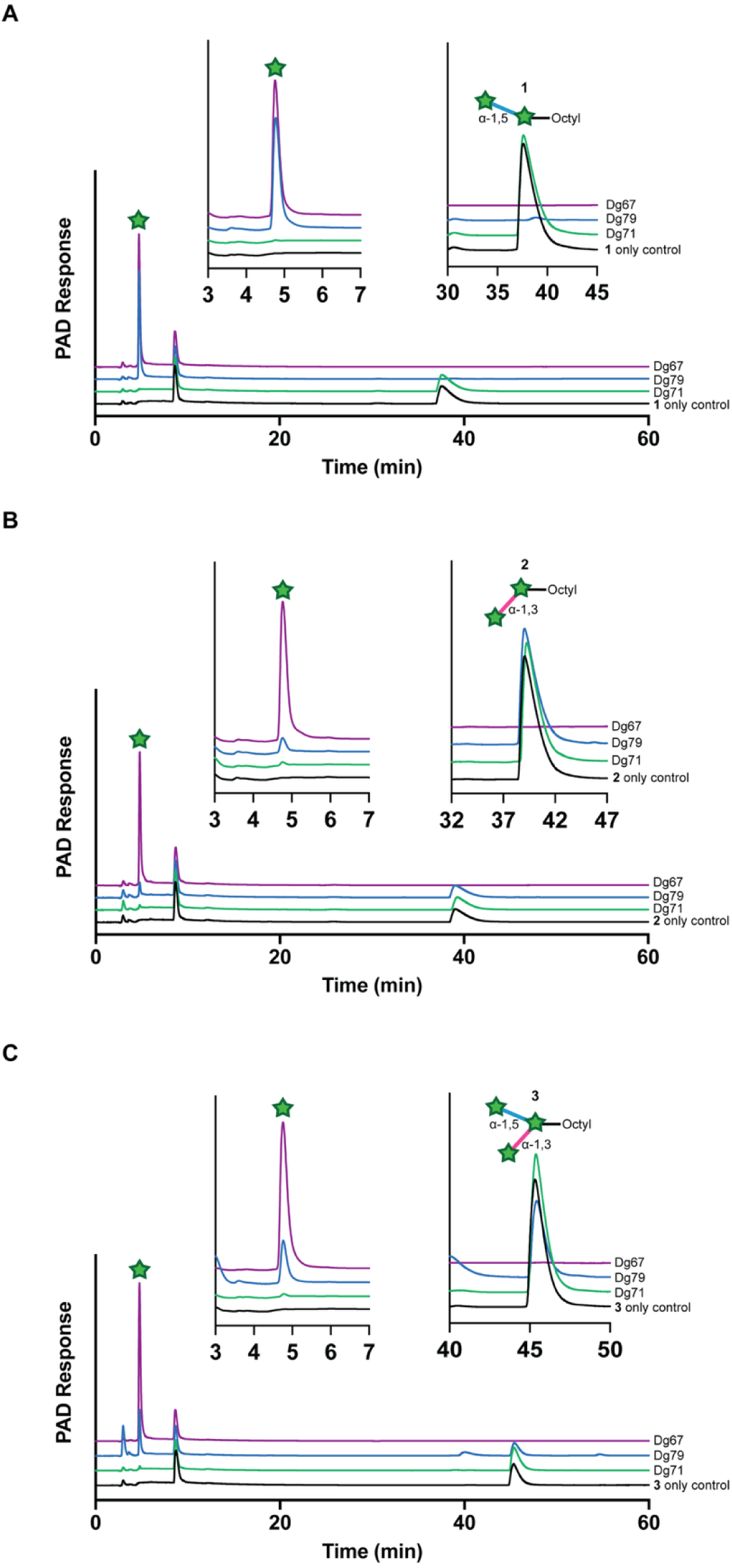
*D. gadei* GH172 enzymes are exo-ᴅ-arabinofuranosidases with distinct linkage preferences. Dg67, Dg79, and Dg71 (1 µM) were incubated with 500 µM of compound **1** (**A**), compound **2** (**B**), or compound **3** (**C**) in 50 mM phosphate–citrate buffer, pH 7.0. Reaction products were analysed by ion chromatography with pulsed amperometric detection (IC-PAD).

Together, these results reveal marked functional divergence among the three *D. gadei* GH172 enzymes, despite each exhibiting exo-ᴅ-arabinofuranosidase activity against AG. Dg67 is a broadly active exo-α-arabinofuranosidase that cleaves both α-1,5- and α-1,3-linkages, whereas Dg79 is a highly selective exo-α-1,5-arabinofuranosidase. In contrast, Dg71 exhibits only weak exo-α-1,3-arabinofuranosidase activity, suggesting that its preferred substrate or linkage was not represented among those tested. Thus, the three enzymes possess distinct substrate specificities that may reflect their different oligomeric assemblies, with Dg67 uniquely capable of cleaving both α-linkage types found in AG.

### Structures of the *D. gadei* GH172 enzymes

In our previous study, SEC-LS analysis showed that Dg67, Dg71 and Dg79 adopt distinct oligomerization states in solution: Dg67 forms a dodecamer, Dg79 a hexamer, and Dg71 a dimer.^2^ To define the structural basis for these different assemblies and to investigate the origins of their differing substrate preferences, we determined the structures of all three GH172 homologues using cryo-EM and X-ray crystallography. Previous structural studies of GH172 enzymes have described an apo dodecamer structure at medium resolution and a high-resolution hexameric enzyme, αFFase1, with difructose dianhydride I synthase/hydrolase activity.^4^ However, αFFase1 has a C-terminal architecture distinct from that of Dg79, and no structures of dimeric GH172 enzymes have previously been reported.

### Cryo-EM structure of Dg71

To define the structural basis of the dimeric GH172 assembly, we determined the structure of Dg71 by single-particle cryo-EM. Imposing C2 symmetry yielded a reconstruction with a global resolution of 3.6 Å (**Figures 3** and **S4, Tables S2/S3**). The conical FSC (cFAR) plot for Dg71 has a cFAR value of 0.33, indicating some orientation bias. This explains the over-estimation in resolution for Dg71 seen with FSC estimated resolution (**Figure S4E**). Each Dg71 protomer comprises three β-jelly roll domains followed, after a linker region, by a fourth β-sheet domain. The third β-jelly-roll contributes residues that complete the active site within a single polypeptide chain (**Figure 3**). Thus, unlike other characterized GH172 enzymes, Dg71 does not require oligomerization to form a complete active site. Instead, dimerization occurs through an interface formed between the fourth β-sheet domains of the two protomers (orange, **Figure 3**).

**Figure 3:**
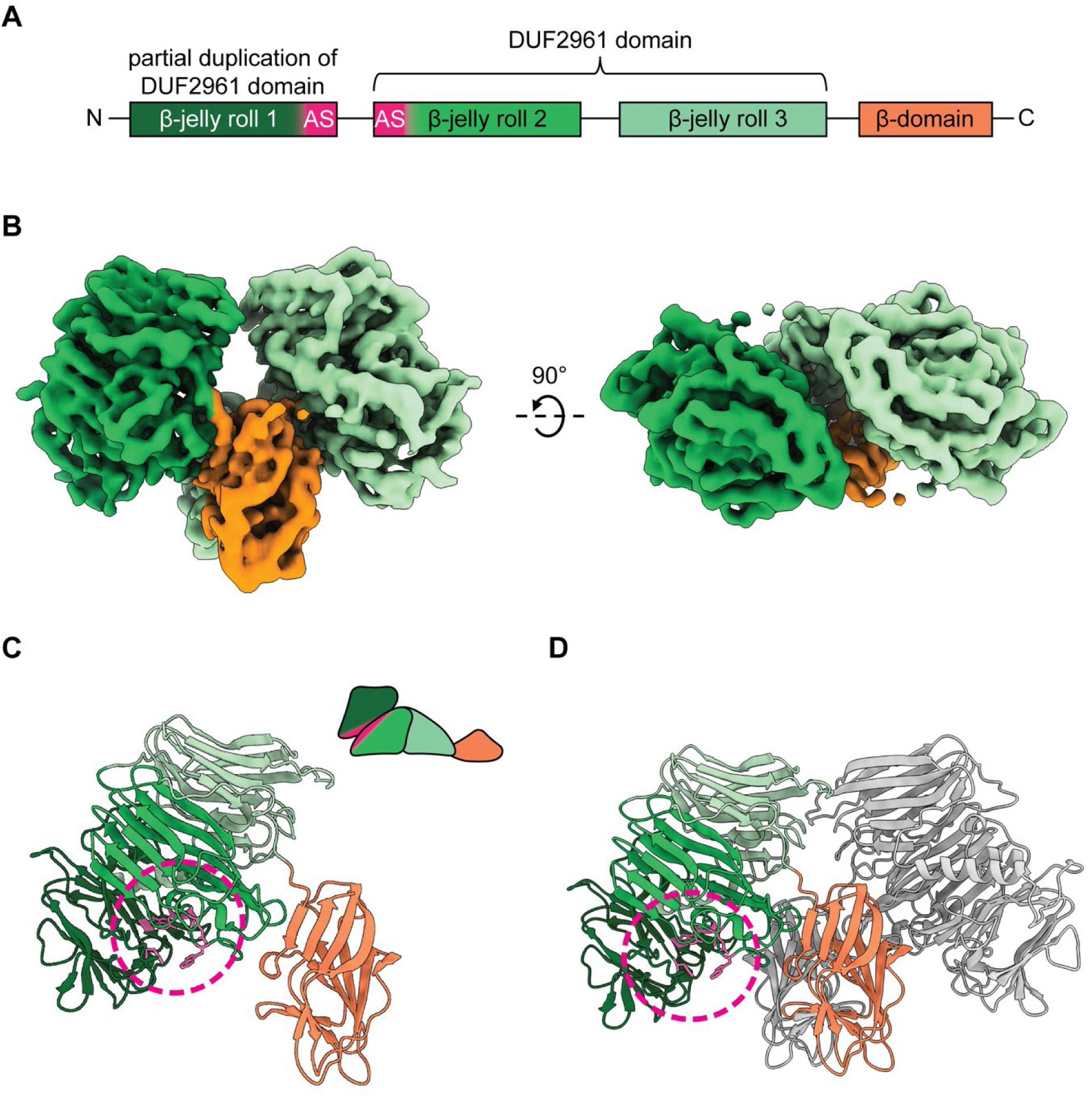
The cryo-EM structure of Dg71. **A:** Domain schematic of Dg71. **B:** Cryo-EM map of the Dg71 dimer, with the two protomers coloured different shades of green and the fourth β-sheet domain in orange, as shown in the domain schematic. The map is contoured at a level of 0.098. **C,D**: Cartoon representations of the Dg71 protomer (**C**) and dimer (**D**). The C-terminal β-sheet domain that mediates dimerization is highlighted in orange. The active site is shown in pink in panels **A**, **C** and **D**.

### **X-** ray crystallography structure of Dg79

We determined the X-ray crystal structure of Dg79 by molecular replacement using the structure of an unpublished GH172 enzyme from *Bacteroides uniformis* (PDB: 4KQ7) as the search model. Data collection and refinement statistics are provided in **Table S4**. The structure was refined to 1.4 Å resolution (PDB: 8AH3).

Each Dg79 protomer adopts a DJR fold with an additional C-terminal α-helix that is absent from the other *D. gadei* GH172 enzymes (**Figure 4A/B**). Six protomers assemble into a dimer-of-trimers, with the interface between the two trimers formed by interlocking C-terminal α-helices (**Figure 4C/D/E**). Seven coordinated calcium ions are present within each trimer: two associated with the active site, and one positioned on the three-fold symmetry axis. The calcium ions are coordinated by D235, which is not conserved in the other *D. gadei* GH172 enzymes. Their assignment as calcium was supported by the coordination geometry, anomalous scattering and metal-ligand bond distances. A calcium ion is also present in the active site of αFFase1, where it contributes to substrate binding through a water-mediated interaction.^4^ Calcium ions similarly assist substrate recognition or catalysis in several other GH families, including GH43 α-L-arabinanases, GH62 α-L-arabinofuranosidases, and GH47 and GH92 α-mannosidases.^12–15^ The dimer-of-trimers quaternary structure is not unique to Dg79 within GH172, as a related assembly is observed for αFFase1.^4^ However, the two enzymes possess distinct C-terminal regions, resulting in differences in the relative arrangement of the trimers and in accessibility of the active site (**Figure S5**).

**Figure 4:**
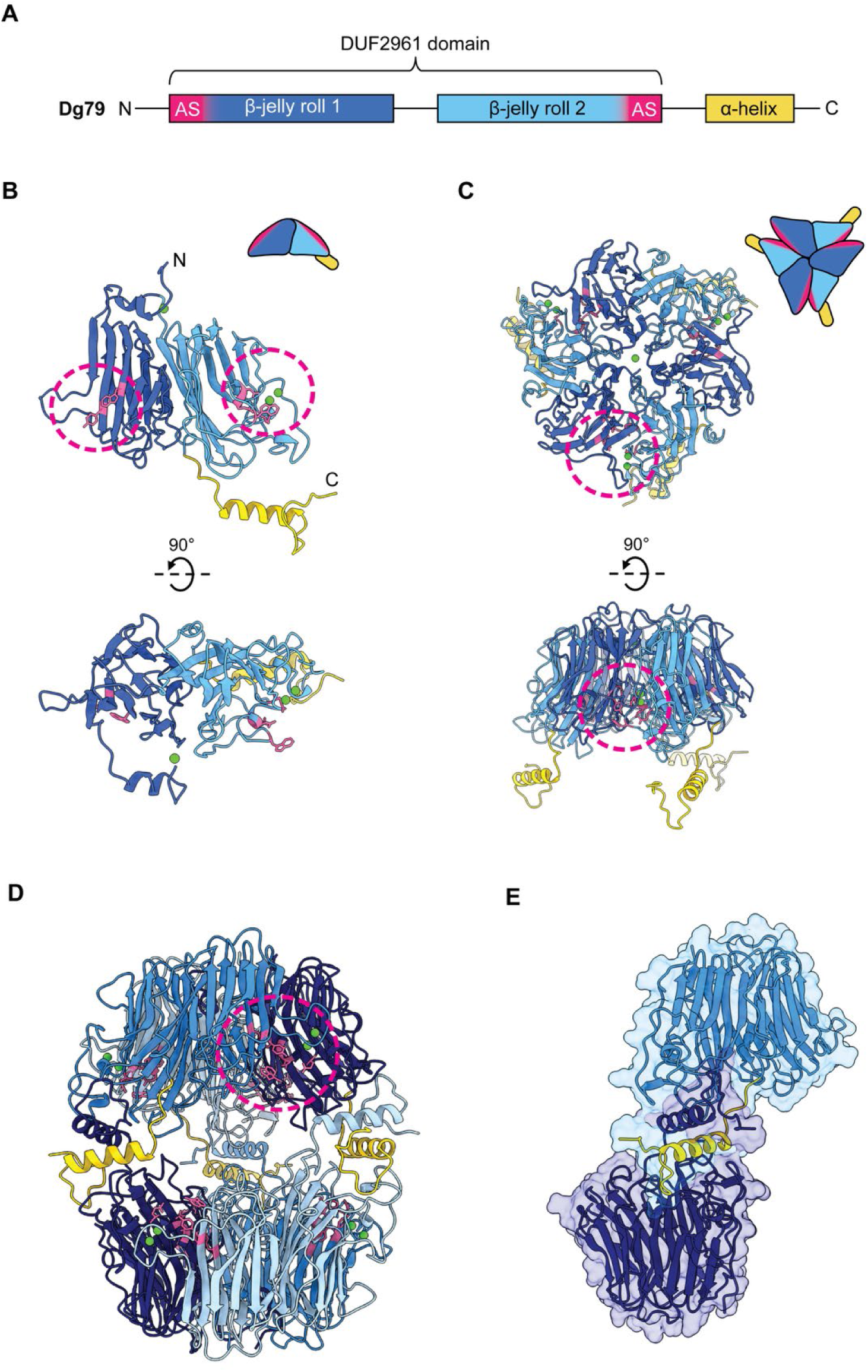
X-ray crystal structure of Dg79. **A**: Domain architecture of Dg79. **B–D**: Cartoon representations of the Dg79 protomer (**B**), trimer (**C**), and dimer-of-trimers quaternary structure (**D**). Active sites are circled in pink, and bound calcium ions are shown as green spheres. The protomer in **B** and subunits in **C** are coloured according to the domain schematic, whereas the six protomers in **D** are shown in different shades of blue for clarity. **E**: Transparent solvent-accessible surfaces overlaid on cartoon representations of protomers from opposing trimers, showing the interlocking C-terminal helices that form the trimer-trimer interface (yellow).

### Cryo-EM structure of Dg67

Given the high molecular weight of dodecameric Dg67 (approximately 520 kDa), we determined its apo structure by single-particle cryo-EM (**Figure S6**). Consistent with the previously reported X-ray crystal structure of the dodecameric GH172 enzyme ExoMA1,^3^ Dg67 displayed tetrahedral (T) symmetry which was imposed during refinement (**Figure 5** and **Tables S5/S6**). The resulting reconstruction has a global resolution of 2.1 Å (**Figure S6C**). The reconstruction reveals 12 protomers arranged as a tetramer-of-trimers (**Figure 5C** and **D**). Each protomer adopts a DJR fold. Assembly of the four trimers into the dodecamer is mediated by contacts involving four surface loops and the C-terminal region of each protomer (**Figure 5E**). As observed for ExoMA1, additional density consistent with a phosphate ion is present at the interfaces between trimers, where it may contribute to assembly or stabilisation of the dodecamer.

**Figure 5:**
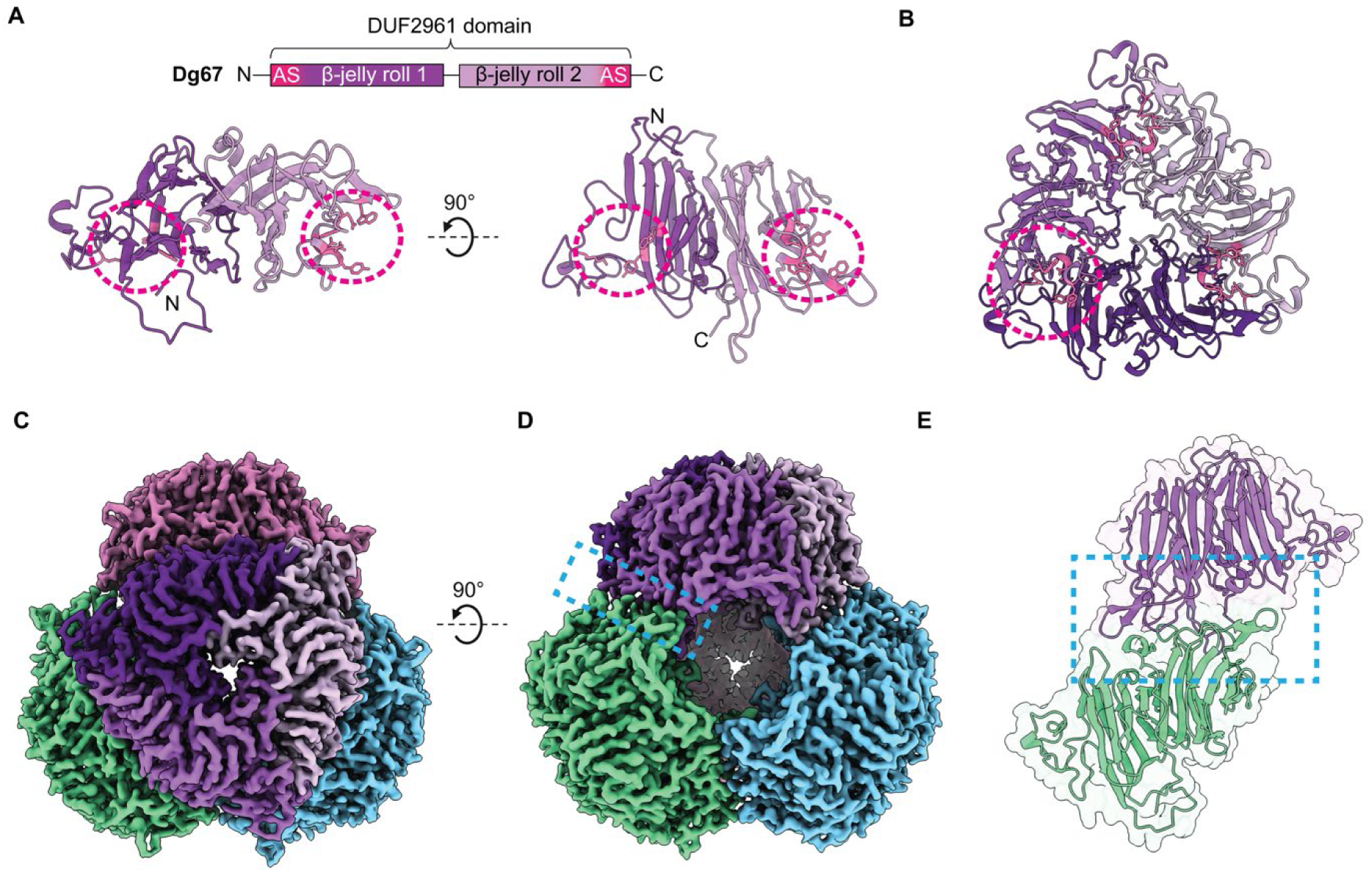
Cryo-EM structure of Dg67. **A**: Domain architecture of Dg67 and cartoon representation of a Dg67 protomer coloured according to the domain schematic. The two regions that together form the active site are circled in pink. **B**: Cartoon representation of the Dg67 trimer, with the three protomers shown in different shades of purple. **C,D:** Cryo-EM maps of the Dg67 dodecamer. In **C,** one trimer is coloured as in **B**, whereas the remaining trimers are coloured pink, green and blue to illustrate the tetramer-of-trimers assembly. The maps are contoured at a level of 0.037. **E**: Cartoon representation of two protomers from adjacent trimers, highlighting the loops that mediate the trimer–trimer interface. The location of this interface is indicated by a blue box in **D** and shown in detail in **E**.

### Arabinose bound structure of Dg67

The structure of Dg67 in complex with its product, ᴅ-arabinose, was determined at a global resolution of 2.6 Å (**Figure S7**). Density corresponding to ᴅ-arabinose in its furanose form was observed in the active sites of all 12 subunits, occupying the putative −1 subsite (**Figure S8A/B**). Because the catalytic residues are conserved among Dg67, Dg71 and Dg79, the Dg67 bound ᴅ-Ara*f* was superimposed into the active sites of Dg71 and Dg79. In both cases the product could be accommodated without steric clashes (**Figure S8C/D**).

Despite extensive efforts, attempts to obtain substrate-bound structures of Dg67 and Dg79 did not produce interpretable ligand density in their active sites. However, these experiments did yield an apo cryo-EM reconstruction of Dg79 (**Figure S9** and **Figure S10A**). The Dg79 crystal structure fitted well within the cryo-EM map (**Figure S10B**). However, density for the loop spanning E207–N221, which lies adjacent to the active site, was poorly defined and could not be modelled confidently (**Figure S10C**). Additional density was observed near C285, consistent with a previously reported spontaneous cysteine modification arising as an artefact of cryo-EM grid preparation (**Figure S10D**)^16^.

### α-ᴅ-Arabinofuranosyl cyclophellitol-based probes Inhibition with aziridine 4

Cyclophellitol-based inhibitors are electrophilic reagents that target the catalytic nucleophile of retaining glycoside hydrolases, forming a covalent enzyme-inhibitor adduct and thereby causing irreversible inhibition.^17^ To investigate the susceptibility of the GH172 enzymes to this class of inhibitor, we synthesized the α-ᴅ-arabinofuranose-configured cyclophellitol aziridine **4** (**Figure S11**).

Preincubation of Dg67 with **4** resulted in loss of activity against pNP-α-ᴅ-Ara*f*, consistent with inhibition through covalent labelling. To determine the enzyme-probe stoichiometry, each GH172 enzyme was incubated with **4** in a 1:1 molar ratio and analysed by intact protein mass spectrometry (**Figure S12**). After incubation for 1 h at 37 °C, Dg67, Dg79, and Dg71 each exhibited a mass increase consistent with formation of a single covalent adduct with **4**, with only negligible amounts of unmodified protein remaining. The efficient labelling of all three enzymes supports a retaining catalytic mechanism involving a covalent glycosyl–enzyme intermediate and establishes aziridine-based cyclophellitols as useful chemical probes for GH172 enzymes. Notably, all three homologues were labelled despite their differing substrate specificities. In combination with the activity against pNP-α-ᴅ-Ara*f*, this result shows that the arabinofuranosyl-configured glycon can be accommodated in each active site in the absence of an extended aglycon and suggests that their differing activities arise primarily from recognition of the glycosidic linkage and surrounding substrate architecture.

### Activity-Based Probes (ABPs) for profiling temperature and pH dependence

We next examined labelling of the GH172 enzymes with the BODIPY-tagged aziridine **5** (**Figure S13**). To determine whether **5** could detect the three *D. gadei* enzymes in a complex biological sample, *D. gadei* was grown on either ᴅ-glucose or ᴅ-arabinan, prepared from *Mycobacterium smegmatis* arabinogalactan by enzymatic removal of the galactan component, as described previously.^2^ Whole cell lysates were incubated with **5** (10 μM) and analysed by SDS-PAGE and in-gel fluorescence detection (**Figure 6A**). Bands corresponding to Dg67 (44.1 kDa), Dg71 (73.9 kDa) and Dg79 (46.0 kDa) were detected only in lysates from cells grown on ᴅ-arabinan. These results indicate that all three enzymes are expressed during growth on arabinan and demonstrate the utility of this probe for profiling GH172 activity in complex samples.

**Figure 6:**
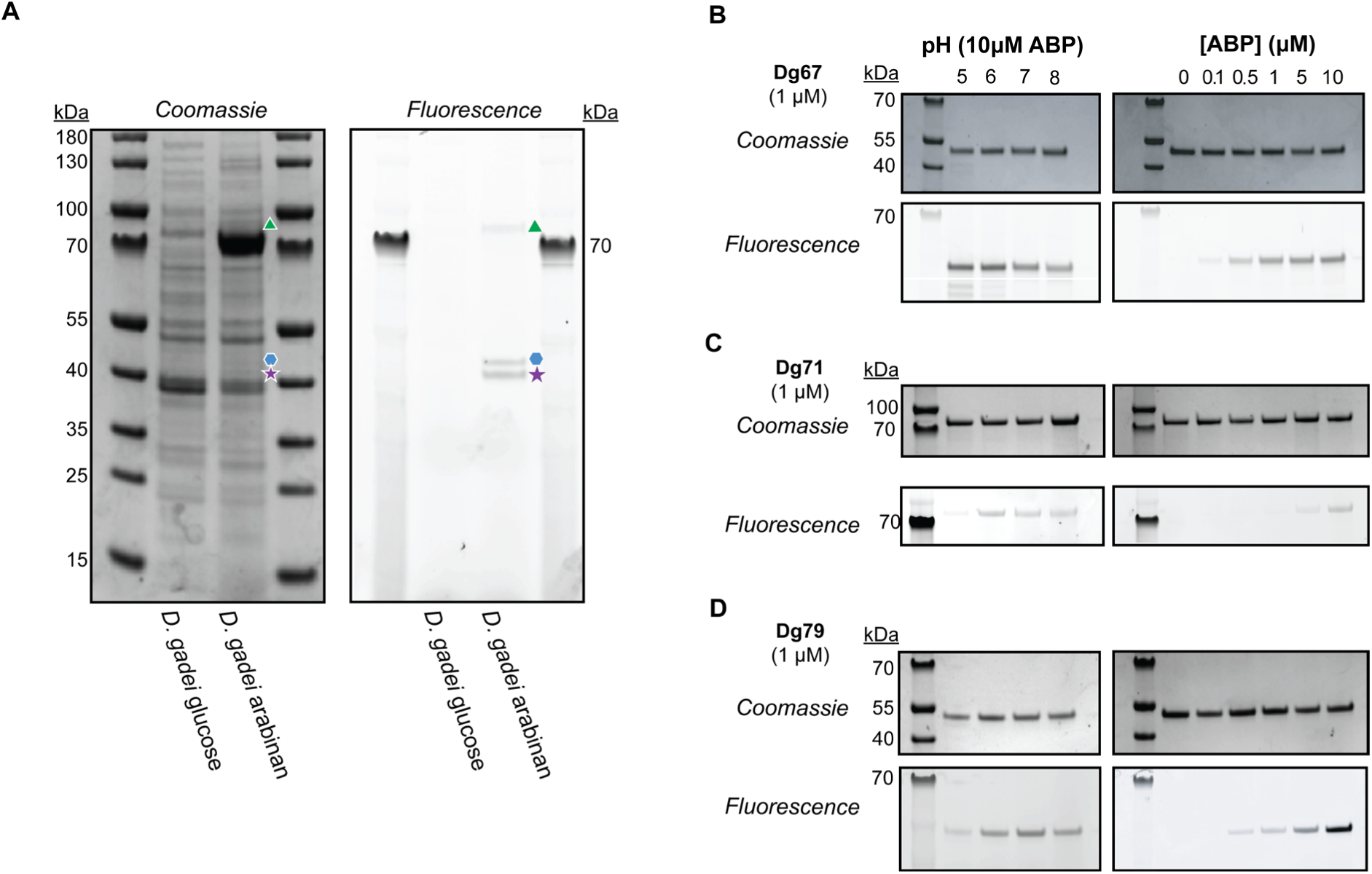
Labelling of *D. gadei* cell lysates and activity profiling of Dg67, Dg71, and Dg79 using probe 5. **A:** *D. gadei* was grown in minimal medium containing either glucose (left lane), or mycobacterial arabinan (right lane). Cell lysates were incubated with **5** (10 μM) and analysed by SDS-PAGE with Coomassie blue or in-gel fluorescence. Bands corresponding to Dg71, Dg79 and Dg67 are indicated by a green triangle, blue hexagon, and purple star, respectively. **B**–**D**: Fluorescent labelling of purified recombinant Dg67 (**B**), Dg71 (**C**) and Dg79 (**D**) with **5** after incubation for 1 h at 37 °C. For pH profiling, reactions contained 1 μM enzyme and 10 μM **5**.

We therefore explored the use of **5** a complementary assay to pNP-α-ᴅ-Ara*f* for rapidly assessing enzyme activity and preferred reaction conditions. This approach is particularly useful for Dg71, which exhibits only weak activity against pNP-α-ᴅ-Ara*f* and requires overnight incubation before a measurable colour change is observed. Recombinant Dg67, Dg71 and Dg79 were incubated with **5**, and the extent of labelling was assessed by intact protein mass spectrometry (**Figure S14**). Unlike aziridine **4**, probe **5** did not completely label any of the three enzymes after 1 h. Approximately 50% of Dg71 remained unlabelled, compared with approximately 20% of Dg67 and Dg79. This difference may reflect variation in the accessibility or architecture of the active sites.

To assess the pH dependence of probe labelling, each GH172 enzyme was incubated with **5** at a 1:10 enzyme-to-probe ratio over the pH range 5–8) for 1 h, followed by SDS-PAGE and in-gel fluorescence analysis (**Figure 6B–D**). Dg67 was labelled across the full pH range, although labelling was less efficient at pH 8. Dg79 was also labelled at all pH values tested, but substantially less efficiently at pH 5. Dg71 showed a similar pH-dependent labelling profile to Dg79, although its overall labelling intensity was lower.

To compare labelling efficiencies, Dg67, Dg71 and Dg79 (1 μM) were incubated at pH 7 with increasing concentrations of **5** (0.1–10 μM). Labelling of Dg67 and Dg79 was detectable at 0.5 μM **5**, corresponding to a 2:1 enzyme-to-probe molar ratio, whereas readily detectable labelling of Dg71 required a tenfold excess of probe. Together with its lower labelling across the pH range tested, these results show that Dg71 reacts more slowly with **5** than Dg67 and Dg79.

We next assessed the effects of **4** and **5** on enzyme thermal stability using nano-differential scanning fluorimetry (nano-DSF). Each enzyme was incubated with **4**, **5** or buffer alone at pH 7 for 1 h at 37 °C, cooled to room temperature and then heated from 20 to 95 °C (**Figures S15** and **S16**; **Table S7**). Among the unmodified enzymes, Dg71 was the most thermostable and Dg79 the least thermostable, with melting temperatures of 70 and 55 °C, respectively. Modification with the aziridine **4** had no measurable effect on the thermal stability of Dg67, but increased the thermal stability of Dg79 by 7 °C. By contrast, modification of Dg71 with **4** decreased its melting temperature by 14 °C (**Figure S15**). Labelling with probe **5** increased the melting temperatures of Dg67 and Dg79 by 10 and 16 °C, respectively, whereas that of Dg71 decreased by 8 °C (**Figure S16**). Thus, covalent probe modification stabilizes Dg67 and Dg79, particularly with **5**, but destabilizes Dg71.

### High-resolution structure of Dg67 with bound inhibitor

Encouraged by the reactivity of **4** and by the well-behaved, high-molecular weight dodecameric assembly of Dg67, which is well suited to accurate pose determination by cryo-EM, we investigated the inhibited complex in detail. Following particle classification and iterative refinement of the map and CTF parameters (**Figure S17**), we obtained a 1.5 Å reconstruction of Dg67 (**Figure 7A**). This map is among the highest-resolution single-particle cryo-EM maps obtained to date. B-factor estimation gave a value of 59 Å^2^, consistent with the high resolution achieved with 1,695,196 particles (**Figure S18)**. The reconstruction reveals a tetrahedrally symmetric complex with an aqueous central cavity. At this resolution, the expected fine structural features were clearly resolved, and the stereochemistry of **4** could be assigned unambiguously. Compound **4** was observed in all 12 active sites of Dg67 (**Figure 7B and Movie S1**). Continuous density was present between the carbon of **4** that mimics the anomeric carbon (C1) and E264, consistent with the formation of a covalent bond (**Figure 7C**). This identifies E264 as the catalytic nucleophile and provides further support for a retaining mechanism in GH172 enzymes. Inspection of the active site showed that the amino group generated by aziridine opening in **4** forms a hydrogen bond with E242, the catalytic acid/base residue of Dg67 (**Figure 7D**). The three Dg67 structures, apo, ᴅ-Araf-bound and in complex with **4**, are highly similar overall, and **4** and ᴅ-Ara*f* occupy closely comparable positions within the active site (**Figure S19A**). The principal difference is observed in the Dg67–**4** map, in which W239 adopts alternative conformations (**Figure S19B** and **Movie S1**). This conformational heterogeneity is not apparent in the apo or ᴅ-Araf-bound structures and may therefore arise from binding of **4**, although it may also be detectable only because of the higher resolution of the Dg67–**4** reconstruction.

**Figure 7:**
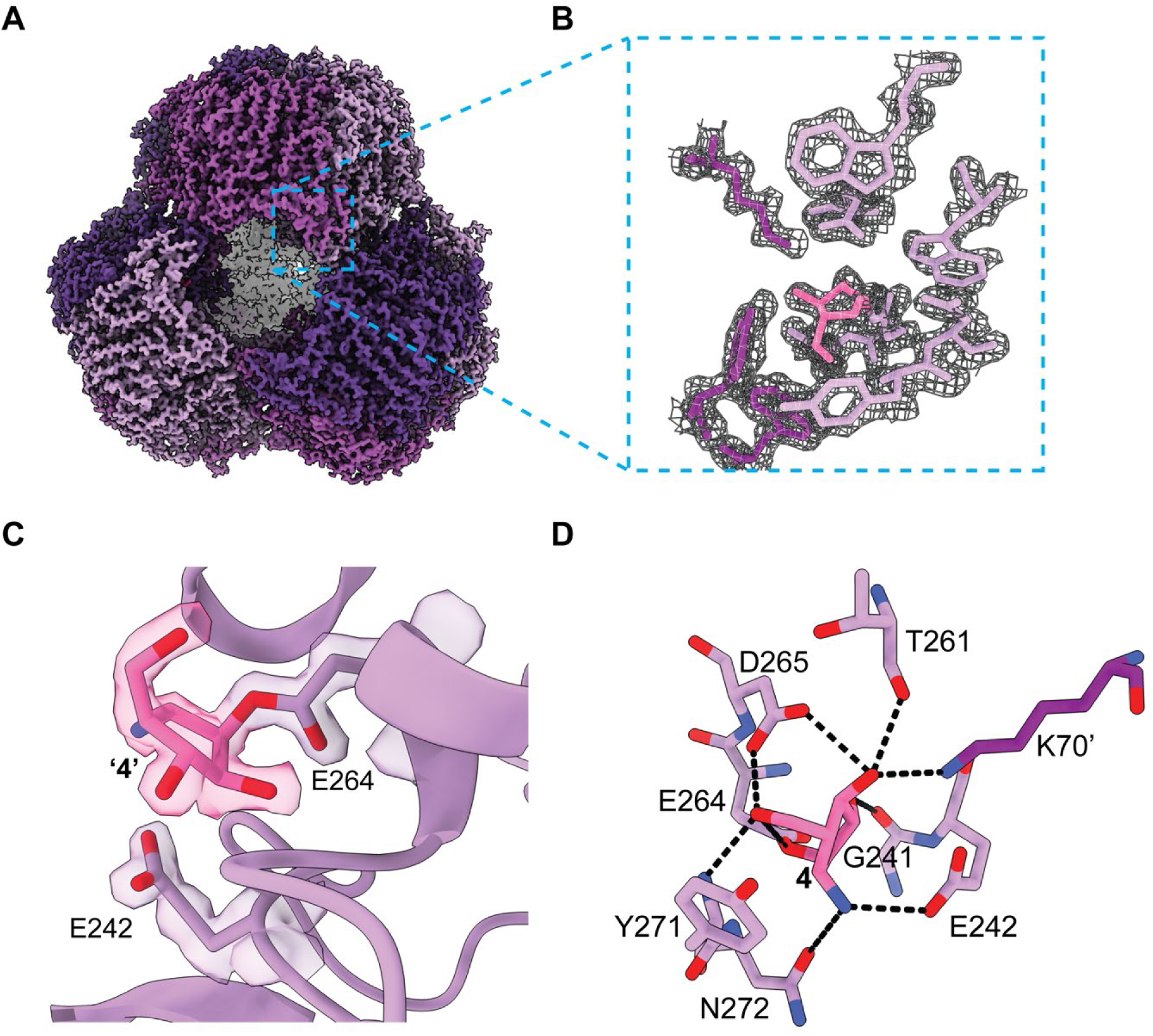
Single-particle cryo-EM reconstruction of Dg67 in complex with 4. **A**: Cryo-EM map of the Dg67-**4** complex at 1.5 Å resolution, with individual protomers shown in different shades of purple. **B**: Mesh representation of the active site map, with density corresponding to **4** shown in pink. **C**: Active site of Dg67 (lilac), with covalently bound **4** (pink). Continuous density between E264 and C1 of **4** is consistent with formation of a covalent bond. The maps in **A–C** are displayed at a threshold level of 0.14. **D**: Active site of the Dg67-**4** complex, with hydrogen-bonding interactions shown as black dashed lines.

### Structural comparison of *D. gadei* GH172 enzymes

Dg67, Dg71 and Dg79 have closely related protomer structures containing a conserved DJR-fold, but assemble into markedly different quaternary structures (**Figures 8** and **S20**). Dg67 forms a dodecamer arranged as a tetramer of trimers, Dg79 forms a dimer of trimers and Dg71 forms a dimer. These structures illustrate the oligomeric diversity possible within family GH172 and show how variation in peripheral loops and C-terminal elements can generate distinct assemblies from a conserved catalytic framework.

**Figure 8:**
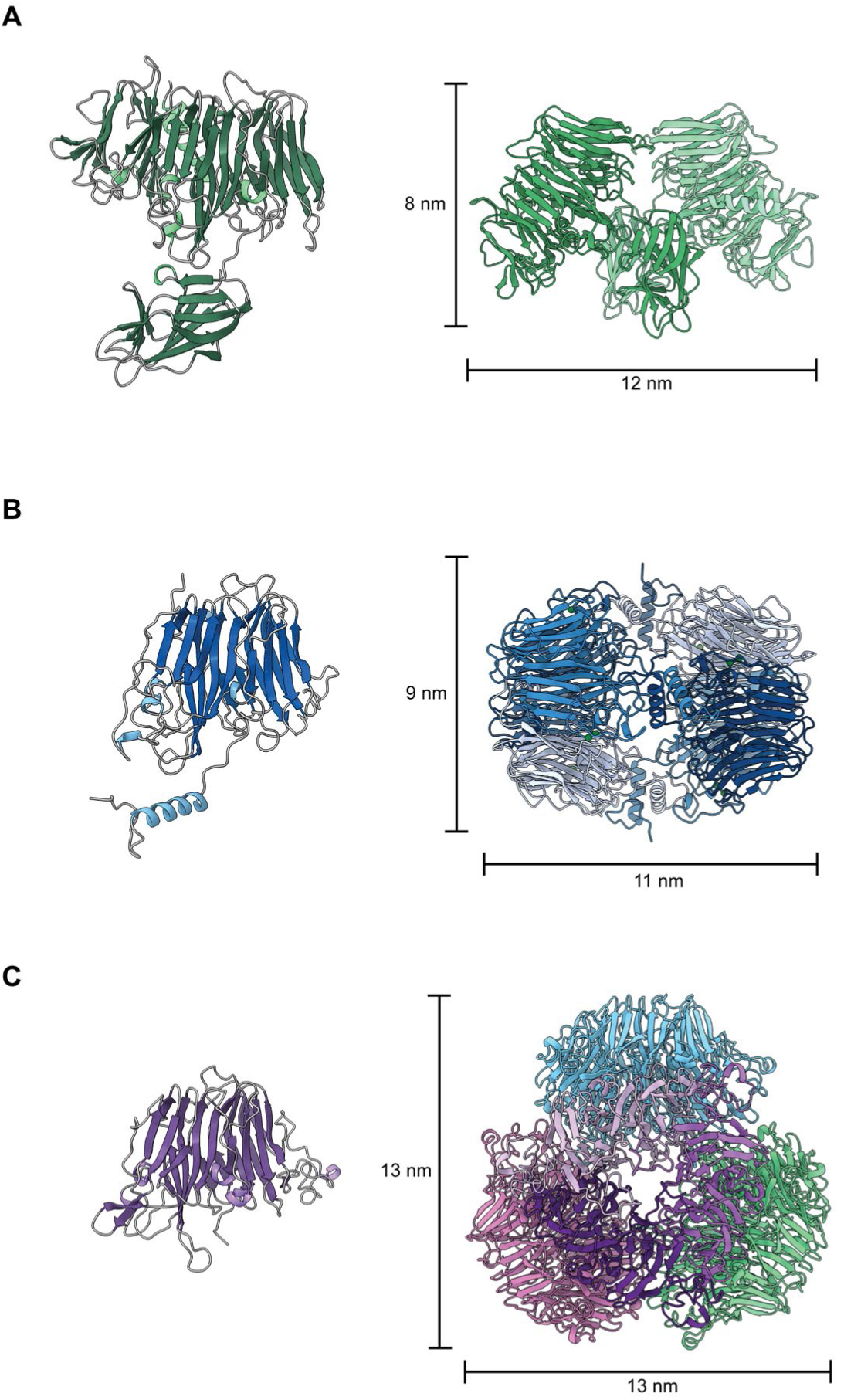
Protomers and quaternary structures of the *D. gadei* GH172 enzymes. **A**: Protomer and dimeric structure of Dg71, shown in shades of green. **B**: Protomer and dimer-of-trimers assembly of Dg79, shown in shades of blues. **C**: Protomer and tetramer-of-trimers assembly of Dg67, with one trimer in shades of purple and the remaining three trimers shown in pink, green and blue for clarity. Scale bars indicate the dimensions of each oligomeric assembly.

Local-resolution analysis of the apo Dg67 and Dg79 cryo-EM reconstructions showed relatively uniform resolution across both assemblies (**Figure S20A/B**), consistent with stable and well-ordered quaternary structures. This interpretation is also consistent with their extensive buried surface areas calculated using PISA analysis (**Table S8**). By contrast, regions surrounding the third β-jelly-roll domain of Dg71 were resolved more poorly than the remainder of the structure (**Figure S20C**), indicating greater local conformational heterogeneity or flexibility. Because this domain contributes residues to the active site, such mobility could influence substrate recognition or access to the active site.

The distinct assemblies are directly related to differences in active-site construction. In Dg67 and Dg79, the active site is formed at the interface between adjacent protomers and therefore depends on oligomerisation, whereas in Dg71 the active site is completed within a single polypeptide (**Figure S21A–C**). The enzymes also differ in monomeric architectures: Dg79 contains an additional C-terminal α-helix, whereas Dg71 contains an additional jelly roll domain and a C-terminal β-sheet domain. The Dg67 and Dg79 protomers superimpose with an RMSD of 0.85 Å across 207 pruned pairs. Dg71 superimposes with Dg67 and Dg79 with RMSDs of 1.03 across 175 pruned pairs, and 1.03 across 170 pruned pairs, respectively (**Figure S21 Di**).

Despite these architectural differences, the catalytic machinery is strongly conserved. Sequence and structural comparisons show that the catalytic residues and several surrounding active site residues are strictly or partially conserved among the three enzymes (**Figure S1** and **Figure S21A-D**). The proposed catalytic nucleophile and acid/base pairs are E264/E242 in Dg67, E406/E387 in Dg71, and E277/E256 in Dg79. The distances between these glutamate residues (5.75 Å in Dg67; 5.76 Å in Dg79; and 6.4 Å in Dg71) are consistent with a retaining catalytic mechanism.^18^ Nevertheless, several non-conserved residues create active-site environments that may help explain the differing substrate preferences of the enzymes (**Figure S21A–D**). The calcium ions observed in Dg79 are absent from Dg67 and Dg71; in Dg67 K70′ occupies the region corresponding to one of these metal-binding sites. Dg67 and Dg79 also contain a tryptophan residue near the edge of the active site (W331 and W212, respectively), although their side chains adopt different orientations in the apo structures. No aromatic residue is present at the equivalent position in Dg71, which may affect its ability to accommodate longer oligosaccharides.

A second aromatic feature is provided by W239 in Dg67 and W253 in Dg79, whereas the corresponding position in Dg71 is occupied by T380. Dg71 compensates for this substitution through Y414, located 34 residues downstream in the sequence, which projects into the same general region and provides aromatic character. Dg71 also contains F53 near the active site, whereas the equivalent region in Dg79 contains K173′. No corresponding side chain projects into this region in Dg67, leaving it with a comparatively open active-site pocket. This more open may contribute to the ability of Dg67 to accommodate branched substrates and cleave both α-1,3 and α-1,5 linkages. Collectively, these local differences are likely to underlie the distinct substrate specificities of the three GH172 enzymes.

## Discussion and Conclusions

This study defines the structures, oligomeric assemblies, and substrate specificities of the three GH172 enzymes from *D. gadei*, Dg67, Dg71 and Dg79. We also report the synthesis and biochemical application of α-ᴅ-arabinofuranosyl cyclophellitol-based inhibitors and activity-based probes for covalent modification and profiling of these enzymes. Finally, the 1.5 Å cryo-EM reconstruction of the Dg67-**4** complex demonstrates the value of high-resolution cryo-EM for mechanistic structural enzymology, allowing the binding and stereochemistry of a covalently bound inhibitor to be unambiguously resolved.

### Activity and specificity of D. gadei GH172 enzymes

All three enzymes reacted more efficiently with the untagged aziridine **4** than with the larger BODIPY-tagged activity-based probe **5**. Because **4** primarily occupies the −1 subsite, whereas the fluorophore and linker of **5** extend beyond this site, this difference may reflect structural interactions within the positive subsites or at the entrance to the active site. This interpretation may be particularly relevant to Dg71, which exhibits weak activity against pNP-α-ᴅ-Ara*f,* synthetic substrates **1-3**, AG, and probe **5**, but is nevertheless efficiently modified by **4**. Together, these observations suggest that Dg71 can recognize the terminal α-ᴅ-Ara*f* residue but has more stringent requirements for the adjoining sugar or glycosidic linkage at the +1 subsite. The preferred substrate of Dg71 therefore remains unidentified. One possibility is that it acts on an arabinose–galactose linkage near the junction between the arabinan and galactan domains of arabinogalactan, while another possibility is that it acts on a succinate modified arabinan such as those present in AG and LAM, although these possibilities will require testing with an appropriately defined substrate.^19^ Overall, Dg67, Dg71 and Dg79 appear to have evolved complementary functions within the same arabinan-degrading locus. Dg67 has the broadest activity, cleaving both α-1,5- and α-1,3-linked Ara*f* residues, whereas Dg79 is selective for α-1,5 linkages. Dg71 displays only limited activity against the substrates examined, consistent with a narrower specificity that is not fully represented by the available compounds. Their collective complementary activities may permit cooperative degradation of the structurally complex arabinan component of mycobacterial arabinogalactan.

### High-resolution structure of the Dg67-**4** complex

The 1.5 Å reconstruction of Dg67 in complex with **4**, together with recent cryo-EM structures of Rubisco bound to transition-state analogue 2-carboxyarabinitol-1,5-bisphosphate at 1.2 Å, β-galactosidase bound to glycosyrin at 1.4 Å, and the NiFe hydrogenase Hucat 1.5 Å, demonstrates that single-particle cryo-EM can achieve resolutions comparable to X-ray crystallography for targets beyond the common cryo-EM test sample apoferritin^20–22^. This resolution also lies within the range reported for crystallographic structures of glycosidases covalently modified by activity-based probes, which have typically been determined at 1.4 to 2.3 Å.^23–32^ The exceptionally high resolution achieved for Dg67 was facilitated by its high molecular weight and tetrahedral symmetry, which allowed accurate particle alignment and averaging during refinement and a large number of asymmetric units to average across. Enzyme discovery and structural biology campaigns have typically favoured compact, soluble targets because of their greater likelihood of crystallisation.^33^ Our results suggest that target selection may instead benefit from a complementary strategy in which smaller proteins are prioritized for crystallography and large, symmetric oligomers are considered both for cryo-EM. Where comparable resolution can be achieved, cryo-EM offers several advantages including substantially lower sample requirements and the avoidance of crystallization. These features are particularly valuable when protein yields are low or small-molecule ligands are available in limited amounts. However, cryo-EM remains dependent on successful specimen preparation and can require substantial computational processing and optimization.

### Development and use of α-ᴅ-arabinofuranosyl cyclophellitol aziridines

Activity-based probes targeting glycosidases have been developed and used for enzymes acting on a range of glycan substrates, including α- and β-mannans, β-glucans, and xylans.^23,34,35^ By contrast, enzymes involved in ᴅ-arabinan degradation have only recently begun to be characterised. GH172 is the only glycoside hydrolase family conclusively associated with α-ᴅ-arabinofuranosidase activity, prompting us to develop the first covalent ᴅ-Ara*f* inhibitors and activity-based probes configured for α-ᴅ-arabinofuranosidases. Although GH172 enzymes are widely distributed across bacterial and fungal species^36^, they are absent from many mycobacterial species, suggesting that additional families of ᴅ-arabinan-degrading enzymes remain to be discovered. Here, we demonstrate the selective labelling of Dg67, Dg71, and Dg79 directly in *D. gadei* cell lysates, establishing the utility of these probes for profiling ᴅ-arabinofuranosidase activity in complex proteomes. These reagents therefore provide a platform for discovering and characterising previously unrecognised ᴅ-arabinan active enzymes, including those involved in arabinan remodelling or degradation in *Mycobacterium tuberculosis* and other pathogenic mycobacteria.

## Author Contributions

Conceptualization: JR, ECL, JNB

Methodology: JR, MH, FK, AS, ECL, JNB

Validation: JR, MH, FK, ECL, JNB

Formal analysis: JR, OA-J, MH, HRB, ECL

Investigation: JR, CM, MH, KJG, MMO, FK

Resources: CM, AB, AS, J-LR, KJG, SH, JT, SJW, HO, KJG

Data curation: JR, EC, JNB

Writing – original draft preparation: JR, ECL, JNB

Writing – review and editing: JR, ECL, JNB, PJM, SJW, HO, HRB, AS

Visualization: JR, ECL, JNB

Supervision: ECL, JNB, PJM, SJW, HO

Funding acquisition: ECL, JNB, PJM, SJW, HO

## Acknowledgements

We thank members of the Newcastle glycobiology group for support and discussions. We thank JMW and GMS for their support. We thank Diamond for access and support of the cryo-EM facilities at the UK national electron Bio-Imaging Centre (eBIC) and our local contact Dr Emma Buzzard. We thank Dr Kate Beckham and Dr Kesha Josts for assistance and use of the Prometheus NT.48. We thank Dr Claudia Schneider for Typhoon FLA use. We thank the entire team at Structura Biotechnology Inc. that designs, develops, maintains and supports the CryoSPARC software system.

## Data availability

Cryo-EM derived maps and models are available from EMDB. Raw micrographs of Dg67 in complex with aziridine **4** are available from EMPIAR. Crystal structure of Dg79 available on PDB.

## Abbreviations

AG: arabinogalactan
GH: glycoside hydrolase
IC-PAD: ion chromatography/pulsed amperometric detection
PUL: polysaccharide utilisation loci
DJR: double β-jelly-roll

