## Supplemental methods and data for "Functional Differentiation of GH172 Arabinofuranosidases Through Divergent Quaternary Structures"

### GH172 Paper Supplementary Information

#### Experimental Methods

*Protein Expression and Purification*

Dg67, Dg71 and Dg79 were produced and purified as previously published[1].

*Production of DFA I*

20 mg/ml of Inulin (Orafti®HP) was incubated with 1µM of IFTase 1 (*Arthrobacter globiformis* P19870) in 50 mM potassium phosphate buffer pH 7.5, at 37 °C for 24 hours. The reaction was then boiled for 10 minutes and stored at -20 °C for further use in assays.

*Production of AG and D-arabinan*

AG was purified from M. smegmatis as done in Al-Jourani, et al.[1] AG was purified into D-arabinan by removing the galactan component using D-galactan degrading enzymes, as previously established.[1]

*Enzymatic Assays*

Reactions were performed with substrates (in water) and enzymes (in 20 mM HEPES pH 7, 150 mM NaCl) with McIlvaine buffer at pH (or the specified pH) as the dominant reaction buffer. Reactions were incubated for 1 or 16 hours (as stated) at 37 °C, and then analysed by IC-PAD, fluorescent gels, or LC-MS.

*Ion chromatographic pulsed amperometric detection (IC-PAD)*

An ICS-6000 system equipped with a CARBOPAC PA-300 anion exchange column (ThermoFisher) was used to analyse the oligosaccharides from enzymatic digestion assays. Detection enabled by PAD using a gold working electrode and a PdH reference electrode with standard Carbo Quad waveform. Buffer A – 100 mM NaOH, Buffer B – 100 mM NaOH, 0.5 M Na Acetate. Each sample was run at a constant flow of 0.25 $ml \cdot min^{-1}$ for 100 min using the following program after injection: 0-10 min; isocratic 100% buffer A, 10-70 min; linear gradient to 60% buffer B, 70-80 min; 100% buffer B. The column was then washed with 10 mins of 500 mM NaOH, then 10 min re-equilibration in 100% buffer A. Data were processed using Chromeleon™ Chromatography Management System and final chromatograms were plotted using GraphPad Prism 10.

*Cryo-EM sample preparation and vitrification*

Samples were prepared for cryo-EM by diluting to suitable concentrations (1 mg/ml for Dg67; 0.7 mg/ml for Dg79; and 0.1 mg/ml for Dg71) and incubating with D-Ara*f* or aziridine **4** (if applicable) for 1 hour at 37 °C. Holey grids (copper. 300 mesh, r 1.2/1.3 from Quantifoil) were glow-discharged using a Pelco glow discharge system. Grids were mounted into a Vitrobot mark IV (Thermofisher) and 2.5 µL of sample was applied and grids were blotted (90% humidity, 8 °C, blot force -5, and blot time 3 seconds) before being plunge-frozen in liquid ethane.

*Cryo-EM data collection*

Grid screening was performed using a Glacios microscope (Thermofisher) equipped with EPU software, University of York. Data collection was performed at the University of York using a Falcon-IV detector for all data sets except Dg67 with aziridine **4** which was collected at eBIC (Diamond Light Source). Information on data collection can be found in **Table S1**.

*Cryo-EM single particle reconstruction*

Processing workflows are shown in supplementary figures (**Figures S4, S6, S8, S9** and **S17**) with 2D-classes, gold-standard FSC curves and final maps displayed. Processing statistics are shown in **Table S2**. Image processing was performed in CryoSPARC 4.7[2]. Movies were imported into CryoSPARC, motion corrected using Patch Motion Correction and then CTF estimated was executed using Patch CTF. Micrographs were then manually curated and analysed by Micrograph Junk Detector. Particles were picked using Blob Picker and particles were curated by rounds of 2D Classification and particle selection. Particles were then subjected to Micrograph Junk Detector to remove those too close to ice and other 'junk'. *Ab initio* models were created without symmetry imposed (C1), followed by Heterogeneous Refinement to remove bad particles before Homogenous Rrefinement. Volumes from Heterogeneous Refinement were analysed in ChimeraX and then Homogeneous Refinement was executed with imposed symmetry (T for Dg67; C2 for Dg71; and D3 for DG79). Masks were created in ChimeraX using the molmap command. Multiple rounds of Homogenous Refinement were performed with per-particle defocus and then Global CTF Refinement. This was followed by Reference Motion Correction and another round of Global CTF Homogenous Refinement. Where applicable, Homogenous Refinement with Eewald sphere correction (EWS) was subsequently executed.

*Cryo-EM single particle high-resolution reconstruction of Dg67-4 complex*

Movies were imported into CryoSPARC, motion-corrected using Patch Motion Correction with a Fourier crop factor of ¾ and then CTF estimation was executed using Patch CTF. Micrographs were analysed by the Micrograph Junk Detector, and outlier micrographs were removed in Manually Curate Exposures, setting the CTF fit resolution (A) to a maximum of 5, Relative Ice Thickness to a maximum of 1.137, and Junk Area (%) maximum of 30. Micrographs were denoised using the Micrograph Denoiser and particles were picked from

428 denoised micrographs using Blob Picker with a particle diameter of 100 – 130 Å. Particles found in junk regions were rejected by using the Micrograph Junk Detector and poor picks were rejected in Inspect Picks with an NCC value of 0.56, and power score of 64-199. Particles were extracted with a box size of 320 pixels, Fourier cropped to 64 pixels and subject to 2D classification with an Initial classification uncertainty factor of 1. Good classes were selected in Select 2D and used for an Ab-Initio Reconstruction job with Window diameter of 0.85-0.99. The selected 2D class averages were used as inputs to Template Picker using all micrographs, with denoising and a particle diameter of 100 Å. Particles found in junk regions were rejected with the Micrograph Junk Detector, and poor picks were rejected in Inspect Picks with an NCC value of 0.31, and power score of 43-172. Particles were extracted with a box size of 320 pixels, Fourier cropped to 64 pixels and subject to 2D classification with an Initial classification uncertainty factor of 1. Particles were then subjected to Homogeneous Refinement using the Ab-Initio initial model, with T symmetry imposed. Particles were re-extracted with a box size of 612 pixels, Fourier cropped to 560 pixels and subsequently underwent Homogenous Refinement with T symmetry, global CTF refinement including Spherical Aberration, Tetrafoil and Anisotropic Magnification, Local CTF refinement with a search range of ± 500, Adaptive Marginalisation, and minimisation over Per-particle scale. A bimodal distribution was observed in the Per-particle scales, so Subset Particles was used with to split the particles into two clusters based on Per-particle scale (optimal) and cluster 1 was carried forwards. XML files were imported using Import Beam Shift, and image beam shift groups for micrographs and associated particles were clustered into 21 groups using Exposure Group Utilities. Homogeneous Refinement was repeated as the previous refinement, except it did not include refinement of Spherical Aberration or Anisotropic Magnification as these are not expected to vary between the exposure groups. Note that no improvement was seen in the resolution with 21 exposure groups compared to using a single group. Reference Motion Correction was run using this input refinement with Fourier crop to box size 600 pixels, and a further Homogeneous Refinement was run with the same setting as the previous refinement except Local CTF refinement used a range of ± 200, and Ewald sphere correction was included. This processing pipeline is shown in **Figure S17** and statistics are in **Tables S5 and S6**.

*Model building and refinement for Dg67 and Dg71*

Initial models for monomers of Dg67 and Dg71 were produced using Alphafold3[3]. These models were refined into cryo-EM maps using Coot[4] and Phenix[5] using real space refine. Model statistics are shown in **Table S3**.

*Structural analysis*

Maps and atomic models were analysed and visualized using ChimeraX v1.10[6]. PDBePISA was used to surface and buried areas of protein structures. [9]

*Crystallography and X-ray data collection*

Crystallization of Dg79 (10 mg/ml) was screened using commercial kits (Molecular Dimensions and Hampton Research) with vapor diffusion sitting drop method. Crystals formed in a buffer containing 0.1 M MOPS/Sodium HEPES pH 7.5, 0.12 M alcohol and 30% EDO_P8K over a period of 2 weeks. Crystals were harvested and flash cooled in liquid nitrogen. X-ray diffraction data were collected at the synchrotron beamline I24t Diamond light source (Didcot, UK) at a temperature of 100 K. The data were integrated using XDS[7] via XIA2 and scaled with Aimless. The space group was confirmed with Pointless[8]. The phase problem was solved by molecular replacement using Phaser[9] and PDB model 4KQ7 as search model from *Bacteroides uniformis*. Subsequently, the structure was auto built with CCP4build on CCP4cloud[10]. The model was improved by rounds of manual building with COOT[4] and refinement with Refmac[11]. The final model was validated with Molprobity[12] and COOT.

*Labelling with aziridines **4** and **5***

Enzyme (1 µL, 10 µM) was mixed with McIlvaine buffer (pH 7, or the indicated pH for pH test experiments; 9 µL; 150 mM). Working stocks of **4** and **5** (typically 100 µM for 10 µM final concentration in assays) were prepared in water from frozen stocks in DMSO (or from freeze dried solid for **4**). 1 µL of the working stock was then added to the enzyme-McIlvaine buffer mixture and incubated at 37 °C for 1 hour with shaking (300 rpm).

*Intact mass LC-MS of recombinant enzymes bound to **4** or **5***

Samples were analysed at 5 µM in 0.1% formic acid. The protein solutions were analysed using Dionex chromatographic system in tandem with a linear ion trap (quadrupole MS) spectrometer ABSciex 6600 system (SCIEX Ontario, Canada). 5 µL of protein solution was injected into the chromatography unit. The protein solution buffer has been exchanged using Fortis C8 column (100´2.1mm, particle size 3 µm) (Fortis Technologies, UK). The mobile phase was 0.1% formic acid in water (buffer A) and 100% acetonitrile (buffer B). The elution gradient (20 min) started at 3% buffer B and increased to 50% buffer B in 16.5 min and completed with 100% buffer B in 18 min. The MS experiment employed electrospray ionisation (ESI) in positive mode covering *m/z* range of 500-2000 *m/z*. The parameters of the MS run were as follows: nebuliser gas 25 psi, turbo gas 10 psi, curtain gas 25 psi, ion spray voltage 4500 V, source temperature 180 °C.

Obtained MS spectra were analysed using UNIDEC software[13].

*Fluorescent gel analysis*

After enzymes were incubated with **5**, SDS-PAGE sample buffer (4x stock, 4 µL) was added. Samples were then centrifuged (13,000 *xg* for 1 min), heated at 95 °C for ten minutes and centrifuged again (13,000 *xg* for 1 min). Samples were loaded entirely into precast gels (NuPAGE Bis-Tris Mini Protein Gels, 4-12%) with prestained protein ladder (5 µL) loaded into an adjacent lane. The gel was ran until the dye front ran off the gel. The gel was then images using a Typhoon FLA 9500 laser scanner and imaged using the Alexa Fluoro 488 channel. The gel was then stained with Quick stain and de-stained in water, prior to be imaged.

*Labelling of D. gadei cell lysate with* ***5***

For *in vivo* enzyme labelling, a wild-type *Dysgonomonas gadei* ATCC BAA-286 glycerol stock was used to inoculate enriched brain-heart infusion broth (37 g/L brain-heart infusion broth, 5 g/L yeast extract) supplemented with 1 µg/ml hematin. The culture was incubated at 37 °C overnight under anaerobic conditions, and 100 µL was used to inoculate Bacteroidota minimal medium with 5 mg/ml glucose as the sole carbon source[1]. After 3 days of anaerobic growth at 37 °C, 100 µL were used to inoculate 3 ml Bacteroidota minimal medium either with 5 mg/ml glucose or 7.5 mg/ml D-arabinan as the sole carbon source, followed by anaerobic cultivation at 37 °C. The equivalent of 0.6 $OD_{600}$ units of cells were collected in microcentrifuge tubes (5,000 xg for 1 min) and the pellets were stored at -20 °C until processing.

*D. gadei* cell pellets were thawed at room temperature and resuspended in lysis buffer (McIlvaine buffer pH 6, 100 mM NaCl, protease inhibitor (Roche cOmplete™, EDTA-free protease inhibitor cocktail), and 25% BugBuster (v/v)). Resuspended pellets were sonicated in an ultrasonic bath (30 seconds on/off, 2 rounds) and mixed in a rotary mixer for 30 minutes at 4 °C. The lysate was clarified by centrifugation at 12000 xg for 20 minutes at 4 °C.

ABP **5** (1 µL, 100 µM) was added to 10 µL of cell lysate and incubated for 1 hour at 37 °C with shaking at 350 rpm. Samples were treated as previously stated for fluoroescent gel analysis.

*Nano-DSF*

Enzyme thermal stability was analysed using a Prometheus NT.48 (NanoTemper) at 30% excitation across a range of 20 to 95 °C. Enzymes were at a concentration of (0.5 mg/ml). Data acquisition was performed using PR.ThermControl (NanoTemper) software and analysed in GraphPad Prism 10.

#### *Chemical Synthesis and Validation*

Compounds **1-3** were synthesized as reported by Han et al. [14], and pNP-Ara*f* as described in Chen et al. [15]

**1-Methoxy-2,6-di-*O*-tosyl-3,4-dihydroxy-α-D-glucopyranoside (MH001)**

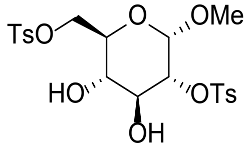

Methyl-α-D-glucopyranoside (1.0 eq., 51.5 mmol, 10.0 g) and $Bu_2SnCl_2$ (0.4 eq., 20.6 mmol, 6.3 g) were added to a stirred solution of $Et_3N$ (2.8 eq., 144 mmol, 20 mL) in dry MeCN (0.1 M, 500 mL). Tosyl chloride (2.4 eq., 124 mmol, 23.6 g) was slowly added and the reaction mixture was stirred at rt under argon atmosphere for 24 h. The reaction mixture was quenched with $H_2O$ (200 mL) and concentrated under reduced pressure. The residue was extracted with EtOAc (3x) and the combined organic layers were washed with $H_2O$, 0.1 M HCl and sat. aq. $NaHCO_3$. The organic phase was dried over $MgSO_4$, filtered and concentrated under reduced pressure. The product was purified by column chromatography (pentane:EtOAc, 9:1 à 1:1) to yield compound **MH001** (25.4 g, 50.5 mmol, 98%). $^1$H NMR (400 MHz, $CDCl_3$) δ 7.84 – 7.72 (m, 4H, $H_{arom,tosyl}$), 7.39 – 7.28 (m, 4H, $H_{aorm,tosyl}$), 4.66 – 4.55 (m, 1H, H-1), 4.28 – 4.23 (m, 2H, H-6a, H-6b), 4.23 – 4.14 (m, 1H, H-2), 3.93 – 3.83 (m, 1H, H-3), 3.72 (dt, *J* = 10.0, 3.6 Hz, 1H, H-5), 3.50 (s, 2H, O*H*-3, O*H*-4), 3.48 – 3.38 (m, 1H, H-4), 3.20 (s, 3H, OC*H*3), 2.43 (d, *J* = 1.2 Hz, 6H, $CH_{3,tosyl}$, $CH_{3,tosyl}$). $^{13}$C NMR (101 MHz, $CDCl_3$) δ 145.1 ($C_{arom}$), 133.2 ($C_{arom}$), 132.7 ($C_{arom}$), 130.0 ($C_{arom}$), 129.9 ($C_{arom}$), 128.1 ($C_{arom}$), 128.1 ($C_{arom}$), 128.0 ($C_{arom}$), 97.2 (C-1), 79.0 (C-2), 71.0 (C-3), 69.8 (C-4), 68.9 (C-5), 68.8 (C-6), 55.9 ($OCH_3$), 55.5 ($CH_{3,tosyl}$). HRMS calculated for $[C_{21}H_{26}O_{10}S_2Na]^+$: 525.0900; found: 525.0859.

**1-Methoxy-2,6-di-*O*-tosyl-3,4-di-*O*-benzoyl-α-D-glucopyranoside (MH009)**

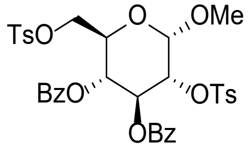

Compound **MH001** (1.0 eq., 50.4 mmol, 25.4 g) was dried by co-evaporation with toluene under argon atmosphere before being dissolved in dry pyridine (0.15 M, 300 mL). Benzoic anhydride (4.0 eq., 202 mmol, 45.6 g) and DMAP (0.3 eq., 15.1 mmol, 1.85 g) were added and the reaction mixture was stirred at rt under argon atmosphere. After 15 h, the reaction mixture was quenched with sat. aq. $NaHCO_3$ and extracted with DCM (3x). The combined organic layers were washed with $H_2O$, sat. aq. $NaHCO_3$ and sat. aq. NaCl. The organic phase was dried over $MgSO_4$, filtered and concentrated under reduced pressure. The product was purified by column chromatography ($CH_2Cl_2$:MeOH, 99.5:0.5) to yield compound **MH009** (35.0 g, 49.3 mmol, 98%). $^1$H NMR (400 MHz, $CDCl_3$) δ 7.81 – 7.66 (m, 4H, $H_{arom}$), 7.65 – 7.54 (m, 4H, $H_{arom}$), 7.54 – 7.43 (m, 2H, $H_{arom}$), 7.40 – 7.26 (m, 9H, $H_{arom}$), 7.25 – 7.17

(m, 2H, $H_{arom}$), 6.98 – 6.90 (m, 2H, $H_{arom}$), 5.82 (t, *J* = 9.7 Hz, 1H, H-3), 5.27 (t, *J* = 9.7 Hz, 1H, H-4), 5.02 (d, *J* = 3.6 Hz, 1H, H-1), 4.52 (dd, *J* = 10.0, 3.6 Hz, 1H, H-2), 4.23 – 4.14 (m, 2H, H-5, H-6a), 4.13 – 4.00 (m, 1H, H-6b), 3.47 (s, 3H, OC$H_3$), 2.35 (s, 3H, C$H_{3,tosyl}$), 2.21 (s, 3H, C$H_{3,tosyl}$). $^{13}$C NMR (101 MHz, $CDCl_3$) δ 165.0 ($C_{arom}$), 164.9 ($C_{arom}$), 145.1 ($C_{arom}$), 145.0 ($C_{arom}$), 133.7 ($C_{arom}$), 133.2 ($C_{arom}$), 132.7 ($C_{arom}$), 132.3 ($C_{arom}$), 130.1 ($C_{arom}$), 129.9 ($C_{arom}$), 129.8 ($C_{arom}$), 129.8 ($C_{arom}$), 128.8 ($C_{arom}$), 128.5 ($C_{arom}$), 128.4 ($C_{arom}$), 128.4 ($C_{arom}$), 128.2 ($C_{arom}$), 128.1 ($C_{arom}$), 127.7 ($C_{arom}$), 124.1 ($C_{arom}$), 97.8 (C-1), 76.4 (C-2), 69.5 (C-3), 68.8 (C-4), 67.8 (C-6), 67.4 (C-5), 56.3 (O$CH_3$), 21.7 ($CH_{3,tosyl}$). HRMS calculated for [C35H34O12S2Na]$^+$: 733.1400; found 733.1383.

**1-Methoxy-2-*O*-tosyl-3,4-di-*O*-benzoyl-6-iodo-α-D-glucopyranoside (MH008)**

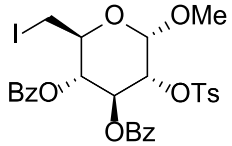

Compound **MH009** (1.0 eq., 88.8 mmol, 63.1 g) and NaI (1.25 eq., 111 mmol, 16.6 g) were dissolved in acetic anhydride (0.5 M, 180 mL) and refluxed under $N_2$ atmosphere for 3 h. The reaction mixture was filtered through Celite which was thoroughly rinsed with $CH_2Cl_2$. The filtrate was washed with aq. sodium thiosulfate, $H_2O$, and sat. aq. $NaHCO_3$. The organic phase was dried over $MgSO_4$, filtered and concentrated *in vacuo*. The product was recrystallized from MeOH to yield compound **MH008** (43.6 g, 65.4 mmol, 74%). $^1$H NMR (400 MHz, $CDCl_3$) δ 7.90 – 7.83 (m, 2H, $H_{arom}$), 7.68 – 7.57 (m, 4H, $H_{arom}$), 7.54 – 7.44 (m, 2H, $H_{arom}$), 7.39 – 7.26 (m, 6H, $H_{arom}$), 6.99 – 6.92 (m, 2H, $H_{arom}$), 5.89 (t, *J* = 9.7 Hz, 1H, H-3), 5.20 (t, *J* = 9.6 Hz, 1H, H-4), 5.09 (d, *J* = 3.7 Hz, 1H, H-1), 4.60 (dd, *J* = 10.1, 3.7 Hz, 1H, H-2), 4.04 – 3.94 (m, 1H, H-5), 3.57 (s, 3H, OC$H_3$), 3.34 (dd, *J* = 11.1, 2.6 Hz, 1H, H-6a), 3.19 (dd, *J* = 11.1, 8.4 Hz, 1H, H-6b), 2.21 (s, 3H, C$H_{3,toysl}$). $^{13}$C NMR (400 MHz, $CDCl_3$) δ 133.7 ($C_{arom}$), 133.1 ($C_{arom}$), 129.9 ($C_{arom}$), 129.8 ($C_{arom}$), 128.5 ($C_{arom}$), 128.2 ($C_{arom}$), 127.7 ($C_{arom}$), 97.8 (C-1), 76.7 (C-2), 72.6 (C-4), 69.1 (C-3), 69.0 (C-5), 56.3 ($CH_3$), 3.6 (C-6). HRMS calculated for [$C_{28}H_{17}O_9SNa$]$^+$: 689.0300; found 689.0312.

**1-Benzyl-4,5-*O*-benzoyl-6-*O*-tosyl-cyclpentaisoxazole (MH066)**

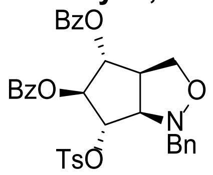

Zinc dust (20 eq., 600 mmol, 39.2 g) was activated with 3 M aq. HCl and subsequently rinsed with $H_2O$ (2x), dioxane (2x) and $Et_2O$ (2x). The activated zinc dust was dried at 50 °C under reduced pressed and argon atmosphere. Compound **MH008** (1.0 eq., 30.0 mmol, 20.0 g) was dissolved in THF/$H_2O$ (9:1). Activated zinc was added and the reaction mixture was sonicated under argon atmosphere. After 7 h, the reaction mixture was filtered over Celite which was thoroughly rinsed with $CH_2Cl_2$. The filtrate was concentrated under reduced pressure. The residue was dissolved in $CH_2Cl_2$ and washed with $H_2O$. The organic phase was dried over $MgSO_4$, filtered and concentrated under reduced pressure. The product was dried by co-evaporation with toluene under $N_2$ atmosphere, before being dissolved in dry toluene (0.2 M, 145 mL). Sodium carbonate (1.5 eq., 43.5 mmol, 4.61 g), *N*-benzyl hydroxylamine hydrochloride (1.5 eq., 43.5 mmol, 6.94 g) were added and the reaction mixture was at 50 °C under $N_2$ atmosphere for 5 h. The reaction mixture was concentrated under reduced pressure, and the residue was dissolved in $CH_2Cl_2$. The product was washed with $H_2O$ and sat. aq. NaCl. The organic phase was dried over $MgSO_4$, filtered and concentrated under reduced pressure. The product was recrystallized from MeOH to yield compound **MH066** (7.58 g, 12.1 mmol, 42% over 2 steps). $^1$H NMR (400 MHz, $CDCl_3$) δ 8.00 – 7.93 (m, 2H, $H_{arom}$), 7.92 – 7.85 (m, 2H, $H_{arom}$), 7.76 – 7.69 (m, 2H, $H_{arom}$), 7.60 – 7.51 (m, 2H, $H_{arom}$), 7.41 (td, *J* = 7.7, 5.5 Hz, 4H, $H_{arom}$), 7.37 – 7.28 (m, 4H, $H_{arom}$), 7.08 (d, *J* = 8.0 Hz, 2H, $H_{arom}$), 5.91 (t, *J* = 8.2 Hz, 1H, H-5), 5.17 (dd, *J* = 8.1, 6.2 Hz, 1H, H-4), 5.09 (dd, *J* = 8.4, 5.5 Hz, 1H, H-6), 4.29 – 4.18 (m, 1H, H-2), 3.95 (d, *J* = 13.5 Hz, 1H, $CH_{2,benzyl}$), 3.91 – 3.80 (m, 2H, $CH_{2,benzyl}$, H-1), 3.33 – 3.19 (m, 1H, H-3), 2.20 (s, 3H, $CH_{3,tosyl}$). $^{13}$C NMR (400 MHz, $CDCl_3$) δ 133.4 ($C_{arom}$), 129.9 ($C_{arom}$), 128.9 ($C_{arom}$), 128.5 ($C_{arom}$), 128.0 ($C_{arom}$), 127.6 ($C_{arom}$), 84.0 (C-2), 80.0 (C-4), 77.1 (C-3), 70.8 (C-6), 70.1 (C-1), 59.4 ($CH_{2,Bn}$), 50.7 (C-5), 21.6 ($CH_3$). HRMS calculated for $[C_{34}H_{31}NO_8SNa]^+$: 636.1700; found 636.1662.

**2,3-*O*-benzoyl-4-hydroxymethyl-azabicyclohexane-5-ol (MH068)**

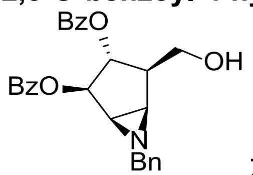

Zinc dust (40 eq., 14 mmol, 0.92 g) was activated with 3 M aq. HCl and subsequently rinsed with $H_2O$ (2x), dioxane (2x) and $Et_2O$ (2x). The activated zinc dust was dried at 50 °C under reduced pressed and argon atmosphere. Isoxazolidine **MH066** (1.0 eq., 0.35 mmol, 0.22 g) was dissolved in AcOH (0.015 M, 23 mL). Activated zinc was added and the mixture was stirred at 35 °C under $N_2$ atmosphere for 2 h. The reaction mixture was filtered over Celite which was thoroughly rinsed with $CH_2Cl_2$. The filtrate was washed with $H_2O$ (2x). The combined aqueous layers were extracted with $CH_2Cl_2$ (3x). The combined organic layers were dried over $MgSO_4$, filtered and concentrated under reduced pressure. The residue was redissolved in MeOH, neutralized with $Et_3N$ until pH≈7, and stirred at rt under $N_2$ atmosphere. After two days, the reaction mixture was concentrated *in vacuo* and redissolved in $CH_2Cl_2$. The mixture was washed with $H_2O$ (2x). The organic phase was dried over $MgSO_4$, filtered and concentrated under reduced pressure. The product was purified by column chromatography (pentane:EtOAc, 70:30 à 50:50) to yield aziridine **MH068** (0.108 g, 0.24

mmol, 70%). $^{1}$H NMR (400 MHz, $CDCl_3$) δ 8.07 – 7.96 (m, 4H, $H_{arom}$), 7.60 – 7.49 (m, 2H, $H_{arom}$), 7.41 (q, *J* = 8.0 Hz, 4H, $H_{arom}$), 7.34 (dd, *J* = 6.6, 3.0 Hz, 2H, $H_{arom}$), 7.13 (dp, *J* = 4.8, 1.9 Hz, 3H, $H_{arom}$), 5.58 (dd, *J* = 6.1, 3.1 Hz, 1H, H-1), 5.45 (dd, *J* = 7.3, 6.1 Hz, 1H, H-6), 3.97 (dd, *J* = 11.1, 5.3 Hz, 1H, H-5a), 3.87 (dd, *J* = 11.1, 6.4 Hz, 1H, H-5b), 3.60 (d, *J* = 13.7 Hz, 1H, $CH_{2,Bn}$), 3.23 (d, *J* = 13.7 Hz, 1H, $CH_{2,Bn}$), 2.65 (dd, *J* = 5.0, 3.1 Hz, 1H, H-2), 2.49 (dd, *J* = 5.0, 3.0 Hz, 1H, H-3), 2.47 – 2.40 (m, 1H, H-4). $^{13}$C NMR (400 MHz, $CDCl_3$) δ 166.6 ($C_{arom}$), 166.5 ($C_{arom}$), 138.6 ($C_{arom}$), 133.4 ($C_{arom}$), 133.3 ($C_{arom}$), 130.0 ($C_{arom}$), 129.9 ($C_{arom}$), 129.7 ($C_{arom}$), 129.7 ($C_{arom}$), 128.5 ($C_{arom}$), 128.4 ($C_{arom}$), 128.4 ($C_{arom}$), 128.1 ($C_{arom}$), 127.6 ($C_{arom}$), 127.2 ($C_{arom}$), 80.0 (C-1), 76.8 (C-6), 62.4 (C-5), 61.3 ($CH_{2,Bn}$), 46.4 (C-4), 43.0 (C-3), 42.6 (C-2). HRMS calculated for $[C27H25NO5Na]^{+}$: 466.1600; found 466.1624.

**2,3-*O*-benzoyl-4-hydroxymethyl-cyclopentene-5-ol (MH072.1)**

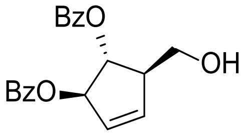

Aziridine **MH068** (1.0 eq., 5.43 mmol, 2.41 g) was dissolved in $CH_2Cl_2$ (0.1 M, 54.3 mL). $NaHCO_3$ (1.5 eq., 8.15 mmol, 0.68 g) and *m*-CPBA (4.0 eq., 21.7 mmol, 3.75 g) were added and the reaction mixture was stirred at 4 °C. After 6 days, the reaction mixture was diluted with $H_2O$ and extracted with $CH_2Cl_2$ (3x). The combined organic layers were dried over $MgSO_4$, filtered and concentrated *in vacuo*. The product was purified by column chromatography (pentane:EtOAc, 95:5 à 60:40) to yield cyclopentene **MH072.1** (1.50 g, 4.41 mmol, 82%). $^{1}$H NMR (400 MHz, $CDCl_3$) δ 8.11 – 8.01 (m, 4H , $H_{arom}$), 7.63 – 7.52 (m, 2H, $H_{arom}$), 7.50 – 7.33 (m, 4H, $H_{arom}$), 6.17 (ddt, *J* = 2.9, 1.9, 0.9 Hz, 1H, H-2), 6.05 (dt, *J* = 5.9, 1.9 Hz, 1H, H-1), 6.03 – 5.98 (m, 1H, H-6), 5.48 (t, *J* = 2.9 Hz, 1H, H-3), 3.90 (dd, *J* = 11.2, 4.3 Hz, 1H, H-5a), 3.78 (dd, *J* = 11.1, 7.5 Hz, 1H, H-5b), 3.01 (dddq, *J* = 6.2, 3.1, 2.1, 1.1 Hz, 1H, H-4). $^{13}$C NMR (400 MHz, $CDCl_3$) δ 167.3 ($C_{arom}$), 166.1 ($C_{arom}$), 135.9 (C-6), 133.7 ($C_{arom}$), 133.6 ($C_{arom}$), 133.4 ($C_{arom}$), 130.3 ($C_{arom}$), 130.2 ($C_{arom}$), 129.9 ($C_{arom}$), 129.8 ($C_{arom}$), 129.7 ($C_{arom}$), 129.5 (C-1), 128.6 ($C_{arom}$), 128.6 ($C_{arom}$), 128.3 ($C_{arom}$), 84.1 (C-2), 81.9 (C-3), 64.3 (C-5), 55.1 (C-4). HRMS calculated for $[C20H18O5Na]^{+}$: 361.1100; found 361.1046.

**2,3-O-benzoyl-5-*O*-(*tert*-butyl-diphenyl-silane)-cyclopentene (MH172)**

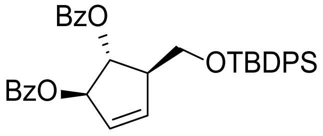

To a solution of alcohol **MH072.1** (1.0 eq., 19.4 mmol, 6.58 g) in DMF (0.3 M, 65 mL) was added imidazole (2.0 eq., 38.8 mmol, 2.56 g) and TBDPS-Cl (1.2 eq., 23.3 mmol, 6.05 mL). The reaction mixture was stirred at rt for 17 h, after which the reaction mixture was diluted with EtOAc and washed with $H_2O$ (3x). The combined aqueous layers were extracted with EtOAc (2x). The combined extracted layers were dried over $MgSO_4$, filtered and concentrated under reduced pressure. The product was purified by column chromatography (pentane:$Et_2O$, 98:2 à 96:4) to yield compound **MH172** (11.6 g, 19.9 mmol, quant.). $^1$H NMR (400 MHz, $CDCl_3$) δ 8.03 (ddd, *J* = 11.9, 8.3, 1.3 Hz, 4H, $H_{arom}$), 7.76 – 7.66 (m, 6H, $H_{arom}$), 7.59 – 7.47 (m, 2H, $H_{arom}$), 7.40 – 7.28 (m, 8H, $H_{arom}$), 6.08 (ddt, *J* = 3.0, 2.1, 1.0 Hz, 1H, H-1), 6.05 (ddd, *J* = 6.0, 2.1, 1.1 Hz, 1H, H-2), 5.98 (dt, *J* = 6.0, 2.0 Hz, 1H, H-3), 5.71 (t, *J* = 3.3 Hz, 1H, H-6), 3.97 – 3.83 (m, 2H, H-5), 3.04 (tdd, *J* = 7.8, 4.2, 2.2 Hz, 1H, H-4), 1.06 (d, *J* = 4.5 Hz, 9H, $H_{t\text{-butyl}}$). $^{13}$C NMR (400 MHz, $CDCl_3$) δ 166.3 ($C_{arom}$), 166.1 ($C_{arom}$), 137.0 ($C_{arom}$), 135.8 ($C_{arom}$), 135.4 ($C_{arom}$), 134.9 ($C_{arom}$), 133.6 ($C_{arom}$), 133.5 ($C_{arom}$), 133.2 ($C_{arom}$), 133.1 ($C_{arom}$), 130.1 ($C_{arom}$), 129.9 ($C_{arom}$), 129.9 ($C_{arom}$), 129.8 ($C_{arom}$), 129.8 ($C_{arom}$), 129.7 ($C_{arom}$), 129.1 ($C_{arom}$), 128.5 ($C_{arom}$), 128.4 ($C_{arom}$), 127.8 ($C_{arom}$), 84.7 (C-1), 80.4 (C-6), 64.7 (C-5), 53.9 (C-4), 26.9 ($C_{t\text{-butyl}}$), 26.7 ($C_{t\text{-butyl}}$). HRMS calculated for $[C_{36}H_{36}O_6SiNa]^+$: 615.2200; found 615.2173.

**5-*O*-(*tert*-butyl-diphenyl-silane)-cyclopentene-2,3-diol (MH174)**

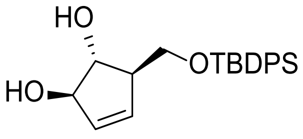

To a solution of compound **MH172** (1.0 eq., 7.0 mmol, 4.04 g) in MeOH (0.3 M, 23 mL) was added NaOMe (25 wt% in MeOH) (0.3 eq., 2.1 mmol, 0.48mL) and the reaction mixture was stirred at rt for 3.5 h. The reaction was neutralized to pH 7 with AcOH and concentrated *in vacuo*. The product was purified by column chromatography ($CH_2Cl_2$:MeOH, 99:1 à 98:2) to yield diol **MH174** (2.32 g, 6.29 mmol, 90%). $^1$H NMR (400 MHz, $CDCl_3$) δ 7.65 (ddt, *J* = 6.5, 2.7, 1.5 Hz, 4H, $H_{arom}$), 7.50 – 7.33 (m, 6H, $H_{arom}$), 5.87 (dt, *J* = 6.0, 2.1 Hz, 1H, H-1), 5.69 (d, *J* = 6.1 Hz, 1H, H-6), 4.59 (s, 1H, H-2), 4.10 (t, *J* = 3.7 Hz, 1H, H-3), 3.84 (dd, *J* = 10.0, 4.4 Hz, 1H, H-$5_a$), 3.63 (dd, *J* = 10.0, 6.2 Hz, 1H, H-$5_b$), 2.71 (s, 1H, H-4), 1.05 (s, 9H, $H_{t\text{-butyl}}$). NMR (400 MHz, $CDCl_3$) δ 135.7 ($C_{arom}$), 135.6 ($C_{arom}$), 133.4 (C-1), 132.5 (C-6), 130.0 ($C_{arom}$), 127.9 ($C_{arom}$), 84.0 (C-3), 83.0 (C-2), 65.2 (C-1), 54.4 (C-5), 26.9 ($C_{t\text{-butyl}}$). HRMS calculated for $[C_{22}H_{28}O_3SiNa]^+$: 391.1700; found 391.1699.

**5- *O*-(*tert*-butyl-diphenyl-silane)-Arabinofuranose-β-epoxide (RvB15)**

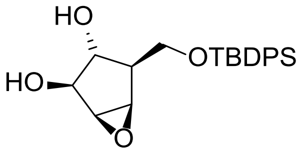

*m*-CPBA (4 eq., 10.34 mmol, 1.78 g) was added to a solution of compound **MH174** (1.0 eq., 2.59 mmol, 0.95 g) in DCM (0.1 M, 26 mL) and the mixture was stirred for two days at 4 °C. The mixture was washed with $H_2O$. The organic layer was dried over $MgSO_4$, filtered and concentrated *in vacuo*. The crude product was purified by silica column chromatography (DCM:MeOH, 99.5:0.5 à 49:1) yielding compound **RvB15** (0.9281 g,

2.41 mmol, 93% yield) as a clear oil. $^1$H NMR (400 MHz, $CDCl_3$) δ 7.70 (dt, *J* = 6.5, 1.7 Hz, 4H, $H_{arom}$), 7.51 – 7.37 (m, 6H, $H_{arom}$), 4.12 (dd, *J* = 5.9, 1.4 Hz, 1H, H-1), 3.93 (d, *J* = 7.6 Hz, 2H, H-5), 3.55 (dd, *J* = 3.1, 1.4 Hz, 1H, H-2), 3.53 – 3.47 (m, 2H, H-3, H-6), 2.24 (td, *J* = 7.5, 1.3 Hz, 1H, H-4), 1.09 (s, 9H, $CH_{3,t\text{-butyl}}$). $^{13}$C NMR (400 MHz, $CDCl_3$) δ 135.6 ($C_{arom}$), 130.0 ($C_{arom}$), 127.9 ($C_{arom}$), 79.5 (C-1), 77.4 (C-6), 63.7 (C-5), 56.4 (C-2), 54.2 (C-3), 47.8 (C-4), 26.9 ($CH_{3,t\text{-butyl}}$). HRMS calculated for $[C_{22}H_{28}O_4SiNa]^+$: 407.1700; found 407.1649.

**2,3-*O*-Benzoyl-5-*O*-(*tert*-butyl-diphenyl-silane)-arabinofuranose-β-epoxide (RvB16)**

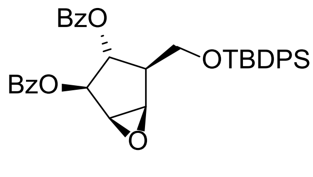

Compound **RvB15** (1.0 eq., 2.41 mmol, 0.9281 g) was dissolved in dry pyridine (0.15 M, 16 mL) under $N_2$ atmosphere. $Bz_2O$ (4 eq., 9.64 mmol, 2.18 g) and DMAP (0.3 eq., 0.723 mmol, 0.088 g) were added and the reaction mixture was stirred overnight at rt. The mixture was quenched with sat. aq. $NaHCO_3$ and extracted with EtOAc. The organic layer was washed with $H_2O$, sat. aq. $NaHCO_3$ and brine. The organic layer was dried over $MgSO_4$, filtered and concentrated *in vacuo*. Purification of the crude product by silica column chromatography (pentane:$Et_2O$, isocratic 90:10) yielded compound **RvB16** (1.3177 g, 2.22 mmol, 93%). $^1$H NMR (400 MHz, $CDCl_3$) δ 8.16 – 8.04 (m, 2H, $H_{arom}$), 7.99 – 7.91 (m, 2H, $H_{arom}$), 7.68 (ddd, *J* = 8.0, 4.2, 1.7 Hz, 4H, $H_{arom}$), 7.62 – 7.55 (m, 2H, $H_{arom}$), 7.55 – 7.29 (m, 11H, $H_{arom}$), 5.61 (dd, *J* = 5.9, 1.5 Hz, 1H, H-2), 5.22 (dd, *J* = 7.4, 5.9 Hz, 1H, H-3), 4.10 – 3.97 (m, 2H, H-5), 3.95 (dd, *J* = 3.1, 1.5 Hz, 1H, H-1), 3.88 (dd, *J* = 3.1, 1.4 Hz, 1H, H-6), 2.58 (dddd, *J* = 9.2, 7.1, 5.4, 1.4 Hz, 1H, H-4), 1.07 (s, 9H, $CH_{3,t\text{-butyl}}$). $^{13}$C NMR (400 MHz, $CDCl_3$) δ 135.6 ($C_{arom}$), 135.6 ($C_{arom}$), 133.4 ($C_{arom}$), 133.2 ($C_{arom}$), 130.0 ($C_{arom}$), 129.9 ($C_{arom}$), 129.8 ($C_{arom}$), 129.8 ($C_{arom}$), 128.5 ($C_{arom}$), 128.4 ($C_{arom}$), 127.9 ($C_{arom}$), 127.8 ($C_{arom}$), 79.6 (C-2), 74.2 (C-3), 62.2 (C-5), 55.1 (C-1), 54.78 (C-6), 46.7 (C-4), 26.9 ($CH_{3,t\text{-butyl}}$ ). HRMS calculated for $[C_{36}H_{36}O_6SiNa]^+$: 615.2200; found 615.2173

**1-azido-2,3-*O*-benzoyl-5-*O*-(*tert*-butyl-diphenyl-silane)-cyclohexan-6-ol and 2,3-*O*-benzoyl-5-OTBDPS-6-azido-cyclohexan-1-ol (RvB17 & RvB18)**

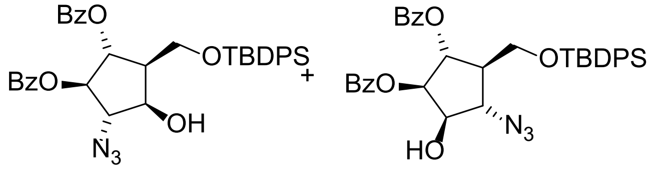

Compound **RvB16** (1.0 eq., 1.92 mmol, 1.3177 g) was dried by co-evaporation with toluene prior to solvation in dry DCM (0.3 M, 6.4 mL). Trimethylsilyl azide (1.5 eq, 2.88 mmol, 0.38 mL) and $BF_3{\cdot}OEt_2$ (2 eq., 3.84 mmol, 1.15 mL) were added and the reaction mixture was stirred overnight at rt under argon atmosphere. The mixture was washed with $H_2O$ and brine. The organic phase was dried over $MgSO_4$, filtered and concentrated *in vacuo*. The crude product was purified by silica column chromatography (pentane:EtOAc, 95:5 ⇒ 90:10) yielding a mixture of compound **RvB17** and **RvB18** (0.7005 g, 1.10 mmol, 57% as a yellowish oil. $^{1}$H NMR (400 MHz, $CDCl_3$) δ 8.17 – 8.09 (m, 1H, $H_{arom}$), 8.08 (d, *J* = 1.5 Hz, 1H, $H_{arom}$), 8.06 – 7.94 (m, 2H, $H_{arom}$), 7.76 – 7.68 (m, 4H, $H_{arom}$), 7.68 – 7.55 (m, 3H, $H_{arom}$), 7.49 (t, *J* = 6.7 Hz, 2H, $H_{arom}$), 7.48 – 7.40 (m, 6H, $H_{arom}$), 7.40 – 7.25 (m, 5H, $H_{arom}$), 5.72 (dd, *J* = 7.0, 3.1 Hz, 1H, H-3), 5.49 (dd, *J* = 6.0, 3.1 Hz, 1H, H-2), 4.55 – 4.42 (m, 1H H-1), 4.13 – 4.08 (m, 1H H-6), 3.97 – 3.89 (m, 2H, H-5), 2.56 (d, *J* = 5.2 Hz, 1H, O*H*), 2.25 – 2.13 (m, 1H H-4), 1.11 (d, *J* = 9.8 Hz, 9H, $CH_{3,t\text{-butyl}}$). $^{13}$C NMR (400 MHz, $CDCl_3$) δ 135.7 ($C_{arom}$), 135.6 ($C_{arom}$), 133.6 ($C_{arom}$), 133.4 ($C_{arom}$), 130.1 ($C_{arom}$), 130.0 ($C_{arom}$), 130.0 ($C_{arom}$), 130.0 ($C_{arom}$), 129.9 ($C_{arom}$), 129.8 ($C_{arom}$), 128.6 ($C_{arom}$), 128.6 ($C_{arom}$), 128.5 ($C_{arom}$), 127.9 ($C_{arom}$), 127.9 ($C_{arom}$), 77.2 (C-2), 75.7 (C-1), 73.6 (C-3), 64.8 (C-6), 60.5 (C-5), 48.4 (C-4), 26.9 ($CH_{3,t\text{-butyl}}$). HRMS calculated for $[C_{36}H_{37}N_3O_6SiNa]^+$: 658.2400; found 658.2343.

**1-azido-2,3-*O*-benzoyl-5-*O*-(*tert*-butyl-diphenyl-silane)-6-*O*-mesyl-cyclohexane and 1-*O*-Mesyl-2,3-*O*-benzoyl-5-*O*-(*tert*-butyl-diphenyl-silane)-6-azido-cyclohexane (RvB19 & RvB20)**

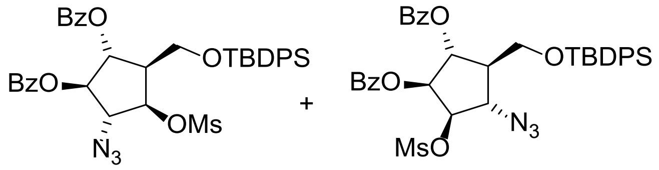

A mixture of compounds **RvB17** and **RvB18** (1.0 eq., 1.1 mmol, 0.70 g) was co-evaporated with toluene and then dissolved in dry DCM (3.66 mL, 3 M). Dry TEA (2 eq., 2.2 mmol, 0.31 mL) and DMAP (0.3 eq., 0.33 mmol, 0.04 g) were added to the reaction mixture. Methanesulfonyl chloride (2 eq., 2.2 mmol, 0.17 mL) was slowly added and the mixture was stirred overnight at rt under argon atmosphere. The reaction was quenched with $H_2O$, extracted with DCM and the organic layer was washed with $H_2O$. The organic layer was dried over $MgSO_4$, filtered and concentrated under reduced pressure. The crude product was purified using silica column chromatography (pentane:EtOAc, isocratic 80:20) yielding a mixture of **RvB19** and **RvB20** (0.6143 g, 0.86 mmol, 78%) as a clear oil. $^{1}$H NMR (400 MHz, $CDCl_3$) δ 8.11 – 7.96 (m, 4H, $H_{arom}$), 7.74 – 7.57 (m, 6H, $H_{arom}$), 7.55 – 7.30 (m, 11H, $H_{arom}$), 5.70 – 5.59 (m, 2H, H-2, H-3), 5.28 – 5.10 (m, 1H, H-1), 4.45 – 4.39 (m, 1H, H-6), 3.94 (t, *J* = 4.0 Hz, 2H, C5**H2**), 3.10 (d, *J*= 22.2 Hz, 3H, $CH_{3,mesyl}$), 2.28 – 2.17 (m, 1H, H-4), 1.11 (d, *J* = 9.3 Hz, 9H, $CH_{3,t\text{-butyl}}$). $^{13}$C NMR (400 MHz, $CDCl_3$) δ 135.7 ($C_{arom}$), 135.5 ($C_{arom}$), 133.9 ($C_{arom}$), 133.8 ($C_{arom}$), 133.6 ($C_{arom}$), 130.0 ($C_{arom}$), 130.0 ($C_{arom}$), 129.9 ($C_{arom}$), 128.7 ($C_{arom}$), 128.6 ($C_{arom}$), 128.0 ($C_{arom}$), 127.9 ($C_{arom}$), 80.4 (C-1), 74.7 (C-2), 74.2 (C-3), 63.4 (C-6), 60.3 (C-5), 48.3 (C-4), 38.6 ($CH_{3,mesyl}$), 26.9 ($CH_{3,t\text{-butyl}}$). HRMS calculated for $[C_{37}H_{39}N_3O_8SSiNa]^+$: 736.2100; found 736.2119.

**2,3-*O*-benzoyl-5-*O*-(*tert*-butyl-diphenyl-silane)-arabinofuranose-α-aziridine (RvB21)**

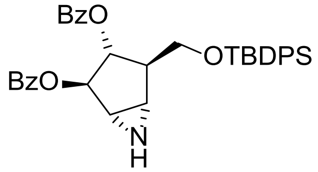

The mixture of **RvB19** and **RvB20** (1.0 eq., 0.86 mmol, 0.61 g) and triphenyl phosphate (3.5 eq., 3.01 mmol, 0.79 g) were suspended in THF/$H_2O$ (4:1, 0.04 M, 21.5 mL). DIPEA (2.5 eq., 2.15 mmol, 0.37 mL) was added and the mixture was stirred for 1h at 60 °C. The THF was evaporated and the mixture was diluted EtOAc. The mixture was washed with $H_2O$, and the aqueous layer was extracted with EtOAc (2x). The combined organic layers were dried with $MgSO_4$, filtered and concentrated in vacuo. Purification of the crude product via silica column chromatography (pentane/EtOAc 90:10 à 70:30) yielded compound **RvB21** (0.3715 g, 0.6277 mmol, 73%) as a clear oil. $^1$H NMR (400 MHz, $CDCl_3$) δ 8.04 (d, $J$ = 7.7 Hz, 2H, $H_{arom}$), 7.96 – 7.86 (m, 2H, $H_{arom}$), 7.77 – 7.69 (m, 4H, $H_{arom}$), 7.66 – 7.54 (m, 2H, $H_{arom}$), 7.53 – 7.45 (m, 2H, $H_{arom}$), 7.45 – 7.32 (m, 8H, $H_{arom}$), 5.48 (s, 1H, H-2), 5.29 (s, 1H, H-3), 4.03 (dd, $J$ = 10.0, 6.9 Hz, 1H H-5), 3.87 (t, $J$ = 9.7 Hz, 1H, H-5), 3.10 – 2.62 (m, 3H H-1,H-4, H-6), 1.11 (s, 9H, $CH_{3,t\text{-butyl}}$). $^{13}$C NMR (400 MHz, $CDCl_3$) δ 135.7 ($C_{arom}$), 135.6 ($C_{arom}$), 133.4 ($C_{arom}$), 129.9 ($C_{arom}$), 129.8 ($C_{arom}$), 129.8 ($C_{arom}$), 128.6 ($C_{arom}$), 128.5 ($C_{arom}$), 127.9 ($C_{arom}$), 127.8 ($C_{arom}$), 79.5 (C-3), 78.9 (C-2), 63.3 (C-5), 50.5 (C-4), 39.2 (C-6), 38.1 (C-1), 26.9 ($CH_{3,t\text{-butyl}}$). HRMS calculated for $[C_{36}H_{37}NO_5SiNa]^+$: 614.2300; found 614.2333.

**5-*O*-(*tert*-butyl-diphenyl-silane)-arabinofuranose-α-aziridine (MH367)**

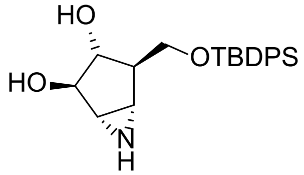

Compound **RvB21** (1.0 eq., 0.628 mmol, 0.372 g) was dissolved in MeOH after which NaOMe (25 wt% in MeOH, 0.3 eq., 0.188 mmol, 0.11 mL) was added. After stirring overnight, the reaction mixture was neutralized with AcOH and concentrated *in vacuo*. The product was purified via column chromatography (DCM:MeOH, 100:0 à 96:4) to yield compound **MH367** (0.077 g, 0.20 mmol, 32%). $^1$H NMR (400 MHz, $CDCl_3$) δ 7.75 – 7.61 (m, 4H, $H_{arom}$), 7.48 – 7.35 (m, 6H, $H_{arom}$), 3.97 (s, 1H, H-2), 3.86 – 3.78 (m, 2H, H-3, H-5), 3.74 (dd, $J$ = 10.7, 3.7 Hz, 1H, H-5), 2.89 (dd, $J$ = 3.8, 1.6 Hz, 1H, H-1), 2.63 (dd, $J$ = 3.9, 1.2 Hz, 1H, H-6), 2.18 (t, $J$ = 3.5 Hz, 1H, H-4), 1.07 (s, 9H, $H_{t\text{-butyl}}$). $^{13}$C NMR (400 MHz, $CDCl_3$) δ 135.9 ($C_{arom}$), 135.7 ($C_{arom}$), 130.3 ($C_{arom}$), 130.3 ($C_{arom}$), 128.1($C_{arom}$), 81.9 (C-3), 76.8 (C-2), 64.8 (C-5), 50.8 (C-4), 39.9 (C-1), 36.5 (C-6), 26.9 ($C_{t\text{-butyl}}$). HRMS calculated for $[C_{22}H_{29}NO_3SiNa]^+$: 406.1809, found: 406.1809

**5-*O*-(*tert*-butyl-diphenyl-silane)-arabinofuranose-α-6-azidohexyl-aziridine (MH371)**

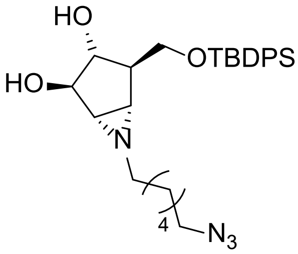

Compound **MH367** (1.0 eq., 21.1 µmol, 8.1 mg) was dried by co-evaporation with toluene under $N_2$ atmosphere prior to solvation in dry THF (0.1 M, 211 µL). DIPEA (1.1 eq., 23.2 µmol, 4.0 µL) and 6-azidohexyl trifluoromethanesulfonate (1.1 eq., 23.2 µmol, 6.39 mg) were added at 0 °C and the reaction mixture was stirred at 0 °C under $N_2$ atmosphere. After 22h LCMS confirmed complete consumption of the starting material. The reaction mixture was diluted with $H_2O$ and extracted with DCM (3x). The organic phase was washed with $H_2O$ and brine, dried over $MgSO_4$, filtered and concentrated under reduced pressure. The product was purified by column chromatography (pet. ether:EtOAc, 70:30) to yield compound **MH371** (6.9 mg, 13.6 µmol, 64%). $^1$H NMR (400 MHz, $CDCl_3$) δ 7.72 – 7.66 (m, 2H, $H_{arom}$), 7.66 – 7.60 (m, 2H, $H_{arom}$), 7.52 – 7.37 (m, 6H, $H_{arom}$), 3.91 (s, 2H, H-2, O*H*), 3.79 (dt, *J* = 6.9, 3.4 Hz, 3H, H-3, H-5, O*H*), 3.71 (dd, *J* = 10.8, 3.7 Hz, 1H, H-5), 3.26 (t, *J* = 6.9 Hz, 2H, $CH_{2,alkyl}$), 2.32 (d, *J* = 4.0 Hz, 1H, H-1), 2.25 (td, *J* = 6.8, 4.7 Hz, 2H, $CH_{2,alkyl}$), 2.12 (t, *J* = 3.6 Hz, 1H, H-4), 2.03 (d, *J* = 4.0 Hz, 1H, H-6), 1.61 (qd, *J* = 6.7, 3.5 Hz, 4H, $CH_{2,alkyl}$), 1.50 (dd, *J* = 8.2, 5.4 Hz, 1H, $CH_{2,alkyl}$), 1.44 – 1.32 (m, 5H, $CH_{2,alkyl}$), 1.07 (s, 9H, $H_{t\text{-butyl}}$). $^{13}$C NMR (400 MHz, $CDCl_3$) δ 135.9 ($C_{arom}$), 135.7 ($C_{arom}$), 130.3 ($C_{arom}$), 130.2 ($C_{arom}$), 128.1 ($C_{arom}$), 82.2 (C-3), 76.4 (C-2), 64.6 (C-5), 57.5 ($CH_{2,alkyl}$), 51.5 ($CH_{2,alkyl}$), 50.8 (C-4), 48.2 (C-1), 44.7 (C-6), 29.7 ($CH_{2,alkyl}$), 28.9 ($CH_{2,alkyl}$), 27.0 ($CH_{2,alkyl}$), 26.9 ($C_{t,butyl}$), 26.8 ($CH_{2,alkyl}$), 19.2 ($C_{t,butyl}$). HRMS calculated for $[C_{28}H_{40}N_4O_3SiNa]^+$: 531.2762, found: 531.2762

**Arabinofuranose-α-6-azidohexyl-aziridine (MH373)**

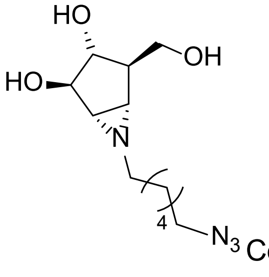

Compound **MH371** (1.0 eq., 13.6 µmol, 6.9 mg) was dried by co-evaporation with toluene under $N_2$ atmosphere and dissolved in THF (0.1 M, 136 µL). To the reaction mixture was added TBAF (1.0 M in THF, 1.5 eq., 20.4 µmol, 20.4 µL) and AcOH (1.0 eq., 13.6 µmol, 0.77 µL) and it was stirred at rt under $N_2$ atmosphere. After 17 h LCMS confirmed complete consumption of the starting material, after which the reaction was concentrated under reduced pressure. The product was purified via column chromatography (DCM:MeOH, 100:0 à 96:4) to yield compound **MH373** (3.6 mg, 13.6 µmol, quant.). $^1$H NMR (400 MHz, MeOD) δ 3.84 (d, *J* = 0.6 Hz, 1H, H-3), 3.67 – 3.54 (m, 3H, H-2, H-5), 3.31 (dt, *J* =

3.3, 1.7 Hz, 12H, C$H_{2,alkyl}$, MeOD), 2.32 – 2.26 (m, 3H, H-1, C$H_{2,alkyl}$), 2.24 (dd, $J$ = 4.0, 1.7 Hz, 1H, H-6), 2.18 – 2.08 (m, 1H, H-4), 1.64 – 1.50 (m, 5H, C$H_{2,alkyl}$), 1.41 (p, $J$ = 3.8 Hz, 4H, C$H_{2,alkyl}$). $^{13}$C NMR (400 MHz, MeOD) δ 81.6 (C-2), 77.6 (C-3), 62.4 (C-5), 58.4 ($C$H$_{2,alkyl}$), 52.5 (C-4) , 52.4 ($C$H$_{2,alkyl}$), 48.4 (C-1), 46.3 (C-6), 30.6 ($C$H$_{2,alkyl}$), 29.9 ($C$H$_{2,alkyl}$), 28.0 ($C$H$_{2,alkyl}$), 27.7 ($C$H$_{2,alkyl}$). HRMS calculated for $[C_{12}H_{22}N_4O_3Na]^+$: 293.1584, found: 293.1584

**Ara*f* aziridine, BODIPY (MH374) aka 5**

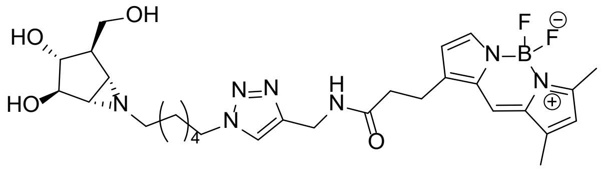

Compound **MH373** (1.0 eq., 13.6 μmol, 3.6 mg) was dissolved in dry DMF (10 mg/mL, 360 μL), after which BODIPY-alkyne (1.1 eq., 14.96 μmol, 4.9 mg), $CuSO_4$ (0.1 M in MilliQ, 0.4 eq., 5.44 μmol, 54.4 μL) and sodium ascorbate (0.1 M in MilliQ, 0.4 eq., 5.44 μmol, 54.4 μL) were added. The reaction mixture was stirred at rt in the dark. After 4 days LCMS confirmed complete consumption of the starting material, after which the reaction mixture was concentrated *in vacuo*. The product was purified by HPLC to yield compound **MH374** (0.6 mg, 1.0 μmol, 7%). $^1$H NMR (850 MHz, MeOD) δ 7.76 (s, 1H, $N_3$C$H$), 7.44 (s, 1H, $H_{Ar}$), 6.98 (d, $J$ = 4.1 Hz, 1H, $H_{Ar}$), 6.30 (d, $J$ = 4.1 Hz, 1H, $H_{Ar}$), 6.22 (s, 1H, $H_{Ar}$), 4.43 (s, 2H, NHC$H_2$), 4.35 (t, $J$ = 7.0 Hz, 2H, C$H_{2,linker}$), 3.85 (s, 1H, H-3), 3.66 – 3.54 (m, 3H, H-2, H-5), 3.24 (t, $J$ = 7.8 Hz, 2H, COC$H_2$), 2.65 (t, $J$ = 7.6 Hz, 2H, COCH$_2$C$H_2$), 2.51 (s, 3H, C$H_3$), 2.35 (s, 1H, H-1), 2.29 (d, $J$ = 5.9 Hz, 6H, C$H_3$, H-6), 2.13 (t, $J$ = 6.4 Hz, 1H, H-4), 1.87 (p, $J$ = 7.2 Hz, 2H, C$H_{2,linker}$), 1.57 – 1.47 (m, 2H, C$H_{2,linker}$), 1.47 – 1.21 (m, 9H, C$H_{2,linker}$). $^{13}$C NMR (850 MHz, MeOD) δ 174.6 ($C_q$), 161.4 ($C_q$), 158.4 ($C_q$), 145.9 ($C_q$), 136.6 ($C_q$), 134.9 ($C_q$), 129.5 ($C_{Ar}$), 125.8 ($C_{Ar}$), 121.4 ($C_{Ar}$), 117.7 ($C_{Ar}$), 81.5 (C-2), 77.5 (C-3), 62.4 (C-5), 52.4 (C-4), 51.3 ($C$H$_{2,linker}$), 49.5 (C-1), 49.4 (C-6), 35.8 (COCH$_2$$C$H$_2$), 35.7 ($C$H$_2$NH), 31.1 ($C$H$_{2,linker}$), 30.3 ($C$H$_{2,linker}$), 27.7 ($C$H$_{2,linker}$), 27.3 ($C$H$_{2,linker}$), 25.6 (CO$C$H$_2$), 24.8 ($C$H$_{2,linker}$), 14.9 ($C$H$_3$), 11.2 ($C$H$_3$). HRMS calculated for $[C_{29}H_{40}BF_2N_7O_4Na]^+$: 622.3095, found: 622.3100

**5-*O*-(*tert*-butyl-dimethyl-silane)-cyclopente-2,3-diol (FK23-037)**

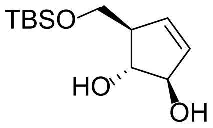

To a solution of cycloalkene **FK-AH006** (700 mg, 2.1 mmol, 1.0 equiv.) in anhydrous $CH_2Cl_2$ (30 mL) under $N_2$ atmosphere was added imidazole (280 mg, 4.14 mmol, 2.0 equiv.). The solution was cooled to 0 °C and TBSCl (410 mg, 1.5 mmol, 1.2 equiv.) was added. The solution was allowed to warm to room temperature and stirred for 16 h. The mixture was diluted with $CH_2Cl_2$ and washed consecutively with sat. aq. solution of $NH_4Cl$ (2 × 50 mL) and water

(50 mL). The organic phase was dehydrated with $Na_2SO_4$, filtered, and concentrated under reduced pressure providing a yellowish oil. The crude material was redissolved in a mixture of anhydrous MeOH and $CH_2Cl_2$ (30 mL, 2:1) under $N_2$ atm. Solid NaOMe (319 mg, 5.90 mmol, 3.0 equiv.) was added and the solution was stirred for 16 h. The mixture was diluted with $CH_2Cl_2$ and washed consecutively with sat. aq. solution of $NaHCO_3$ (50 mL) and water (50 mL). The organic phase was dehydrated with $Na_2SO_4$, filtered and concentrated under reduced pressure. Purification via silica gel flash column chromatography (20% to 70% EtOAc in pentane) yielded cycloalkene **FK23-037** (370 mg, 1.51 mmol, 72%) as colourless solid**.** $^1$H NMR (400 MHz, $CDCl_3$): δ = 5.86 (dt, *J* = 6.0, 2.1 Hz, 1H), 5.70 (dd, *J* = 6.0, 2.2 Hz, 1H), 4.48 (d, *J* = 8.7 Hz, 1H), 3.97 (q, *J* = 3.3 Hz, 1H), 3.82 (dd, *J* = 9.8, 4.2 Hz, 1H), 3.60 (dd, *J* = 9.8, 5.4 Hz, 1H), 2.92 – 2.83 (m, 1H), 2.71 – 2.62 (m, 1H), 2.47 – 2.40 (m, 1H), 0.88 (s, 9H), 0.06 (s, 6H). $^{13}$C NMR (101 MHz, $CDCl_3$): δ = 133.6, 132.7, 83.4, 82.7, 64.1, 54.6, 26.1, 18.5, -5.4. HRMS calculated for $[C_{12}H_{24}O_3SiNa]^+$: 267.13869, found: 267.13876.

**5-*O*-(*tert*-butyl-dimethyl-silane)-arabinofuranose-β-epoxide (FK23-043)**

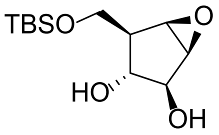

To a solution of cycloalkene **FK23-037** (185 mg, 0.76 mmol, 1.0 equiv.) in anhydrous $CH_2Cl_2$ (40 mL) under $N_2$ atm. was added solid $NaHCO_3$ (191 mg, 2.27 mmol, 3.0 equiv.). To the stirred suspension solid *m*-CPBA (261 mg, 1.51 mmol, 2.0 equiv.) was added at room temperature. After 6 h sat. the mixture was diluted with $NaHCO_3$ (30 mL) and water (20 mL) before extracted with $CH_2Cl_2$ (3 × 100 mL). The combined organic phases were dehydrated with $Na_2SO_4$, filtered and concentrated under reduced pressure. Purification of the crude mixture via silica gel flash column chromatography (70% to 100% $Et_2O$ in pentane) yielded epoxide **FK23-043** (171 mg, 0.66 mmol, 87%) as colourless solid. $^1$H NMR (400 MHz, $CDCl_3$): δ = 4.10 (ddd, *J* = 7.3, 5.8, 1.5 Hz, 1H), 3.93 (dd, *J* = 9.5, 7.1 Hz, 1H), 3.85 (dd, *J* = 9.5, 8.6 Hz, 1H), 3.54 (dd, *J* = 3.1, 1.5 Hz, 1H), 3.49 (td, *J* = 7.5, 6.1, 2.3 Hz, 1H), 3.44 (dd, *J* = 3.1, 1.3 Hz, 1H), 2.46 (d, *J* = 2.7 Hz, 1H), 2.42 (d, *J* = 7.4 Hz, 1H), 2.12 (dtd, *J* = 8.5, 7.2, 1.3 Hz, 1H), 0.90 (s, 9H), 0.09 (s, 6H). $^{13}$C NMR (101 MHz, $CDCl_3$): δ = 79.5, 78.2, 63.3, 56.5, 54.1, 47.9, 26.0, 18.4, -5.4. HRMS calculated for $[C_{12}H_{24}O_4SiNa]^+$: 283.13361, found: 283.13379.

**Arabinofuranose-β-epoxide (FK-AH017)**

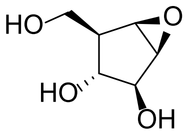

To a solution of epoxide **FK23-043** (10.0 mg, 38.4 µmol, 1.0 equiv.) in anhydrous THF (0.5 mL), solid TBAF on silica (1.5 mmol/g, 51 mg, 76.8 µmol, 2.0 equiv.) was added under argon atm. and stirred for 8 h. The reaction mixture was evaporated to dryness. The crude residue was purified via silica gel flash column chromatography (0% to 15% MeOH in $CH_2Cl_2$) yielded epoxide **FK-AH017** (4.1 mg, 28.1 µmol, 73%) as colourless solid. $^{1}H$ NMR (500 MHz, MeOD): δ = 3.94 (dd, *J* = 6.1, 1.6 Hz, 1H), 3.81 (dd, *J* = 10.6, 4.7 Hz, 1H), 3.67 (t, *J* = 10.1 Hz, 1H), 3.50 – 3.44 (m, 1H), 3.18 (dd, *J* = 7.6, 6.1 Hz, 1H), 2.01 – 1.93 (m, 1H). $^{13}C$ NMR (126 MHz, MeOD): δ = 80.3, 76.0, 61.7, 57.6, 54.9, 49.5. HRMS calculated for $[C_6H_{10}O_4Na]^+$: 169.04713, found: 169.04730.

**2,3,5-*O*-(*tert*-butyl-dimethyl-silane)-arabinofuranose-β-epoxide (FK23-045)**

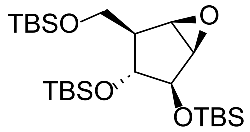

A solution of epoxide **FK23-043** (161 mg, 0.62 mmol, 1.0 equiv.) and imidazole (147 mg, 2.16 mmol, 3.5 equiv.) in anhydrous $CH_2Cl_2$ (20 mL) was cooled to 0°C under nitrogen atm. (ice). TBSOTf (155 µl, 0.68 mmol, 1.1 equiv.) and DMAP (10 mg) were added, and the ice bath was removed. The reaction mixture stirred for 4 h. After full conversion (TLC, $R_f$ = 0.95 in $CH_2Cl_2$) the mixture was diluted with $NaHCO_3$ (20 mL) and water (20 mL) before extracted with $CH_2Cl_2$ (3 × 70 mL). The combined organic phases were dehydrated with $Na_2SO_4$, filtered and concentrated under reduced pressure. Purification of the crude mixture via silica gel flash column chromatography (30% to 100% $CH_2Cl_2$ in pentane) yielded epoxide **FK23-045** (275 mg, 0.56 mmol, 91%) as colourless solid. $^{1}H$ NMR (300 MHz, $CDCl_3$) δ = 4.08 (dd, *J* = 5.4, 1.4 Hz, 1H), 3.83 (dd, *J* = 9.7, 4.5 Hz, 1H), 3.70 (dd, *J* = 10.5, 9.7 Hz, 1H), 3.57 (dd, *J* = 3.2, 1.3 Hz, 1H), 3.43 (dd, *J* = 3.2, 1.4 Hz, 1H), 3.31 (dd, *J* = 7.3, 5.4 Hz, 1H), 2.03 (dddd, *J* = 10.5, 7.3, 4.5, 1.4 Hz, 1H), 0.95 (s, 9H), 0.92 (s, 9H), 0.87 (s, 9H), 0.17 (s, 3H), 0.14 (s, 3H), 0.09 (s, 6H), 0.06 (s, 3H), 0.02 (s, 3H). $^{13}C$ NMR (75 MHz, $CDCl_3$) δ = 81.05, 76.28, 61.62, 56.88, 54.57, 48.78, 26.06, 25.98, 25.83, 18.41, 18.24, 17.85, -3.79, -4.00, -4.20, -4.57, -5.33. HRMS calculated for $[C_{24}H_{53}O_4Si_3]^+$: 489.32462, found: 489.32462.

**2,3,5-*O*-(*tert*-butyl-dimethyl-silane)-arabinofuranose-α-aziridine (FK23-111)**

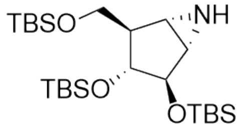

A solution of epoxide **FK23-045** (100 mg, 0.20 mmol, 1.0 equiv.) in anhydrous $CH_2Cl_2$ (20 mL) was cooled to 0°C (ice) under an argon atmosphere. To the solution $TMSN_3$ (534 µl, 4.1 mmol, 20.0 equiv.) and $BF_3{\cdot}Et_2O$ (28 µl, 0.22 mmol, 1.1 equiv.) were added. The solution was slowly warmed up to 15 °C over a period of 3 h. After full conversion (TLC, $R_f$ = 0.83 in $CH_2Cl_2$) the mixture was diluted with $NaHCO_3$ (20 mL) and extracted with $CH_2Cl_2$ (3 × 40 mL). The combined organic phases were dehydrated with $Na_2SO_4$, filtered and concentrated under reduced pressure. The crude was dissolved in anhydrous $CH_2Cl_2$ (10 mL) followed by the addition of TES-H (120 µL). The mixture was cooled on ice after which TFA (310 µL) was added, the solution was stirred for 1 hour while attaining to room temperature. The reaction was quenched by the addition of sat. aq. $NaHCO_3$ (10 mL) and water (10 mL). The aqueous phase was extracted with $CH_2Cl_2$ (3 × 40 mL). The combined organic phases were dehydrated with $Na_2SO_4$, filtered and concentrated under reduced pressure. Polar residues of the leftover were removed by gel filtration ($CH_2Cl_2$ /pentane, 1:1) yielded crude azido alcohol **FK23-108** (101 mg, 0.19 mmol, 93%, isomeric mixture) as colourless solid.

To a solution of crude azido alcohol **FK23-108** (55 mg, 0.10 mmol, 1.0 equiv.) in anhydrous $CH_2Cl_2$ (3.0 mL), $Et_3N$ (43 µL, 0.30 mmol, 3.0 equiv.) and DMAP (1.3 mg, 10 µmol, 0.1 equiv.) were added at 0°C (ice). MsCl (10 µL, 0.13 mmol, 1.3 equiv.) was added subsequently and the mixture was allowed to warm up to room temperature and stirred overnight. The reaction mixture was diluted with sat $NH_4Cl$ (10 mL) and extracted with $CH_2Cl_2$ (3 × 20 mL). The combined organic phases were dehydrated with $Na_2SO_4$, filtered and concentrated under reduced pressure.

The crude was redissolved in anhydrous THF (3.0 mL) under an argon atmosphere followed by the addition of $PPh_3$ (54 mg, 0.21 mmol, 2.0 equiv.). The reaction mixture was stirred for 4 h at room temperature before DIPEA (200 µL) and $H_2O$ (400 µL) were added. After 16 h the reaction mixture was further diluted with $H_2O$ (5.0 mL) and the aqueous phase was extracted with EtOAc (3 × 20 mL). The combined organic phases were washed with brine, dehydrated with $Na_2SO_4$, filtered and concentrated under reduced pressure. Purification of the crude mixture via silica gel flash column chromatography (0% MeOH to 1% MeOH in $CH_2Cl_2$) yielded aziridine **FK23-111** (32 mg, 66 µmol, 64%) as colourless oil. $^1H$ NMR (500 MHz, $CDCl_3$) δ = 3.97 (s, 1H), 3.85 (s, 1H), 3.64 (dd, *J* = 9.8, 7.1 Hz, 1H), 3.57 (t, *J* = 9.4 Hz, 1H), 2.38 (s, 1H), 2.33 (s, 1H), 2.22 (dd, *J* = 9.1, 7.1 Hz, 1H), 0.90 (s, 9H), 0.89 (s, 9H), 0.85 (s, 9H), 0.10 (s, 3H), 0.09 (s, 3H), 0.07 – 0.03 (m, 12H). $^{13}C$ NMR (126 MHz, $CDCl_3$) δ = 80.12, 79.37, 62.98, 52.32, 40.43, 37.78, 25.95, 25.77, 25.72, 18.33, 17.97, 17.77, -4.70, -4.81, -4.92, -5.37, -5.40. HRMS calculated for $[C_{24}H_{53}NO_3Si_3]^+$: 488.34060, found: 488.34013.

**Arabinofuranose-α-aziridine (FK23-115) aka 4**

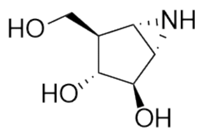

FK23-111 (9.0 mg, 18.3 µmol, 1.0 equiv.) was dissolved in THF (600 µL) in a 2.0 mL Eppendorf vial (PP). To the solution TBAF (1.0M in THF, 92 µL, 5.0 equiv.) was added and the reaction mixture was stirred for 3 h at room temperature. To the mixture $SiO_2$ was added and stirred for additional 20 minutes. The crude reaction mixture was directly purified via silica gel flash column chromatography (neutral silica, 5% to 100% acetone in $CH_2Cl_2$). To remove remaining TBA salts the product was loaded on neutral Amberlite 120 $H^+$ (pH adjustment via $NH_4OH$). The Amberlite was washed with $H_2O$ (3x 10 mL) followed by releasing of the product by rinsing the Amberlite with $NH_4OH$ (1M, 10 mL). After 3 cycles of lyophilization aziridine **FK23-115** was yielded (1.3 mg, 8.9 µmol, 49%) as colourless solid. $^1H$ NMR (500 MHz, $D_2O$) δ = 4.03 (s, 1H), 3.87 (s, 1H), 3.65 (s, 1H), 3.64 (s, 1H), 2.60 (s, 2H), 2.21 (t, $J$ = 6.9 Hz, 1H). $^{13}C$ NMR (126 MHz, $D_2O$) δ = 79.34, 76.42, 61.07, 49.88, 40.03, 38.52. HRMS calculated for $[C_6H_{12}NO_3]^+$: 146.08117, found: 146.08128.

*Solvents, TLC visualization and NMR.*

Reagents and solvents were purchased from commercial sources and were used without purifying unless stated otherwise. Anhydrous solvents were dried and stored over activated 3 Å or 4 Å molecular sieves under a N2 atmosphere unless stated otherwise. Traces of water were removed by co-evaporation with anhydrous toluene (if indicated, under an argon atmosphere) for reactions or analysis requiring anhydrous conditions. Reaction progress was monitored via TLC analysis on aluminum sheets ((TLC silica gel 60 F254, Merck Supelco®) or (ALUGRAM® Xtra SIL G UV254, Macherey-Nagel)). Compounds were visualized using UV-detection (254 nm or 366 nm) and/or sprayed with a cerium molybdate spray (solution of (NH4)6Mo7O24·H2O (25 g/L) and (NH4)4Ce(SO4)4·2H2O (10 g/L) in 10% aq. sulfuric acid) or a sulfuric acid spray (5% sulfuric acid in EtOH, v/v) or a ninhydrin spray (solution of ninhydrin (1 g/L) in EtOH/AcOH, 20/1, v/v) or an alizarin spray (1 mM alizarin in MeOH) followed by charring at ±200°C or a KMnO4 spray (KMnO4 (20 g/L) and K2CO3 (10 g/L) in water) followed by heating with a heat gun.

1H, 11B, 13C, and 31P NMR spectra were recorded on Bruker AV-400 (400 MHz), Bruker AV-400 Wide Bore (400 MHz), Bruker AV-500 (500 MHz), Bruker AV-600 (600 MHZ), and Bruker AV-850 (850 MHz) instruments. Deuterated chloroform was stored over K2CO3 and activated 3 Å molecular rods (size 1/16 in., Sigma Aldrich). 11B, 19F and 31P chemical shifts are indirectly referenced to BF3·OEt2, CFCl3 and H3PO4 (0.00 ppm), respectively, according to the IUPAC method. Chemical shifts are denoted by δ and reported in parts per million (ppm) and are relative to the residual signal of the solvent or of tetramethylsilane (TMS) (0.00 ppm) in the case of 1H spectra measured using deuterated chloroform with 0.01% TMS as additive. Coupling constants are denoted by the letter J and reported in Hz. All 13C NMR spectra are proton decoupled, measured using the attached proton test (APT) experiment, and are presented with even signals (Cq and CH2) as positive and odd signals (CH and CH3) as negative. Structural assignments were made with additional information from HH-COSY, HH-NOESY, HSQC, and HMBC experiments.

*Devices managed by Koen*

*TLC-MS* analysis was performed on an Advion Interchim Scientific expression-L® CMS (ESI) coupled to a Plate Express® Automated TLC plate reader (eluted with MeOH/H2O/formic acid, 9/1, v/v + 0.1% formic acid, flow rate 0.12 mL/min).

LC-MS analysis was performed on a Thermo Scientific Vanquish Focused UHPLC+ system (split sampler FT, Quaternary pump F, DAD detector HL) coupled to a Thermo LCQ Fleet ion-trap mass-spectrometer (LCF10684) (ESI) equipped with a C18 column (NUCLEODUR C18 Gravity, 4.6 × 50 mm, 3 µm, Macherey-Nagel) – in combination with a solvent system consisting of A = Milli-Q® H2O, B = acetonitrile; linear gradient of B in A + 0.1% TFA , 1 mL/min.

*Tandem mass spectrometry (MS/MS)* was performed on an Agilent 1260 Infinity II LC system (G7129A vial sampler, G7115A DAD WR, G7111B quaternary pump) coupled to an Agilent 6475 LC/TQ (G6475A, triple quadrupole) (ESI) equipped with a C18 column (NUCLEODUR C18 Gravity, 4.6 × 50 mm, 3 µm, Macherey-Nagel) ) or C18AQ column (Develosil RPAQEOUS, 3 µm, 100 × 4.6 mm, NOMURA (via Phenomenex)) or C30 column (Fusion (now FlexFire) C30, 5 µm, 150 × 4.6 mm, NOMURA (via Phenomenex)) or “C6” column (NUCLEODUR Phenyl-Hexyl, 3 µm, 50 × 4.6 mm, Macherey-Nagel) or C4 column (NUCLEOSIL 120-5 C4, 5 µm, 150 × 4.6 mm, Macherey-Nagel) or “C2” column (Vydac 219TP, 5 µm, 150 × 4.6 mm, GRACE) – in combination with a solvent system consisting of A = Milli-Q® H2O, B = acetonitrile; linear gradient of B in A + additive (0.1% TFA or 0.1% formic acid or 0.1% acetic acid or 10 mM NH4OAc or 10 mM NH4HCO3 (only when using NUCLEODUR C18), or 10 mM HNEt3OAc), 1 mL/min. Alternatively, highly polar compounds were separated using the same system equipped with a HILIC column (NUCLEODUR HILIC, 3 µm, 50 × 4.6 mm, Macherey-Nagel) or “Diol” column (YMC-Pack Diol-120, 5 µm, 150 × 4.6 mm, YMC) – in combination with a solvent system consisting of A = Milli-Q® H2O and B = acetonitrile; linear gradient of A in B + additive (0.1% formic acid or 0.1% acetic acid or 10 mM NH4OAc), 1 mL/min. Product ions were determined and optimized beforehand using a column bypass.

*High-resolution mass spectra (HRMS)* were recorded by direct injection on a Thermo Scientific Q Exactive HF Orbitrap mass spectrometer coupled to an Ultimate 3000 Nano LCEN ESI probe (source voltage 3.5kV, no sheath gas flow, capillary temperature 275 °C) with resolution R = 240.000 at m/z=400 (mass range m/z 160-2000 or until a maximum of 6000). Calculated and observed m/z values refer to the exact mass of the [M+X] species. Samples were introduced by flow injection at 25 µl/min (ACN/MilliQ® H2O, 1/1, v/v + 0.1% FA) using the loading pump. HRMS samples were prepared with an approximate concentration of 1 µM using a mixture of acetonitrile and Milli-Q® water (1/1, v/v), 2 µL injection.

*Semi-preparative high performance liquid chromatography (semiprep-HPLC)* was performed on an Agilent 1260 Infinity II LC system (G1761A binary pump, G7157A autosampler, G7114A VWD detector, G1364E fraction collector, G7110B iso pump, G7170B MS flow modulator) coupled to an Agilent InfinityLab mass spectrometer (G6135B LC/MSD XT) (ESI) equipped with a semi-preparative C18 column (NUCLEODUR C18 Gravity, 5 µm, 250 × 10 mm, Macherey-Nagel) or C30 column (Fusion (now FlexFire) C30, 5 µm, 250 × 10 mm, NOMURA (via Phenomenex)) or C18AQ column (Develosil RPAQEOUS, 5 µm, 250 × 10 mm, NOMURA (via Phenomenex)) or "C6" column (NUCLEODUR Phenyl-Hexyl, 5 µm, 250 × 10 mm, Macherey-Nagel) or C4 column (NUCLEOSIL 120-5 C4, 5 µm, 250 × 10 mm, Macherey-Nagel) or "C2" column (Vydac 219TP, 5 µm, 250 × 10 mm, GRACE) – in combination with a solvent system consisting of A = 50 mM NH4OAc in Milli-Q® H2O or A = 50 mM NH4HCO3 in Milli-Q® H2O (only when using NUCLEODUR C18) or A = 0.2% TFA in Milli-Q® H2O or A = 1% AcOH in Milli-Q® H2O and B = acetonitrile; linear gradient of B in A, 5 mL/min. Alternatively, highly polar compounds were separated using the same system equipped with a semi-preparative HILIC column (NUCLEODUR HILIC, 5 µm, 250 × 10 mm, Macherey-Nagel) – in combination with a solvent system consisting of A = 50 mM NH4OAc in Milli-Q® H2O or A = 1% AcOH in Milli-Q® H2O and B = acetonitrile; linear gradient of A in B, 5 mL/min.

*Preparative high-performance liquid chromatography (prep-HPLC)* was performed on a Waters AutoPurification LC system (Waters 2767 sample manager, Waters 2545 binary gradient module, Waters SFO System Fluidics Organizer, Waters 515 HPLC pump, Waters 2998 PDA detector, Waters Flow Splitter) attached to a Waters SQ detector Acquity Ultra Performance LC equipped with a preparative C18 column (NUCLEODUR C18 Gravity, 5 µm, 150 × 21 mm, Macherey-Nagel) in combination with a gradient consisting of A = Milli-Q® H2O and B = acetonitrile or A = 0.2% TFA in Milli-Q® H2O and B = acetonitrile; linear gradient of B in A, 15 mL/min.

*Automated flash column chromatography* was performed on Biotage® Isolera™ or Selekt™ systems using pre-packed cartridges purchased from Screening Devices B.V. (Ultrapure Irregular Silica Gel, 40-63 µm, 60Å) unless stated otherwise.

### Supplementary Figures

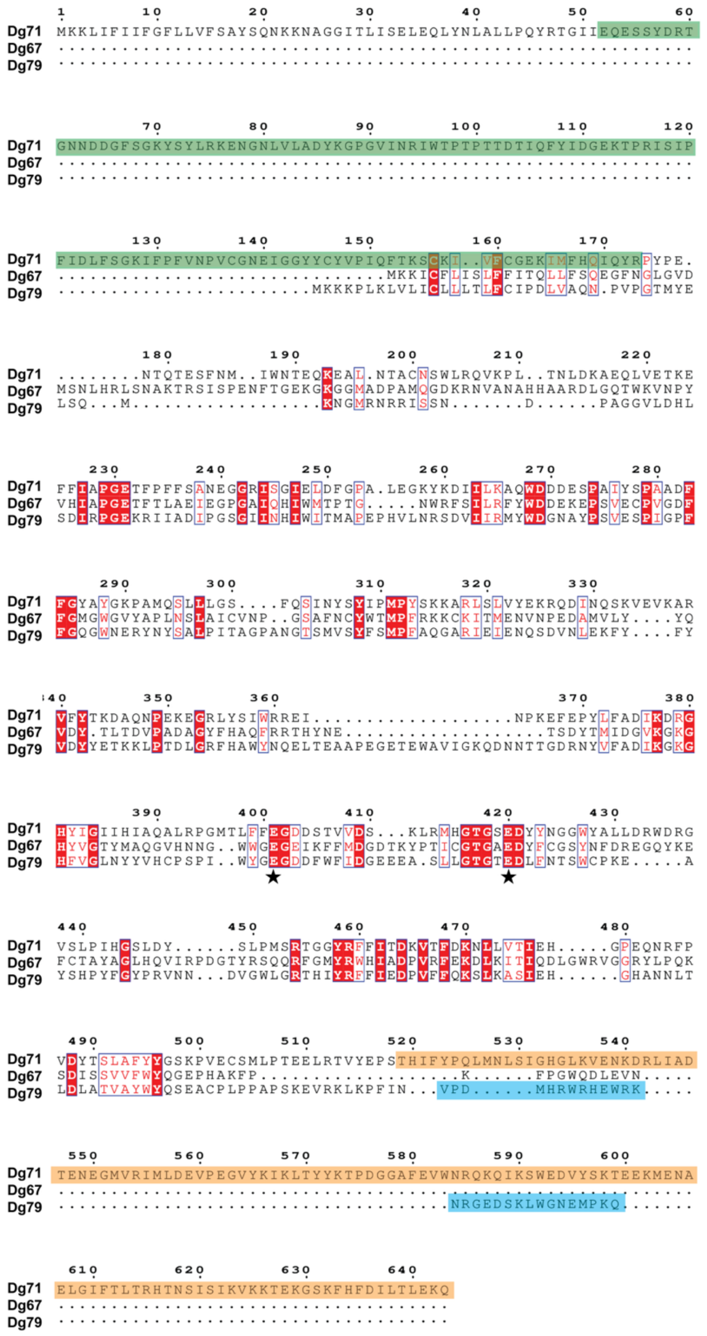

**Figure S1: Protein sequence alignment of native *D. gadei* GH172 proteins highlighting key residues.**

Residues with a red background are conserved and red text indicates partially conserved residues. Blue boxed residues show extended regions of sequence conservation. Catalytic residues are highlighted with black stars. The additional DJR in Dg71 which completes its active site is highlighted in green, and its β-domain which allows dimerisation is shown in orange. The α-helix in Dg79 which interlocks with another allowing the dimerisation of trimers, is shown in blue. Protein sequences were aligned using Clustal Omega[16] and formatted using ESPript3.[17]

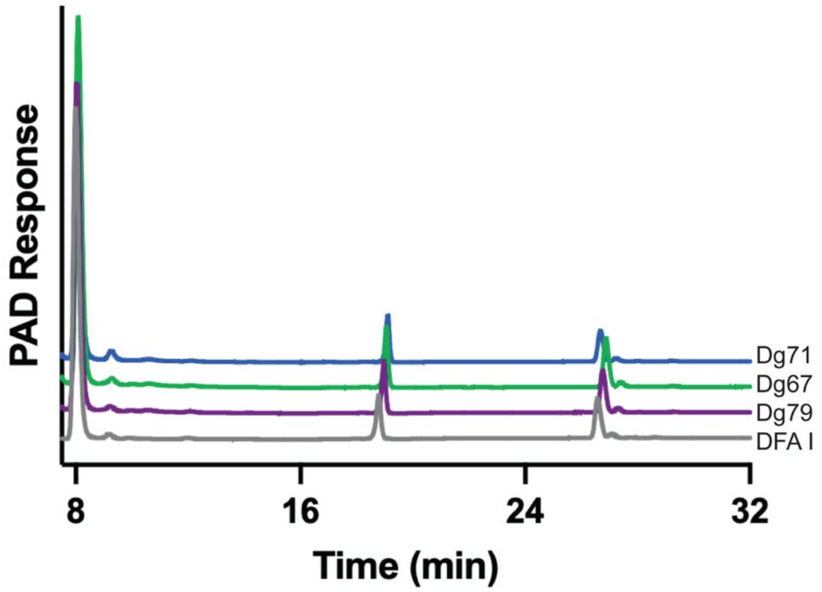

**Figure S2: *D. gadei* GH172 enzymes display no activity on DFA I**

Dg67, Dg71, and Dg79 (1 µM) were incubated overnight with DFA I (2 mg/ml) and analysed by IC-PAD.

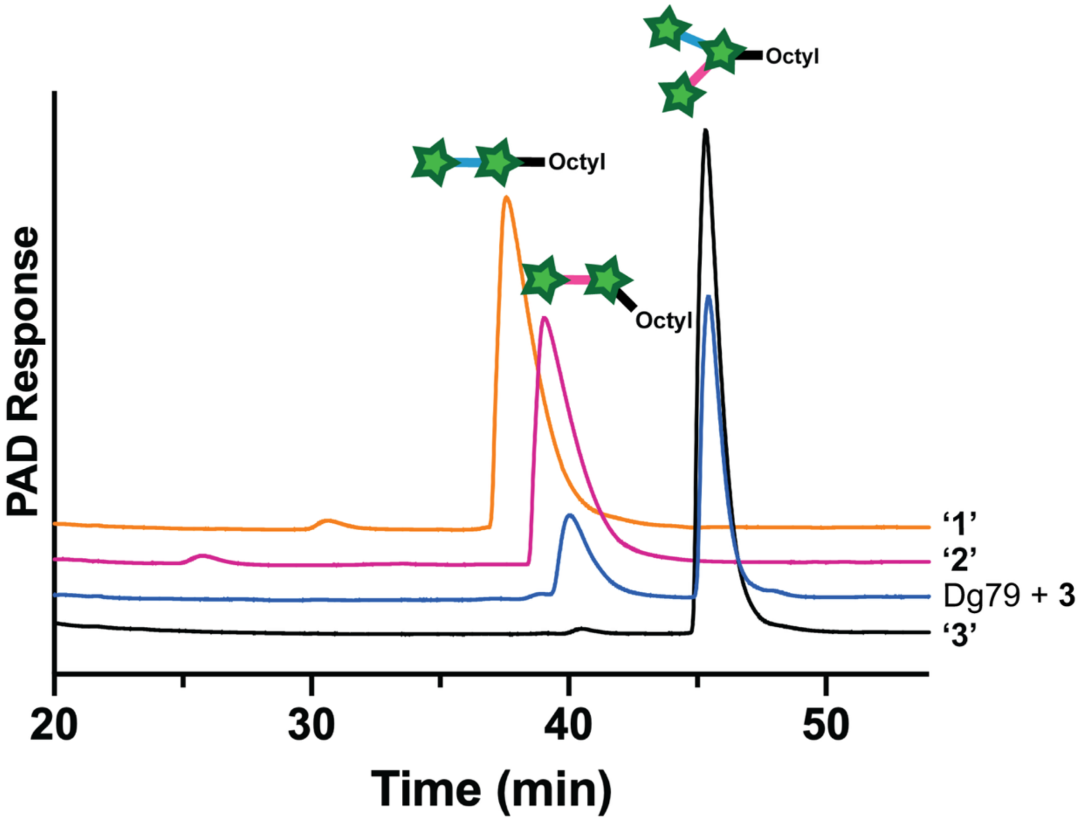

**Figure S3: Dg79 preferentially removes the 1,5-linked ᴅ-Ara*f* from branched substrate 3**

Dg79 (1 µM) was incubated with **3** (0.5 mM) for 16 hours and analysed by IC-PAD.

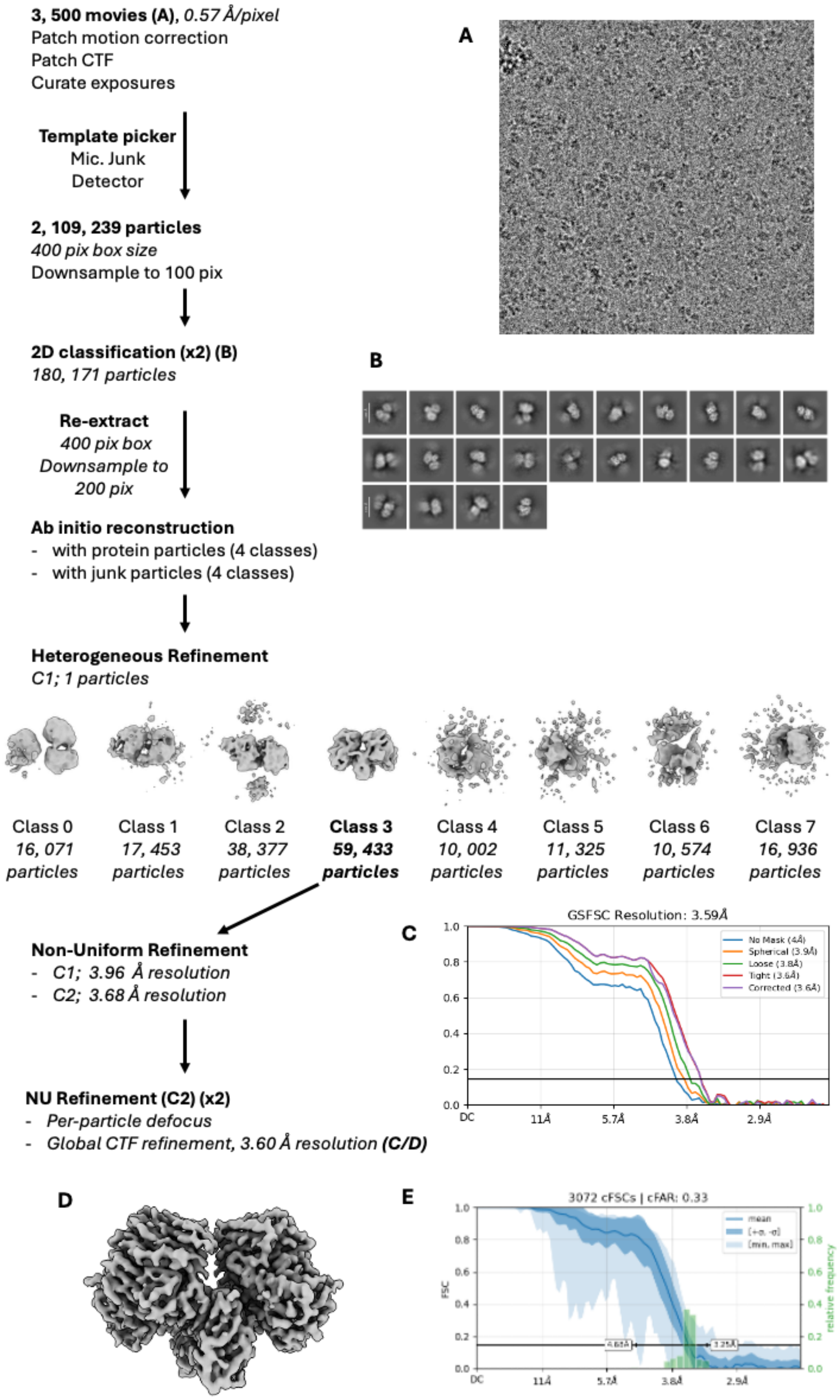

**Figure S4: CryoSPARC processing pipeline for Dg71**

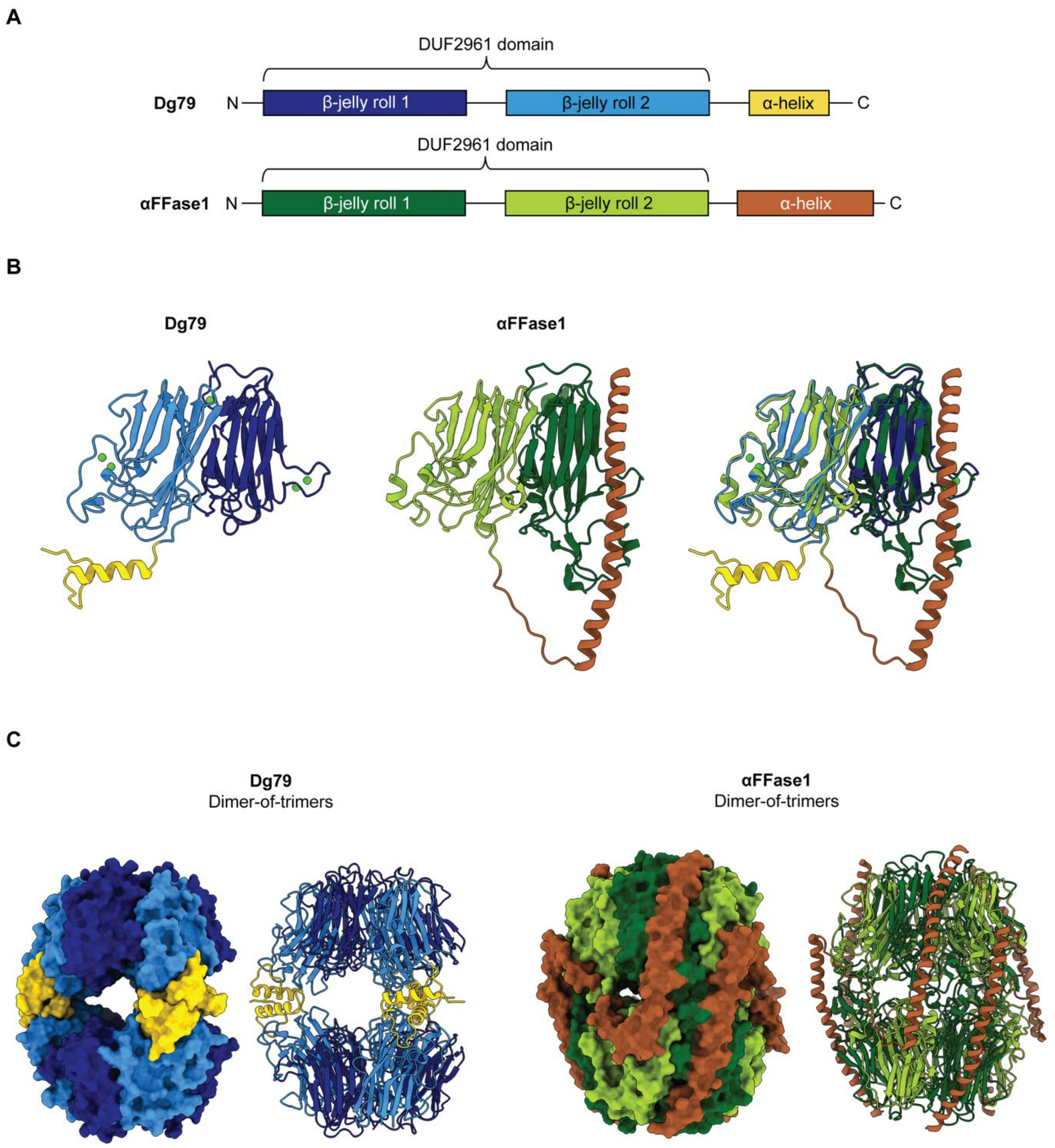

**Figure S5: Structural comparison of Dg79 and αFFase1**

**A:** Domain schematics of Dg79 and αFFase1 showing the longer α-helical domain in αFFase1. **B:** Monomeric subunits of Dg79 and αFFase1 coloured by domain schematics. Overlay of the monomers on the right. **C:** Quaternary structures of Dg79 and αFFase1 shown in surface and cartoon representations.

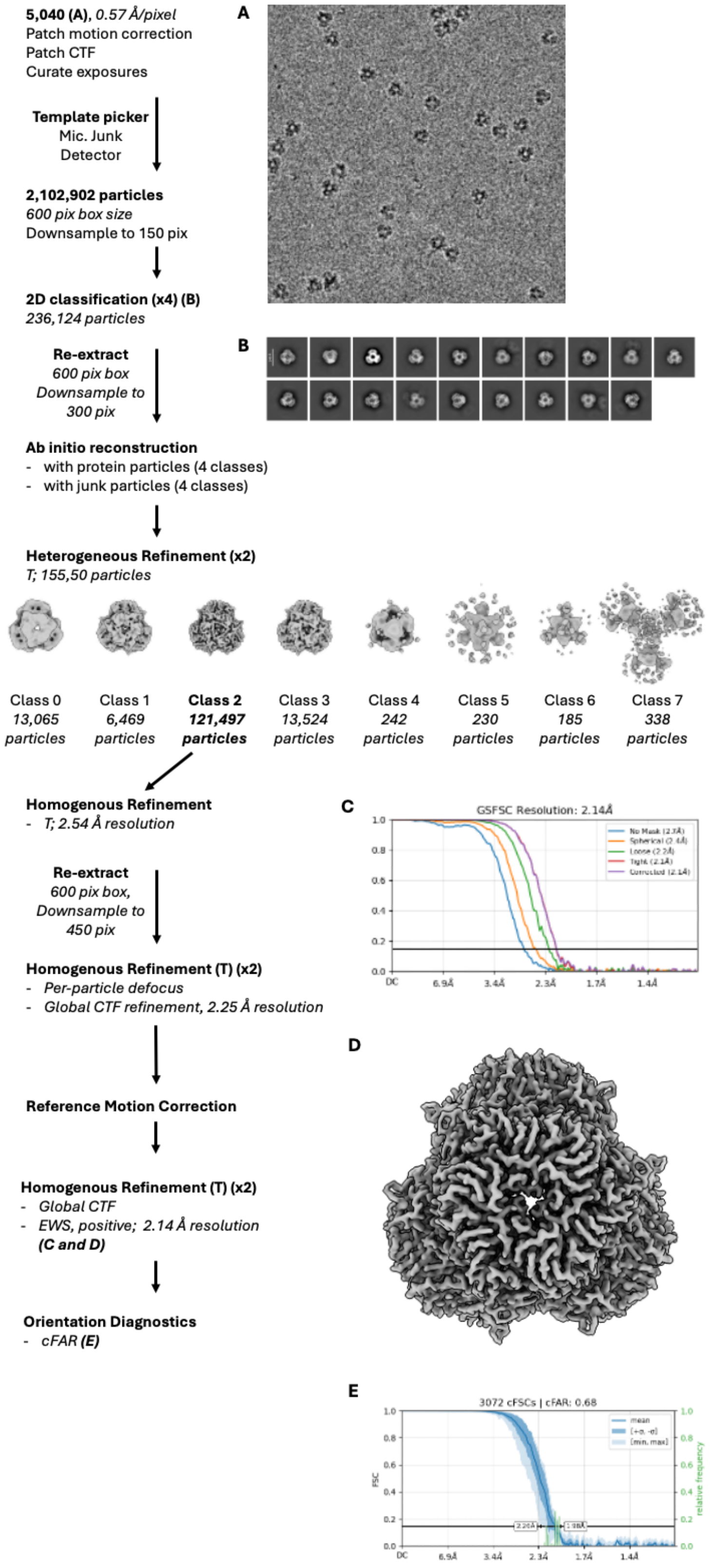

**Figure S6: CryoSPARC processing pipeline for Dg67**

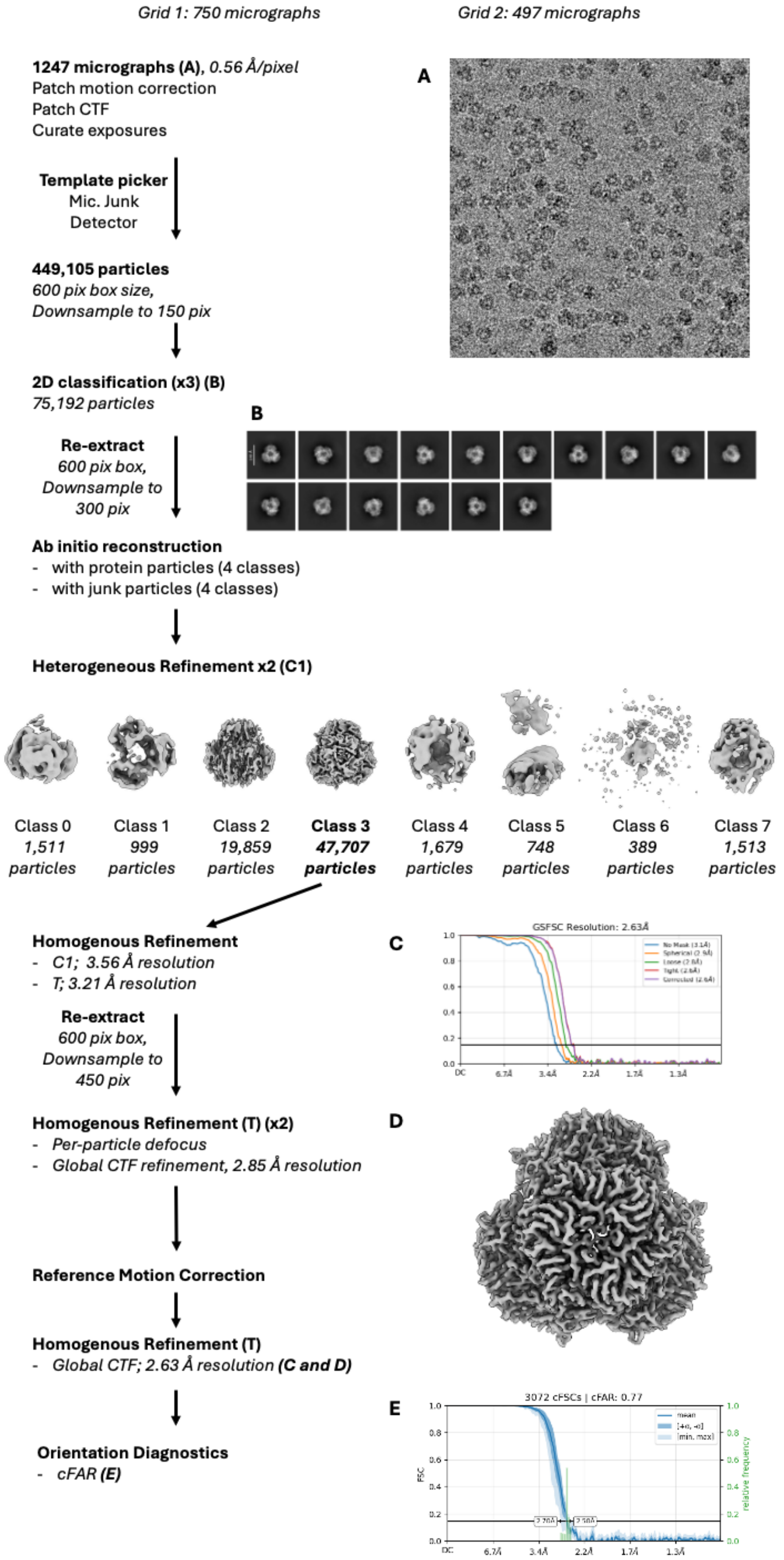

**Figure S7: CryoSPARC processing pipeline for Dg67 incubated with D-arabinose**

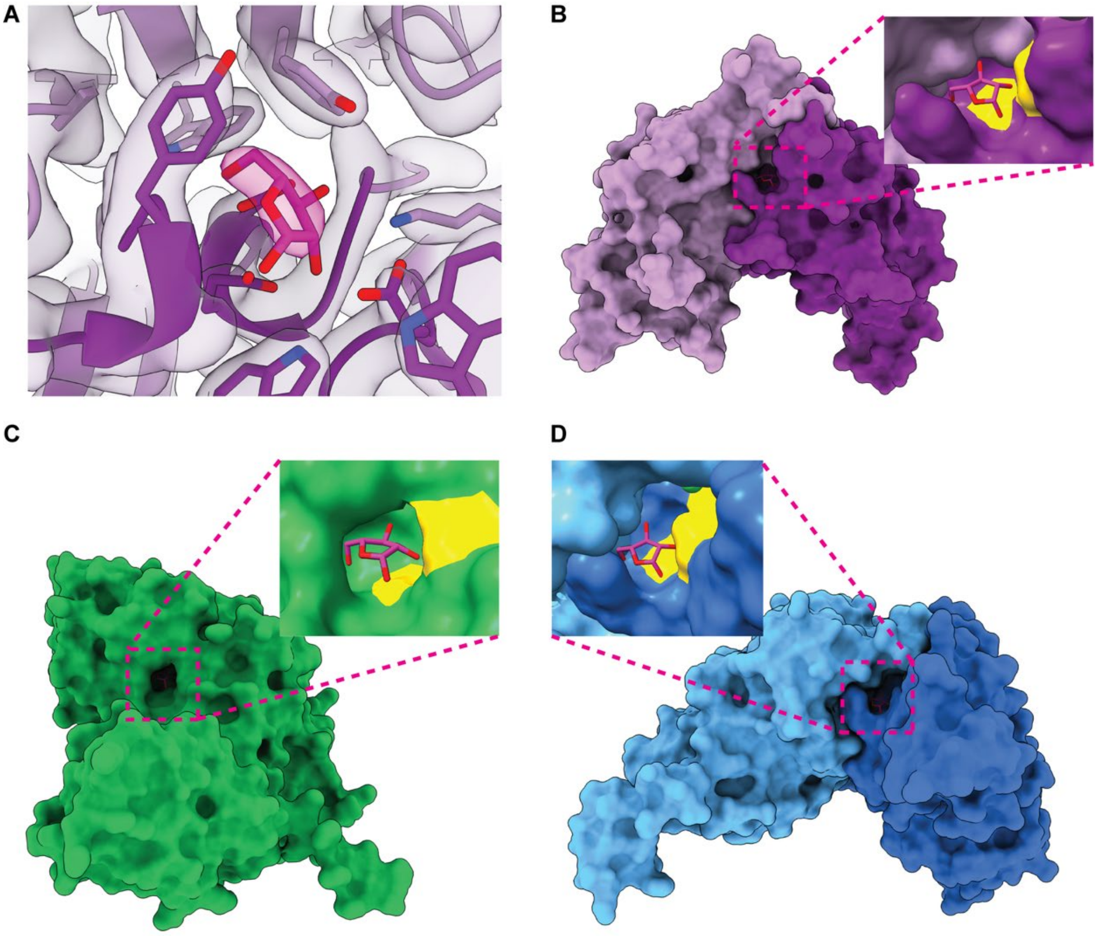

**Figure S8: Dg67 with bound D-Ara*f*.**

**A**: Visualisation of cryo-EM density of the active site of Dg67 with D-Ara*f* bound. The D-Ara*f* density is shown in pink. **B**: The solvent accessible surface area of the Dg_02467 with D-Ara*f* cryo-EM structure showing D-Ara*f* (pink) in the active site pocket. **C/D**: Superimposition of Dg71 (**C**) and Dg79 (**D**) with the D-Ara*f* molecule bound to Dg67 (pink). Subpanels in **B**, **C** and **D** show zoomed in active site with catalytic residues shown in yellow.

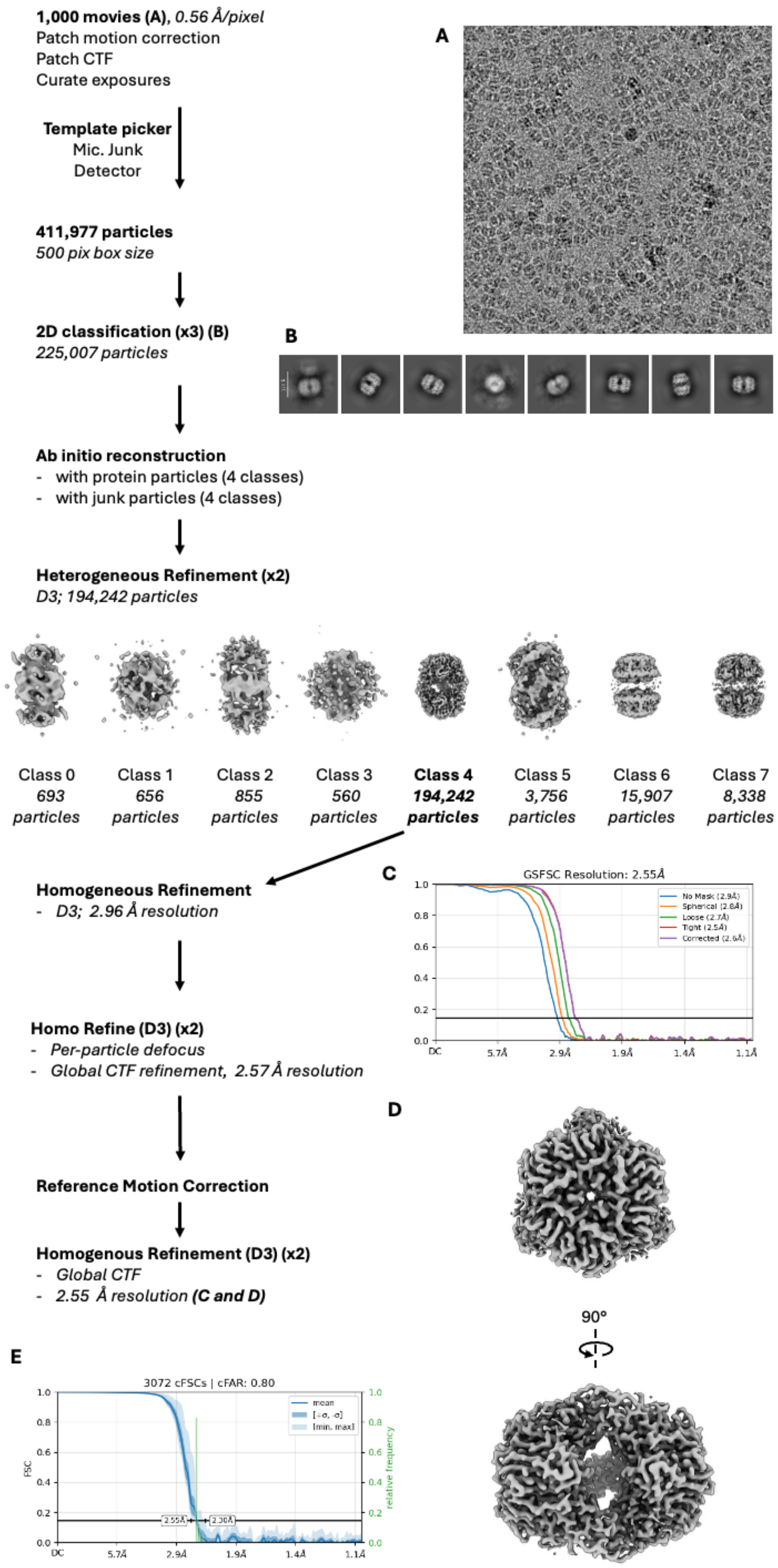

**Figure S9: CryoSPARC processing pipeline for Dg79**

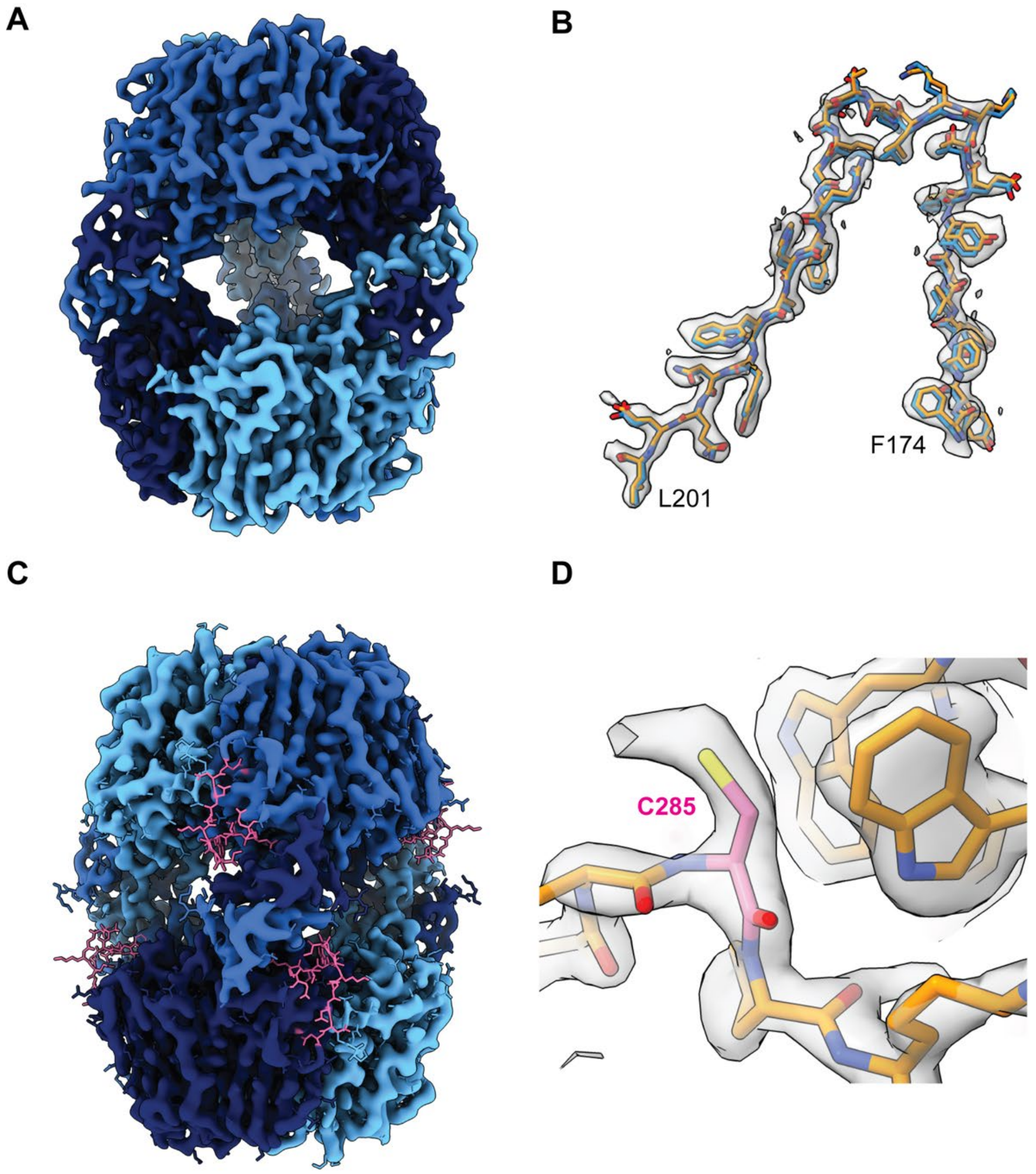

**Figure S10: Dg79 crystal structure docked into Dg79 cryo-EM map**

**A**: The cryo-EM electronic potential map of Dg79 with D3 imposed symmetry. Monomers of Dg79 are coloured in shades of blue. **B**: Representative area of the atomic model from the crystal structure of Dg79 (blue) and the cryo-EM model (orange) docked into the cryo-EM map. **C**: Cryo-EM map highlighting the missing density from the loop E207 to N221 (pink in atomic model). **D**: Extra density around C285 (pink) from a grid freezing modification.

**Ara*f*** [illegible]

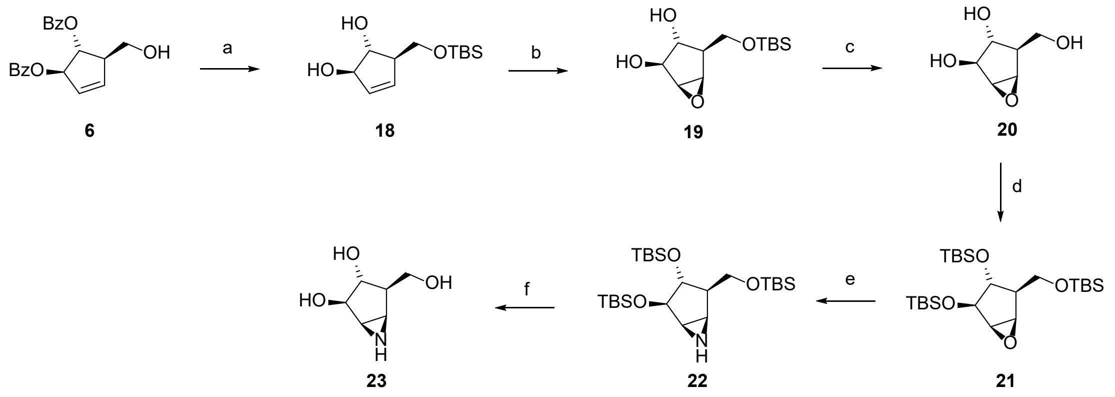

a) 1) TBSCl, imidazole, DCM; 2) NaOMe, MeOH/DCM (2:1), 72%; b) *m*-CPBA, $NaHCO_3$, DCM, 87%; c) TBAF, THF, 73%; d) TBSOTf, imidazole, DMAP, DCM, 91%; e) 1) $TMSN_3$, $BF_3$•$OEt_2$, DCM, 2) TES-H, TFA, DCM; 3) MsCl, DMAP, $Et_3N$, DCM; 4) $PPh_3$, DIPEA, $H_2O$, THF, 64%; f) TBAF, $SiO_2$, THF, 49%.

**Figure S11: Synthesis scheme for aziridine inhibitor 4**

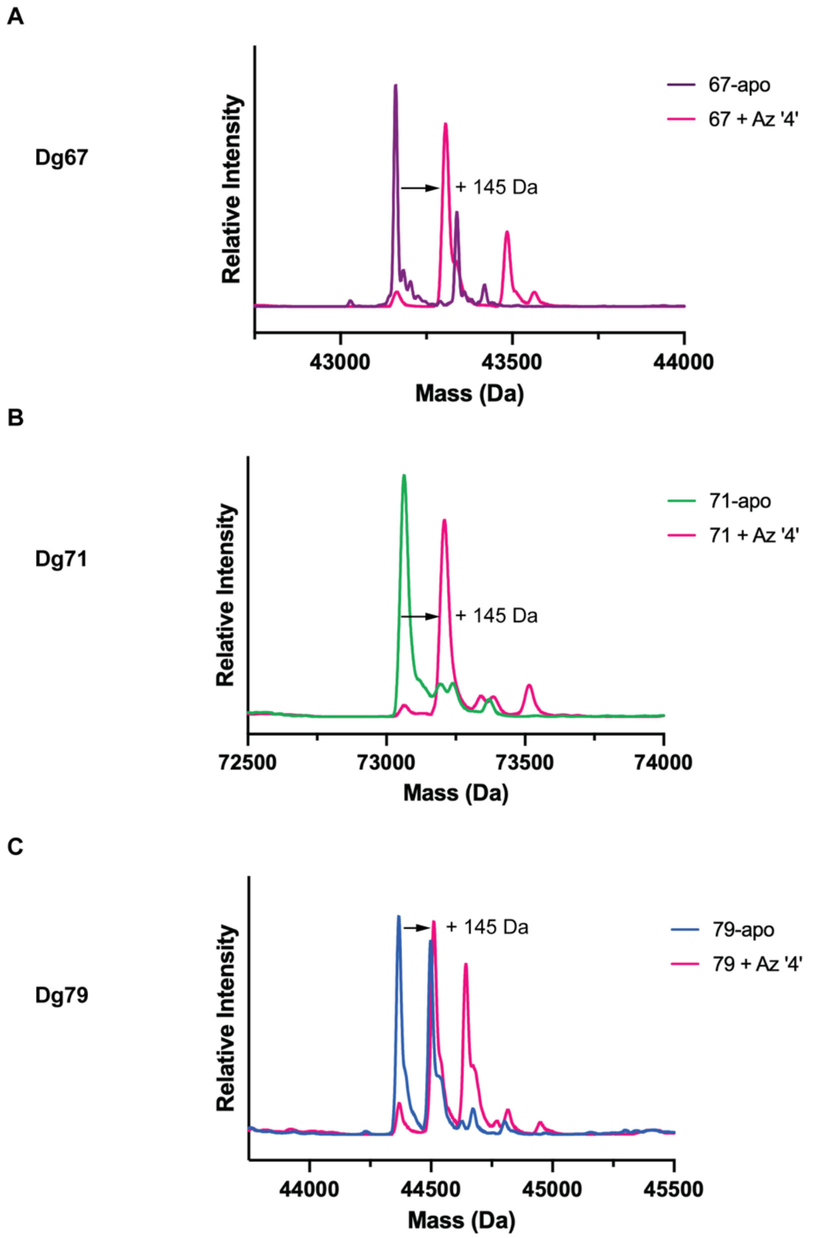

**Figure S12: LC-MS of aziridine inhibition of GH172 enzymes**

Intact mass spectra of Dg67 (**A**), Dg71 (**B**), and Dg79 (**C**) in apo form (purple, green and blue respectively) and after reaction with **4** (pink).

**Araf** monosaccharide BODIPY probe

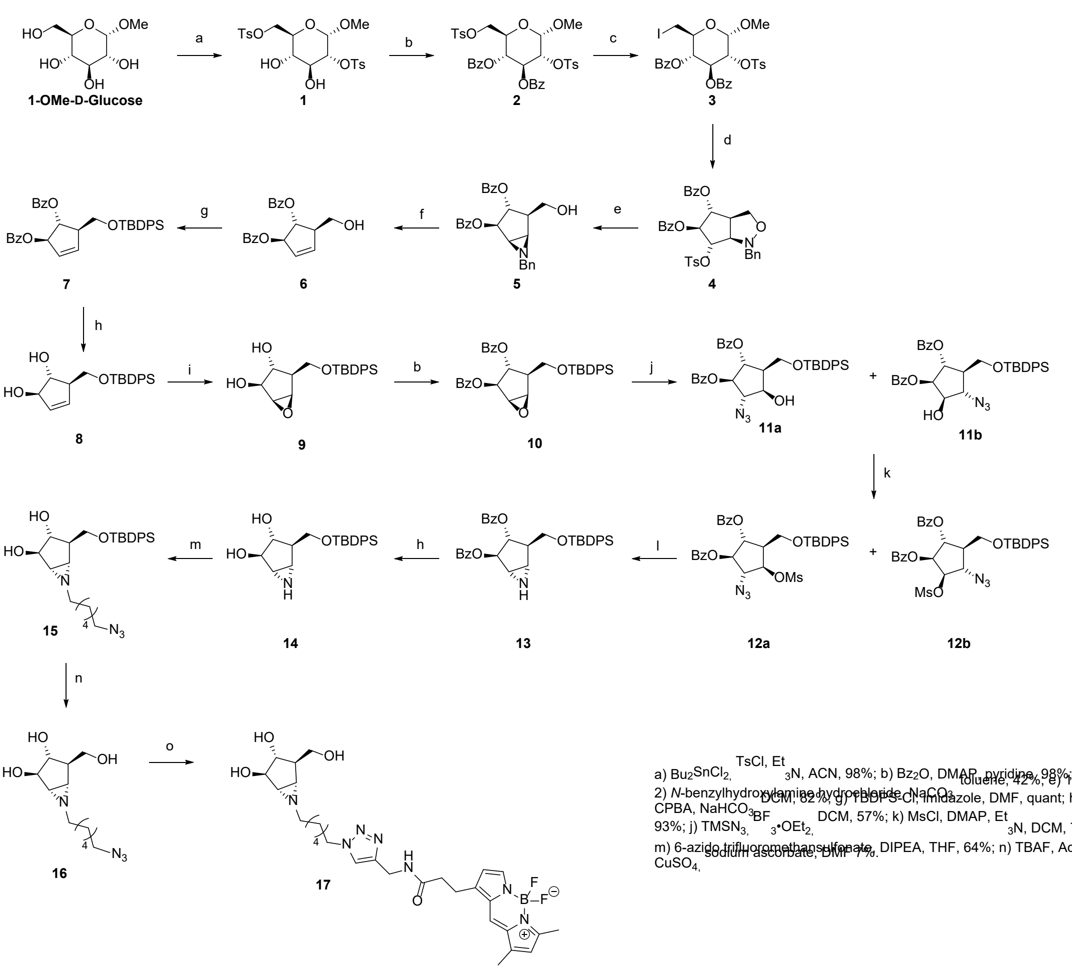

a) $Bu_2SnCl_2$, TsCl, $Et_3N$, ACN, 98%; b) $Bz_2O$, DMAP, pyridine, 98%; c) NaI, $Ac_2O$, toluene, 42%; d) 1) Zn, $NH_4Cl$, EtOH; 2) *N*-benzylhydroxylamine hydrochloride, $NaCO_3$, DCM, 62%; e) 1) Zn, AcOH; 2) $Et_3N$, MeOH, 70%; f) *m*-CPBA, $NaHCO_3$, DCM, 57%; g) TBDPS-Cl, imidazole, DMF, quant; h) NaOMe, MeOH, 90%; i) *m*-CPBA, DCM, 93%; j) $TMSN_3$, $BF_3$•$OEt_2$, DCM, 57%; k) MsCl, DMAP, $Et_3N$, DCM, 78%; l) $PPh_3$, DIPEA, THF/$H_2O$ (4:1), 73%; m) 6-azido trifluoromethansulfonate, DIPEA, THF, 64%; n) TBAF, AcOH, THF, quant.; o) BODIPY-alkyne, $CuSO_4$, sodium ascorbate, DMF 7%.

**Figure S13: Synthetic scheme for BODIPY-probe 5**

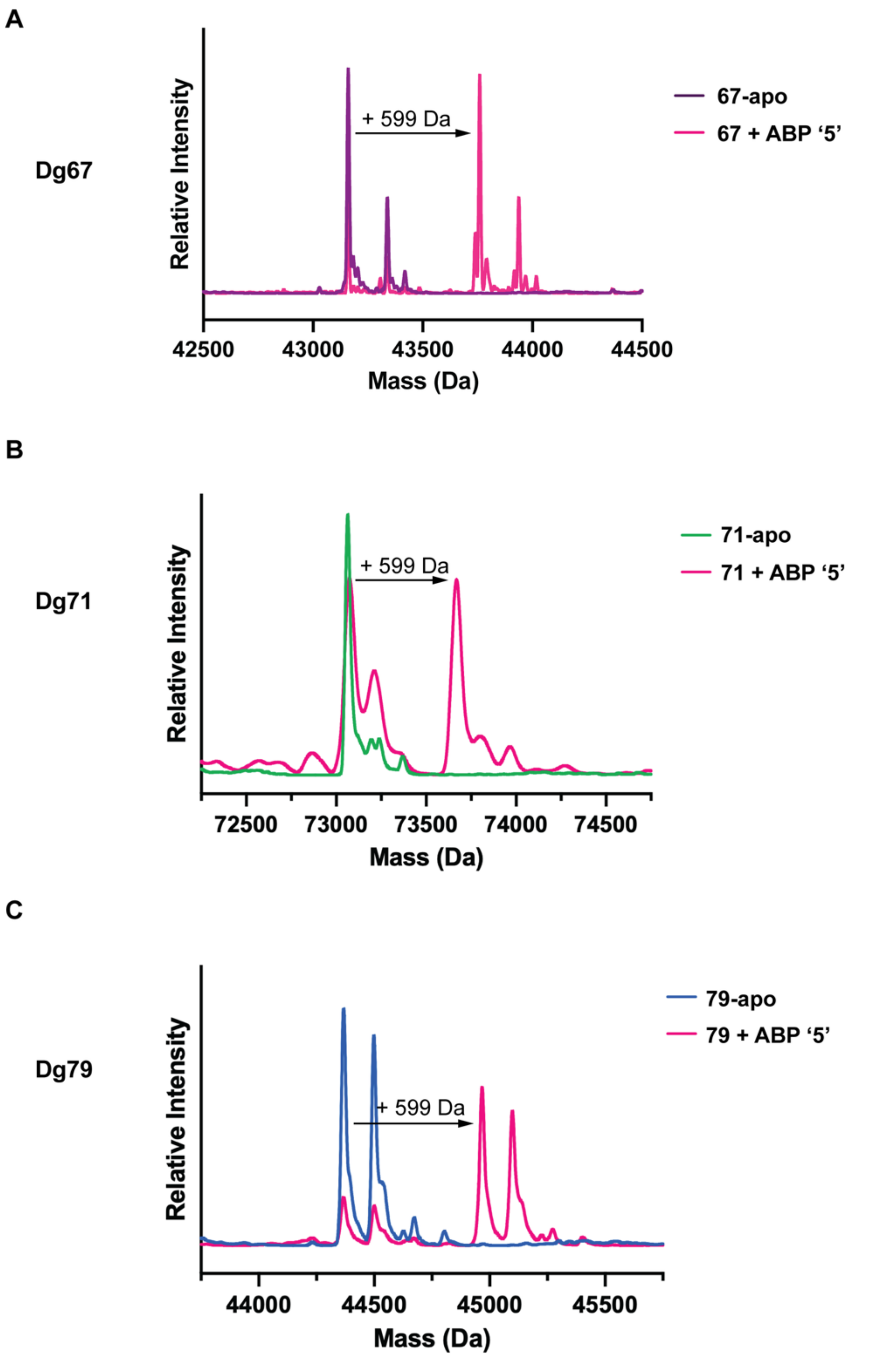

**Figure S14: LC-MS spectra of Dg67, Dg71, and Dg79 after incubation with ABP 5.**

Intact mass spectra of Dg_02467 (**A**), Dg_02471 (**B**), and Dg_02479 (**C**) in apo form (purple, green and blue respectively) and after reaction with ‘**5**’ (pink, MW=599.49 Da).

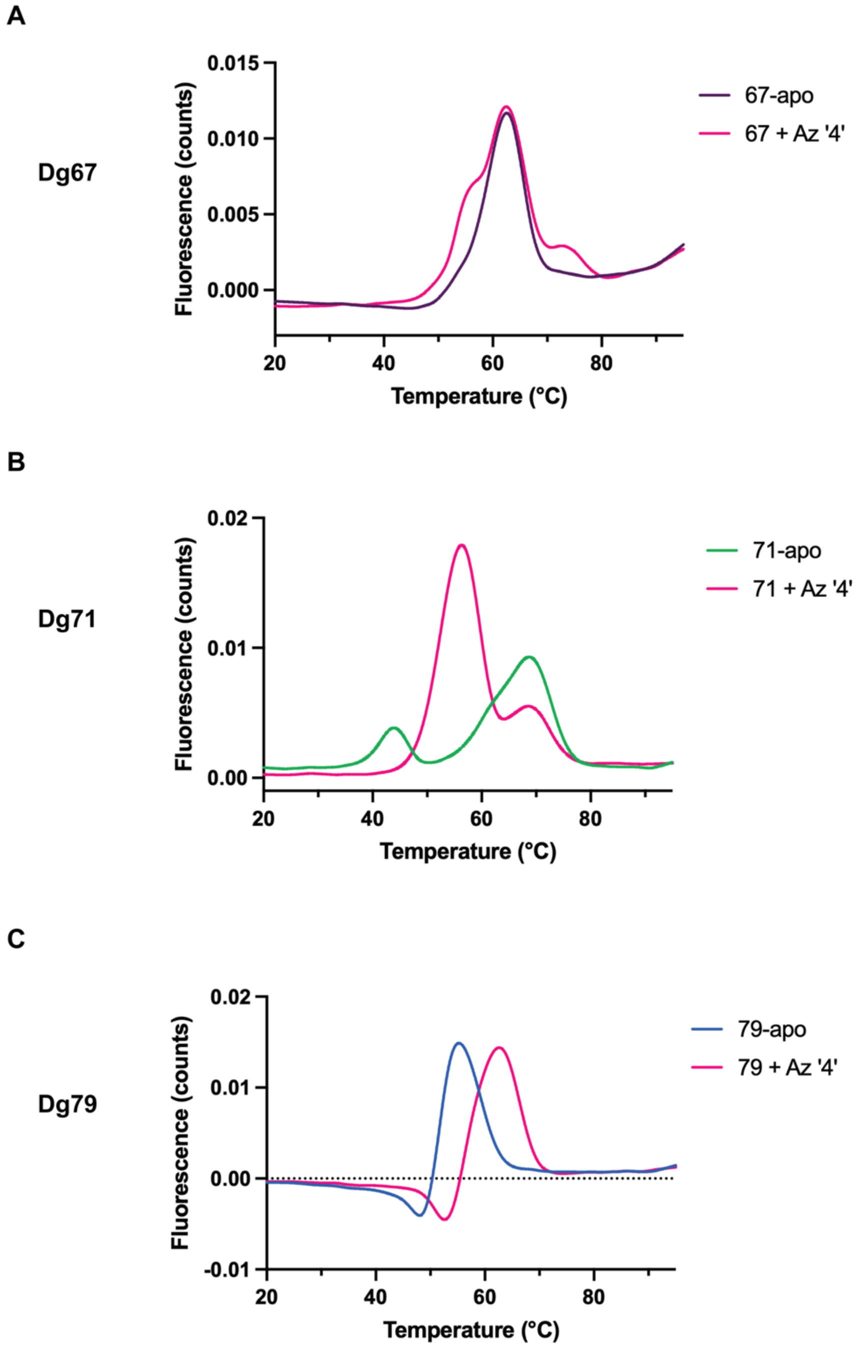

**Figure S15: Thermal profiling of GH172s with 4**

Thermal stability curves of *D. gadei* GH172 enzymes from nano differential scanning fluorimetry (nanoDSF). Dg67 (**A**), Dg71 (**B**), and Dg79 (**C**) in apo state are shown in purple, green and blue respectively. Enzymes labelled with **4** are shown in pink.

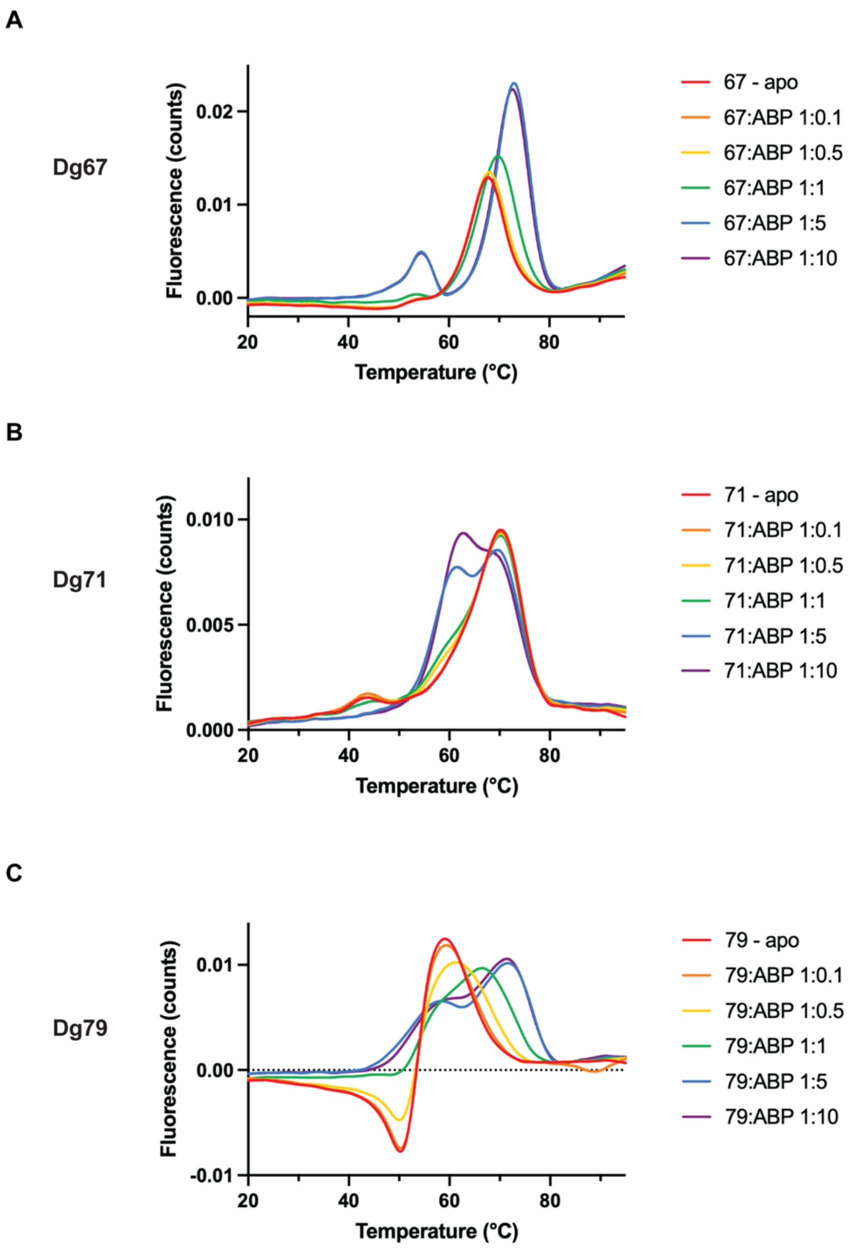

**Figure S16: NanoDSF titrations with ABP 5**

Thermal stability curves of Dg67 (**A**), Dg71 (**B**), and Dg79 (**C**) with increasing concentration of ABP ‘**5**’. Enzymes in apo state are shown in red. Enzyme:‘**5**’ molar ratios used were: 1:0.1 (orange); 1:0.5 (yellow); 1:1 (green); 1:5 (blue); and 1:10 (purple)

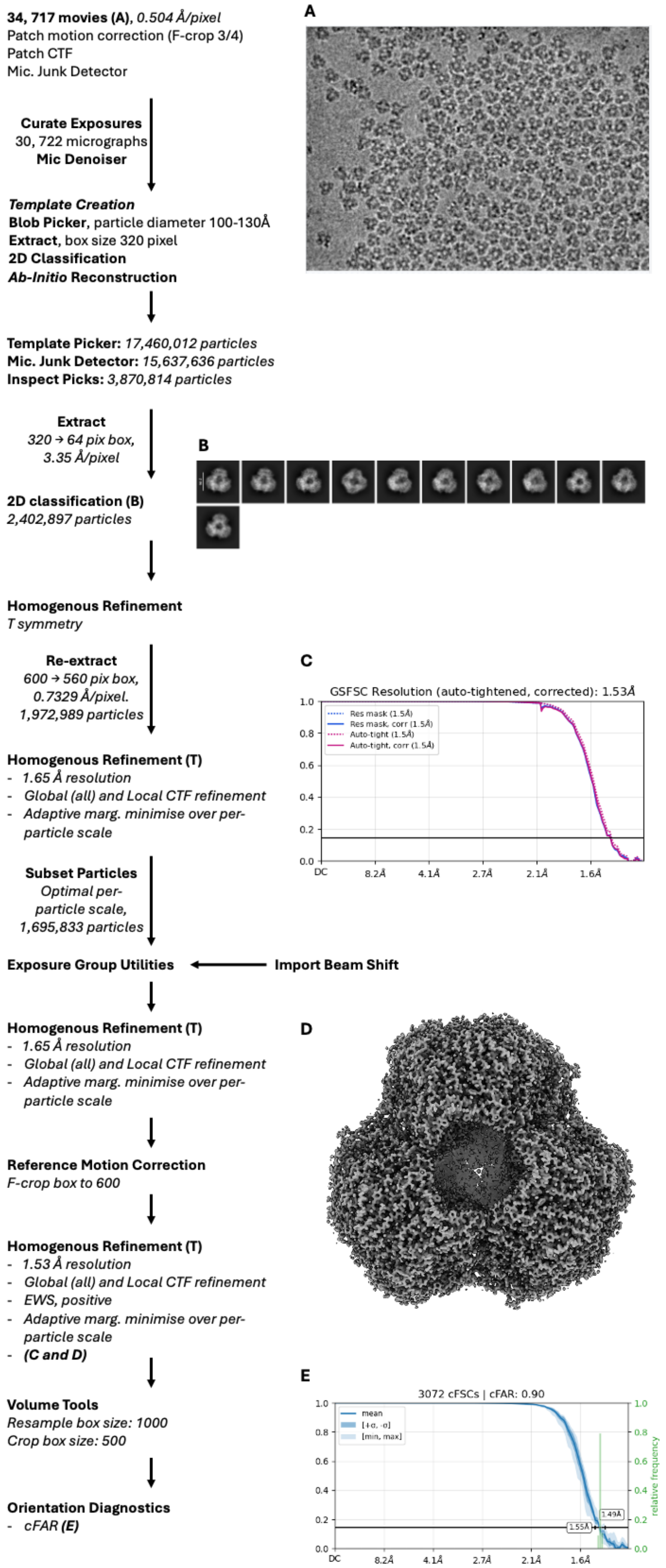

**Figure S17: CryoSPARC processing pipeline for Dg67 in complex with *4***

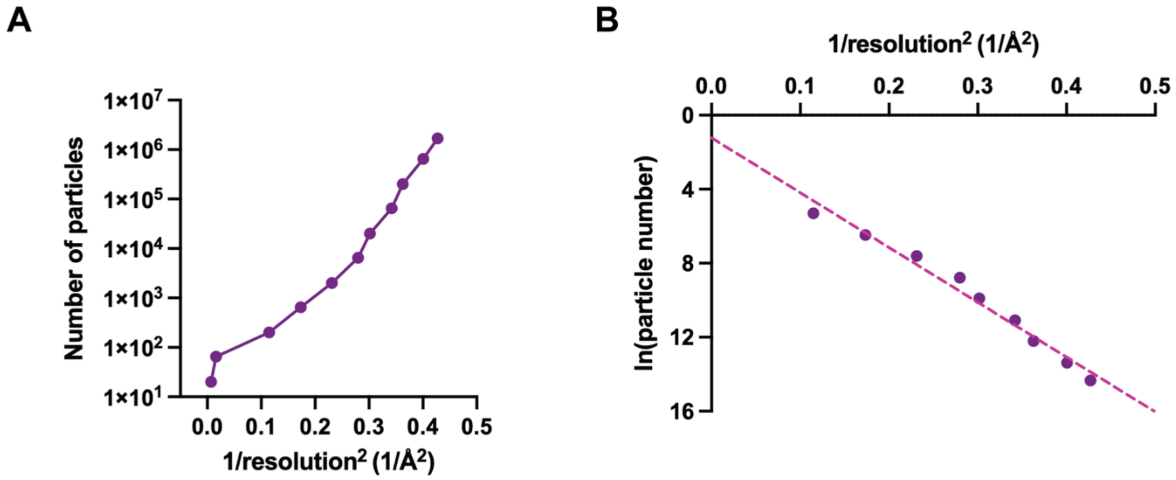

**Figure S18: Dg67 map resolution as a function of number of particles**

**A**: Data presented as discussed by Rosenthal & Henderson [18] **B**: Rosenthal-Henderson plot with a slope of 29.6 to calculate B-factor.

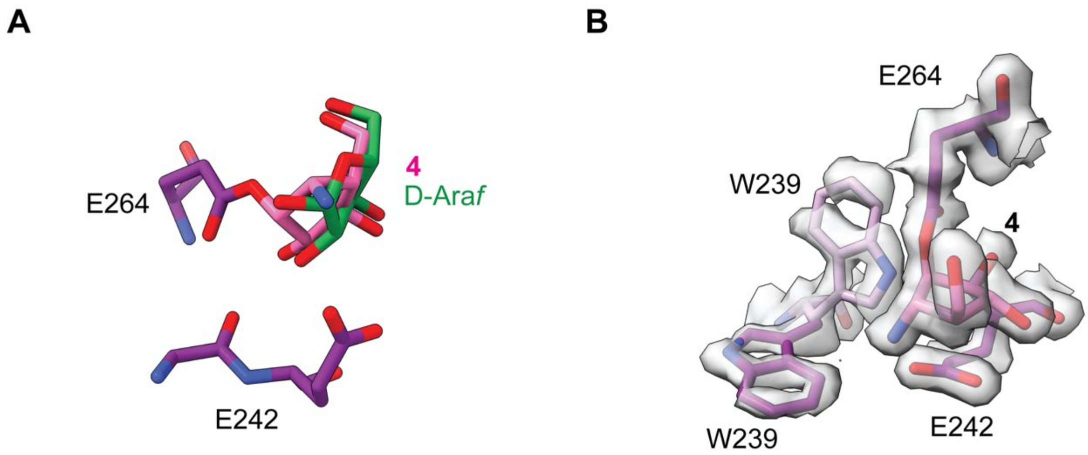

**Figure S19: Comparison of Dg67 cryo-EM structures.**

**A**: Overlay of D-Ara*f* (green) and aziridine **4** (pink) bound to catalytic residues from their respective structures. **B:** The two conformations of W239 observed in Dg67

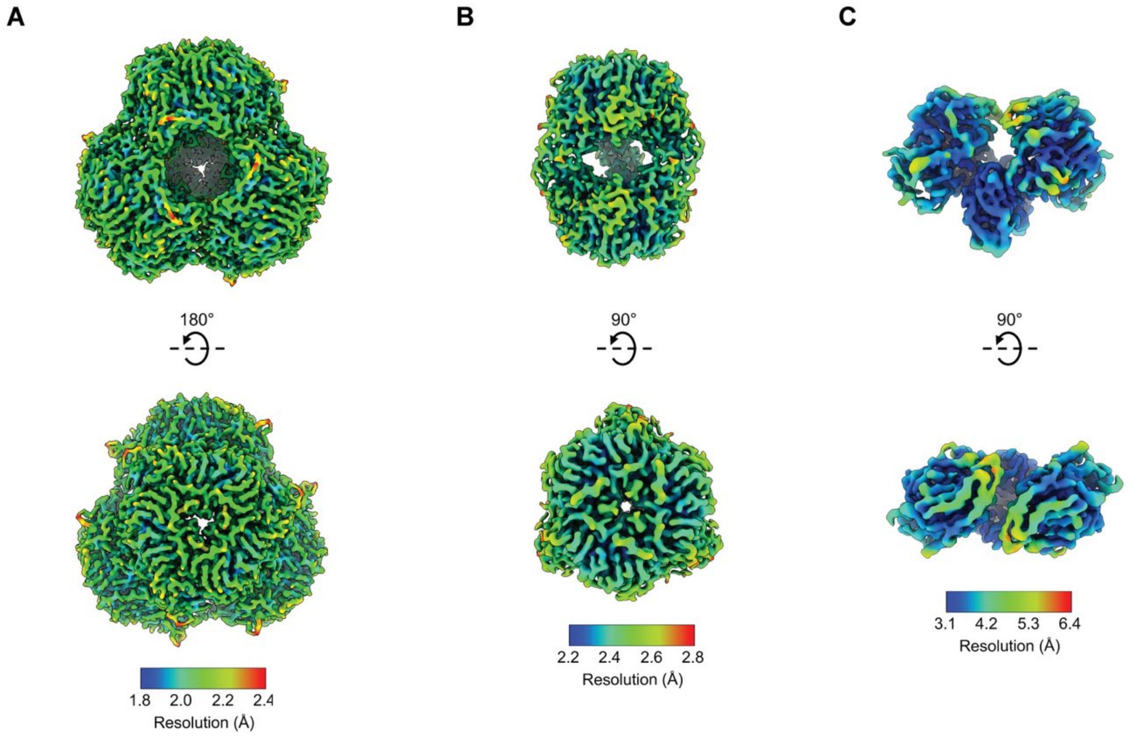

**Figure S20: Local resolution of single particle reconstructions of Dg67, Dg79 and Dg71.**

Local resolution of Dg67 (**A**), Dg79 (**B**) and Dg 71 (**C**). Colour keys for the map resolution shown below each GH172 reconstruction.

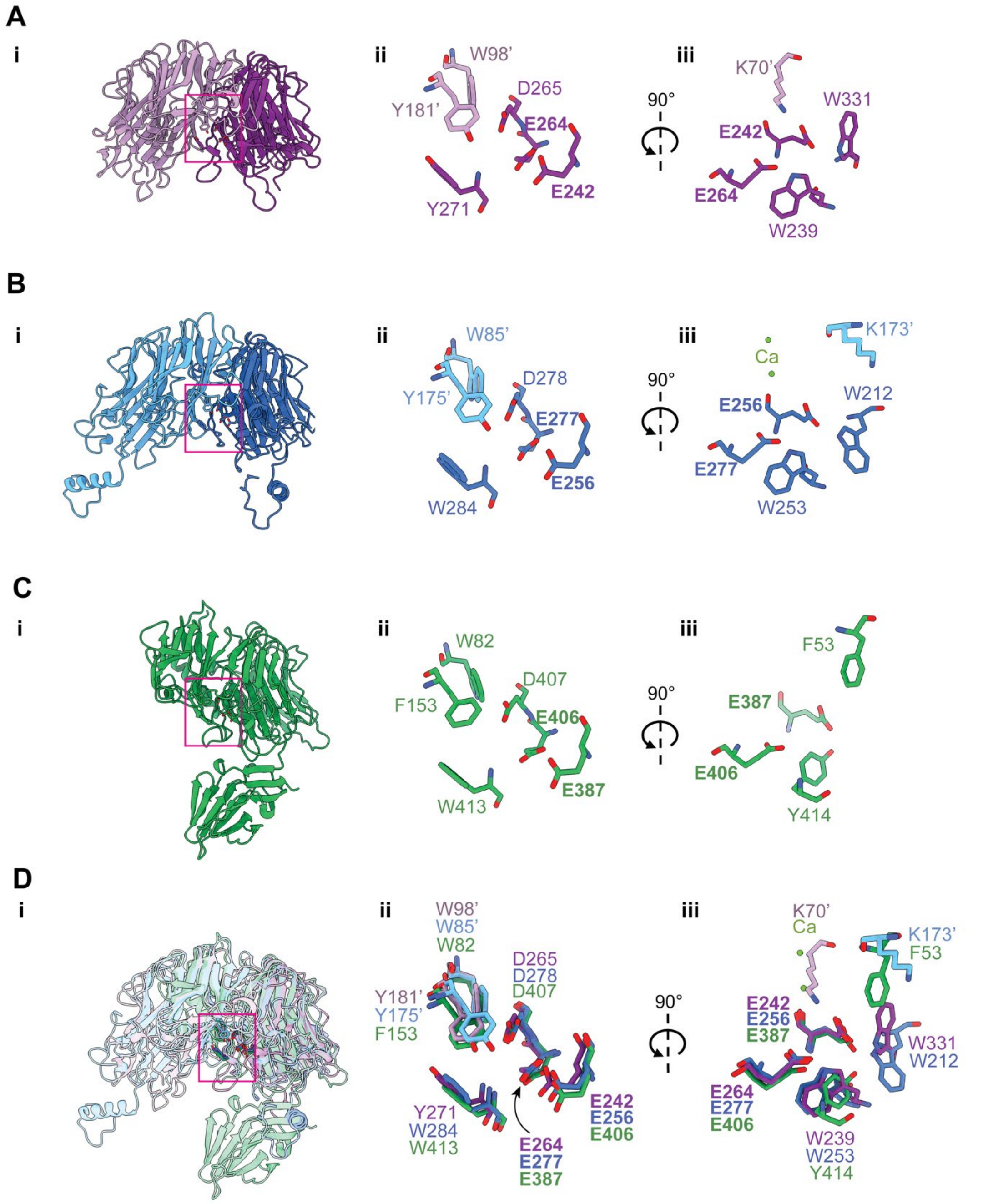

**Figure S21: Comparison of active sites of Dg67, Dg79, and Dg71 from cryo-EM structures.**

Dg_02467, Dg_02479, and Dg02471 are shown in purple (**A**), blue (**B**) and green (**C**) respectively. **Ai/Bi**, Secondary structure cartoons of dimers of Dg_02467 (**Ai**) **and** Dg_02479 (**Bi**). **Ci**, secondary structure of a monomer of Dg_02471. **Di,** overlay of **Ai**, **Bi** and **Ci**. Active sites are highlighted with pink boxes. **A/B/C/Dii**, conserved active site residues of Dg_02467, Dg_02479, and Dg02471. **A/B/C/Diii**, variable active site residues in Dg_02467, Dg_02479, and Dg02471. Catalytic residues are shown in bold.

**Figure S22: Uncropped gels for Figure 6 in this paper**

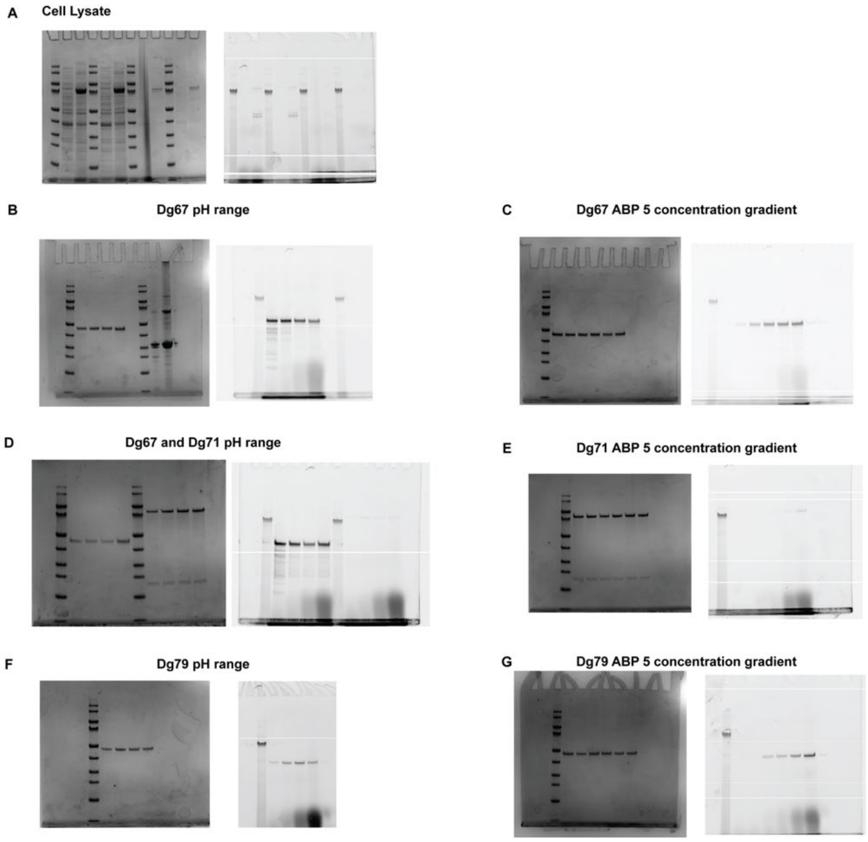

### Supplementary Tables

**Table S1: Estimated D-Ara*f* production from Dg67, Dg71, and Dg79**

| | Estimated D-Ara*f* produced (μM) | | |
|---|---|---|---|
| | Substrate **1** (1,5 linkage) | Substrate **2** (1,3 linkage) | Substrate **3** (1,3/1,5 branched) |
| Dg67 | 324 | 344 | 564 |
| Dg71 | n/a | 14 | 0.9 |
| Dg79 | 327 | 93 | 130 |

**Table S2: Cryo-EM data collection and analysis of Dg71 and Dg79**

| **Data collection** | **Dg71** | **Dg79** |
|---|---|---|
| Date | 20/06/2022 | 08/07/2025 |
| Microscope | York Glacios | York Glacios |
| Detector | Falcon-IV | Falcon-IV |
| Acquisition mode | AFIS | AFIS |
| Pixel size | 0.57 | 0.57 |
| Accelerating voltage | 200 | 200 |
| Spherical aberration | 2.7 | 2.7 |
| Total dose | 50 | 50 |
| Defocus range | -2.0 to -0.6 in 0.2 increments | -1.6 to -0.6 in 0.2 increments |
| Micrographs recorded | 3,500 | 1,000 |
| **Data processing** | | |
| Software | CryoSPARC 4.7 | CryoSPARC 4.7 |
| Alignment Software | Patch motion corr | Patch motion corr |
| Dose weighting | yes | yes |
| CFT fitting | Patch CTF | Patch CTF |
| Correction | Full | Full |
| Curated movies | 3,182 | 945 |
| Particle picking software | Blob picker | Blob picker |
| Particle picking method | Circular blobs/templates | Circular blobs/templates |
| Particles picked | 2,109,239 | 542,777 |
| Particles in final 2D classes | 180,171 | 225,007 |
| Initial model generation method | Ab initio C1 | Ab initio C1 |
| Initial model generation software | CryoSPARC 4.7 | CryoSPARC 4.7 |
| Particles in final 3D reconstruction | 59,433 | 192,387 |
| Alignment software | CryoSPARC 4.7 | CryoSPARC 4.7 |
| Reconstruction software | CryoSPARC 4.7 | CryoSPARC 4.7 |
| Box size pixels | 500 (F-crop 250) | 512 |
| Box size (Å) | 287 | 287 |
| Voxel size (Å) | 1.148 | 0.56 |
| Symmetry | C2 | D3 |
| Map resolution (GC-FSC 0.143) (Ã) | 3.6 | 2.5 |
| Sharpening B-factor (Å2) | 129 | 98 |
| **Depositions** | | |
| EMDB | tbc | tbc |
| EMPIAR | n/a | n/a |

**Table S3: Model building and refinement statistics for Dg71 and Dg79**

| Model building and refinement | Dg71 | Dg79 |
|---|---|---|
| Software | Phenix.refine | Phenix.refine |
| Refinement algorithm | Real space | Real space |
| Resolution cutoff (Å) | 3.6 | 2.5 |
| FSC model-vs-map = 0.5 (Å) | 4.0 | 2.7 |
| **Model** | | |
| Number of amino acids | 609 | 356 |
| Waters | 0 | 46 |
| Ligands | 0 | 3 |
| Bond length outliers | 0 | 0 |
| Bond angle outliers | 1 | 0 |
| **RMS deviations** | | |
| Bonds (Å) | 0.004 | 0.002 |
| Angles (˚) | 0.795 | 0.460 |
| **Validation** | | |
| Molprobity score | 1.69 | 1.13 |
| Clash score | 10.27 | 3.38 |
| Rotamer outliers (%) | 1.78 | 0.33 |
| Cb outliers (%) | n/a | n/a |
| CaBLAM outliers (%) | 3.41 | 3.78 |
| **Ramachandran** | | |
| Favoured | 89.18 | 98.57 |
| Outlier | 0.07 | 0 |
| Model vs Data CC (mask) | 0.85 | 0.92 |
| **B-factors** | | |
| Protein | 102.66 | 94.02 |
| Waters | 0 | 97.52 |
| Ligands | 0 | 130.42 |
| **Map viewing levels** | | |
| Map | 0.06 | 0.03 |
| Sharp | 0.06 | 0.03 |
| A | 0.075 | 0.07 |
| B | 0.075 | 0.07 |
| Mask | 0.9 | 0.9 |
| **Depositions** | | |
| PDB ID | tbc | tbc |

**Table S4: Data collection and refinement statistics for the crystal structure of Dg79**

| | **Dg79** |
|---|---|
| **Date** | 12/08/2020 |
| **Source** | I24 |
| **Wavelength (Å)** | 0.9786 |
| **Space group** | H 3 2 |
| ***a, b, c*** **(Å)** | 89.15, 89.15, 289.32 |
| **α, β, γ (°)** | 90, 90, 120 |
| **No. of measured reflections** | 895646 (28001) |
| **No. of independent reflections** | 87401 (4237) |
| **Resolution (Å)** | 68.11 - 1.40 (1.42 - 1.40) |
| **$CC_{1/2}$** | 0.999 (0.412) |
| ***I/σI*** | 12.7 (0.6) |
| **Completeness (%)** | 99.9 (99.0) |
| **Redundancy** | 10.2 (6.6) |
| **$R_{work/}$ $R_{free}$** | 14.40/18.30 |
| **Protein** | 3052 |
| **Ligand/Ions** | 3 |
| **Water** | 205 |
| **Protein** | 25.6 |
| **Ligand/Ions** | 27.8 |
| **Water** | 35.6 |
| **Bond lengths (Å)** | 0.0194 |
| **Bond angles (°)** | 2.1850 |
| **Ramachandran plot (%) Favoured/Outliers** | 97.3 / 0 |
| **PDB** | 8AH3 |

**Table S5: Cryo-EM data collection and analysis of Dg67**

| Data collection | D67-apo | Dg67-Ara | Dg67-A4 |
|---|---|---|---|
| Date | 20/06/2022 | 08/07/2025 | 17/04/2025 |
| Microscope | York Glacios | York Glacios | eBIC Krios mI |
| Detector | Falcon-IV | Falcon-IV | Gatan K3 |
| Acquisition mode | AFIS | AFIS | AFIS |
| Pixel size | 0.57 | 0.57 | 0.504 |
| Accelerating voltage | 200 | 200 | 300 |
| Spherical aberration | 2.7 | 2.7 | 2.7 |
| Total dose | 50 | 50 | 40.31 |
| Defocus range | - 1.6 to -0.6 in 0.2 increments | -1.6 to -0.6 in 0.2 increments | 1.6 to -0.6 in 0.2 increments |
| Micrographs recorded | 5,040 | 1,247 | 34,717 |
| **Data processing** | | | |
| Software | CryoSPARC 4.7 | CryoSPARC 4.7 | CryoSPARC 4.7 |
| Alignment Software | Patch motion corr | Patch motion corr | Patch motion corr |
| Dose weighting | yes | yes | yes |
| CFT fitting | Patch CTF | Patch CTF | Patch CTF |
| Correction | Full | Full | Full |
| Curated movies | 4,662 | 1,049 | 33,617 |
| Particle picking software | Blob picker | Blob picker | Blob picker |
| Particle picking method | Circular blobs/templates | Circular blobs/templates | Circular blobs/templates |
| Particles picked | 2,949,475 | 621,472 | 19,547,825 |
| Particles in final 2D classes | 236,124 | 75,192 | 3,095,166 |
| Initial model generation method | Ab initio C1 | Ab initio C1 | Ab initio C1 |
| Initial model generation software | CryoSPARC 4.7 | CryoSPARC 4.7 | CryoSPARC 4.7 |
| Particles in final 3D reconstruction | 121,392 | 47,664 | 1,695,196 |
| Alignment software | CryoSPARC 4.7 | CryoSPARC 4.7 | CryoSPARC 4.7 |
| Reconstruction software | CryoSPARC 4.7 | CryoSPARC 4.7 | CryoSPARC 4.7 |
| Box size pixels | 600 | 600 | 1000 (crop 500) |
| Box size (Å) | 342 | 342 | 453 |
| Voxel size (Å) | 0.57 | 0.57 | 0.41 |
| Symmetry | T | T | T |
| Map resolution (GC-FSC 0.143) (Ã) | 2.1 | 2.6 | 1.5 |
| Sharpening B-factor (Å2) | 57 | 92 | 59 |
| **Depositions** | | | |
| EMDB | tbc | tbc | tbc |
| EMPIAR | n/a | n/a | tbc |

**Table S6: Model building and refinement statistics for Dg67**

| Model building and refinement | Dg02467-apo | Dg02467-Ara | Dg 02467-Az |
|---|---|---|---|
| Software | Phenix.refine | Phenix.refine | Phenix.refine |
| Refinement algorithm | Real space | Real space | Real space |
| Resolution cutoff (Å) | 2.1 | 2.6 | 1.5 |
| FSC model-vs-map = 0.5 (Å) | 2.2 | 2.8 | 1.6 |
| **Model** | | | |
| Number of amino acids | 336 | 336 | 336 |
| Waters | 50 | 0 | 62 |
| Ligands | 0 | 4 | 4 |
| Bond length outliers | 0 | 0 | 0 |
| Bond angle outliers | 0 | 0 | 0 |
| **RMS deviations** | | | |
| Bonds (Å) | 0.002 | 0.003 | 0.002 |
| Angles (˚) | 0.532 | 0.550 | 0.499 |
| **Validation** | | | |
| Molprobity score | 1.35 | 1.40 | 1.30 |
| Clash score | 3 | 3 | 2.8 |
| Rotamer outliers (%) | 0 | 0 | 0 |
| Cb outliers (%) | n/a | n/a | n/a |
| CaBLAM outliers (%) | 4 | 4 | 4 |
| **Ramachandran** | | | |
| Favoured | 96 | 96 | 97 |
| Outlier | 0.27 | 0.27 | 0.27 |
| Model vs Data CC (mask) | 0.94 | 0.9 | 0.96 |
| **B-factors** | | | |
| Protein | 66 | 95 | 62 |
| Waters | 66 | 0 | 60 |
| Ligands | 70 | 110 | 58 |
| **Map viewing levels** | | | |
| Map | 0.03 | 0.03 | 0.03 |
| Sharp | 0.03 | 0.03 | 0.03 |
| A | 0.07 | 0.07 | 0.07 |
| B | 0.07 | 0.07 | 0.07 |
| Mask | 0.9 | 0.9 | 0.9 |
| **Depositions** | | | |
| PDB ID | tbc | tbc | tbc |

**Table S7: Nano-DSF protein unfolding temperatures**

| Protein | Protein only | Protein + 4 | Protein + 5 |
|---|---|---|---|
| **Dg67** | 62.5 ± 0.1 °C | 62.6 ± 0.3 °C | 72.7 ± 0.2 °C |
| **Dg71** | 70.2 ± 0.1 °C | 56.3 ± 0.0 °C | 62.8 ± 0.7 °C |
| **Dg79** | 55.4 ± 0.2 °C | 62.5 ± 0.1 °C | 71.3 ± 0.2°C |

**Table S8: PISA analysis of quaternary structures of Dg67, Dg71 and Dg79**

| Protein | Surface Area (Å$^2$) | Buried Area (Å$^2$) |
|---|---|---|
| **Dg67 (dodecamer)** | 124,080 | 85,671 |
| **Dg71 (dimer)** | 46,067 | 4,118 |
| **Dg79 (hexamer)** | 66,750 | 47,220 |

### NMR Spectra

**MH001** in $CDCl_3$

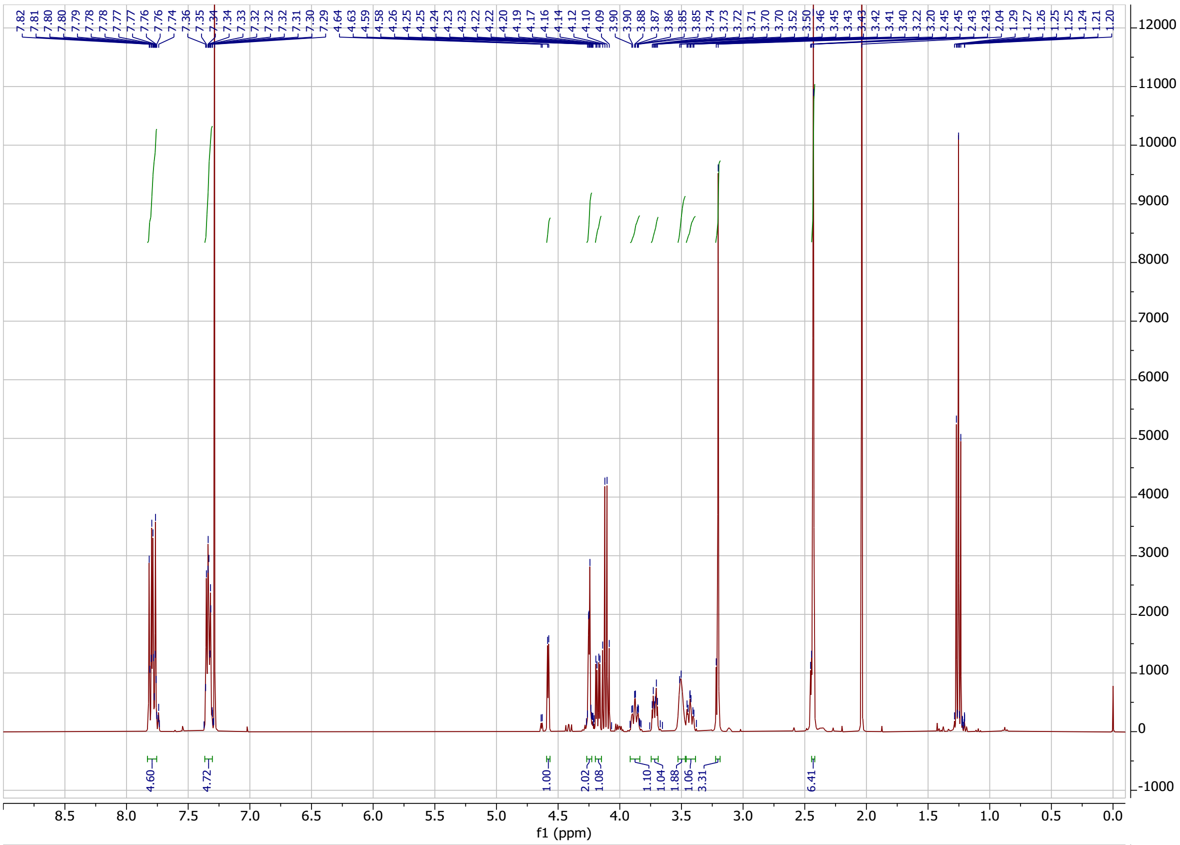

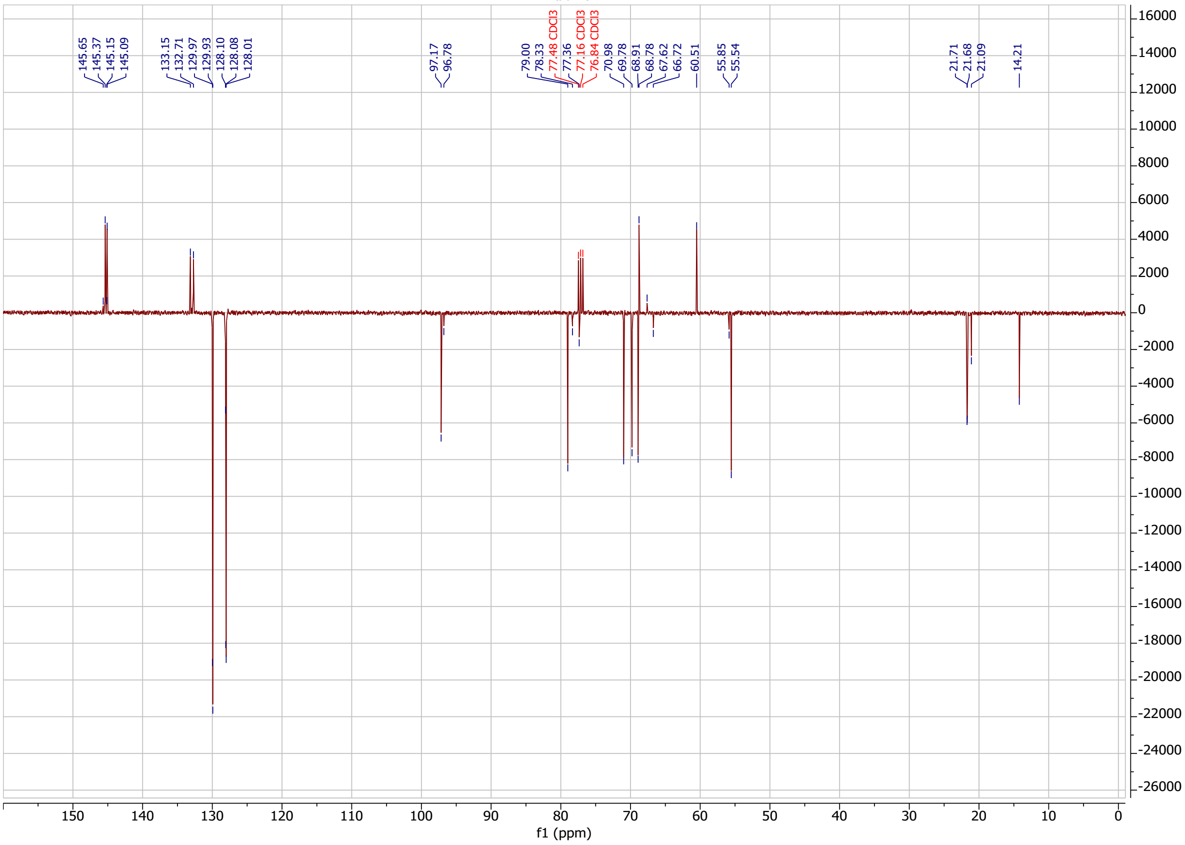

**MH009** in $CDCl_3$

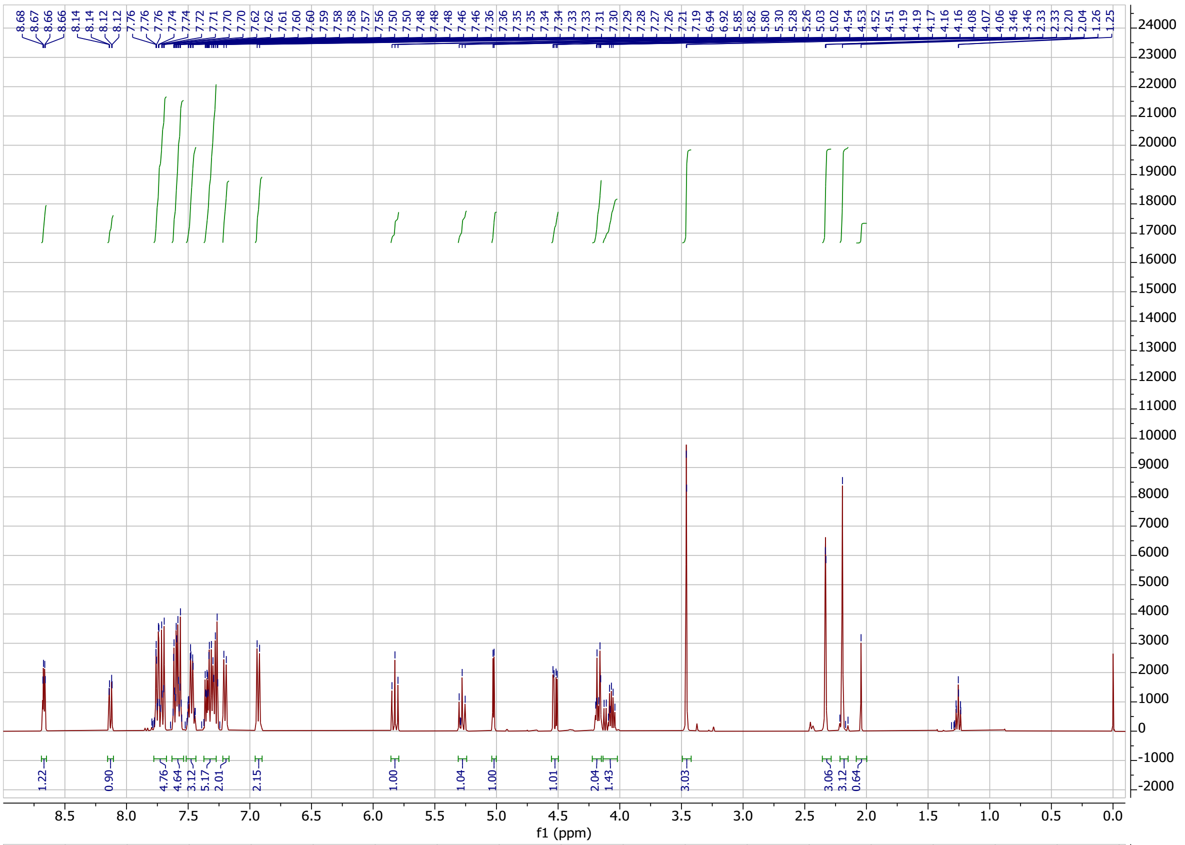

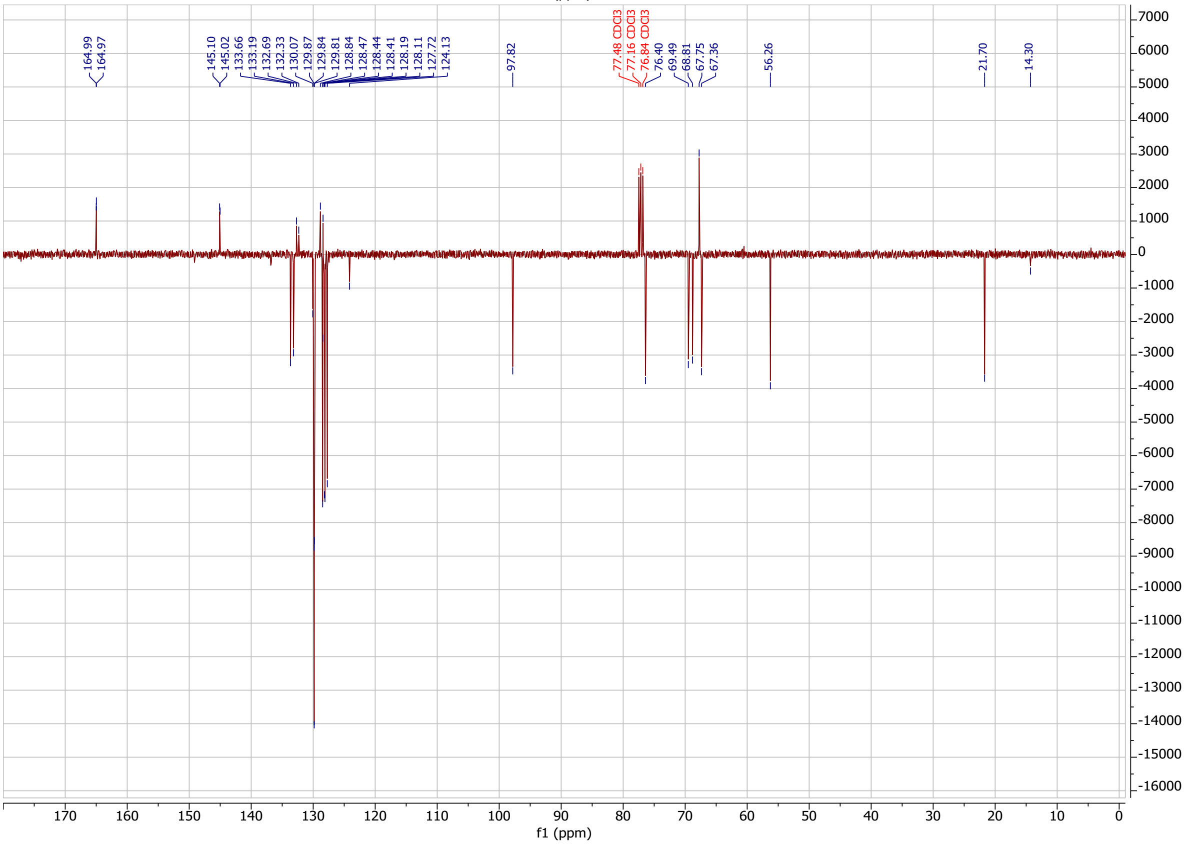

**MH008** in $CDCl_3$

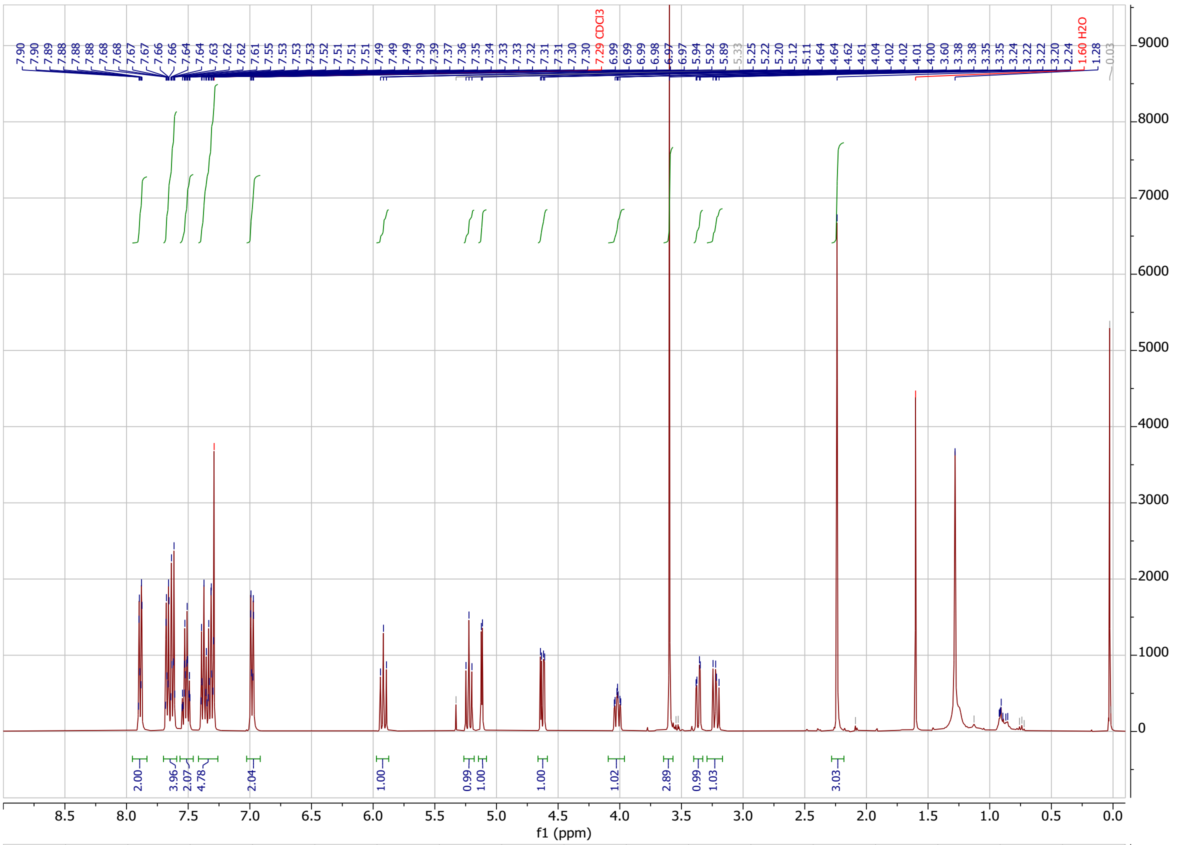

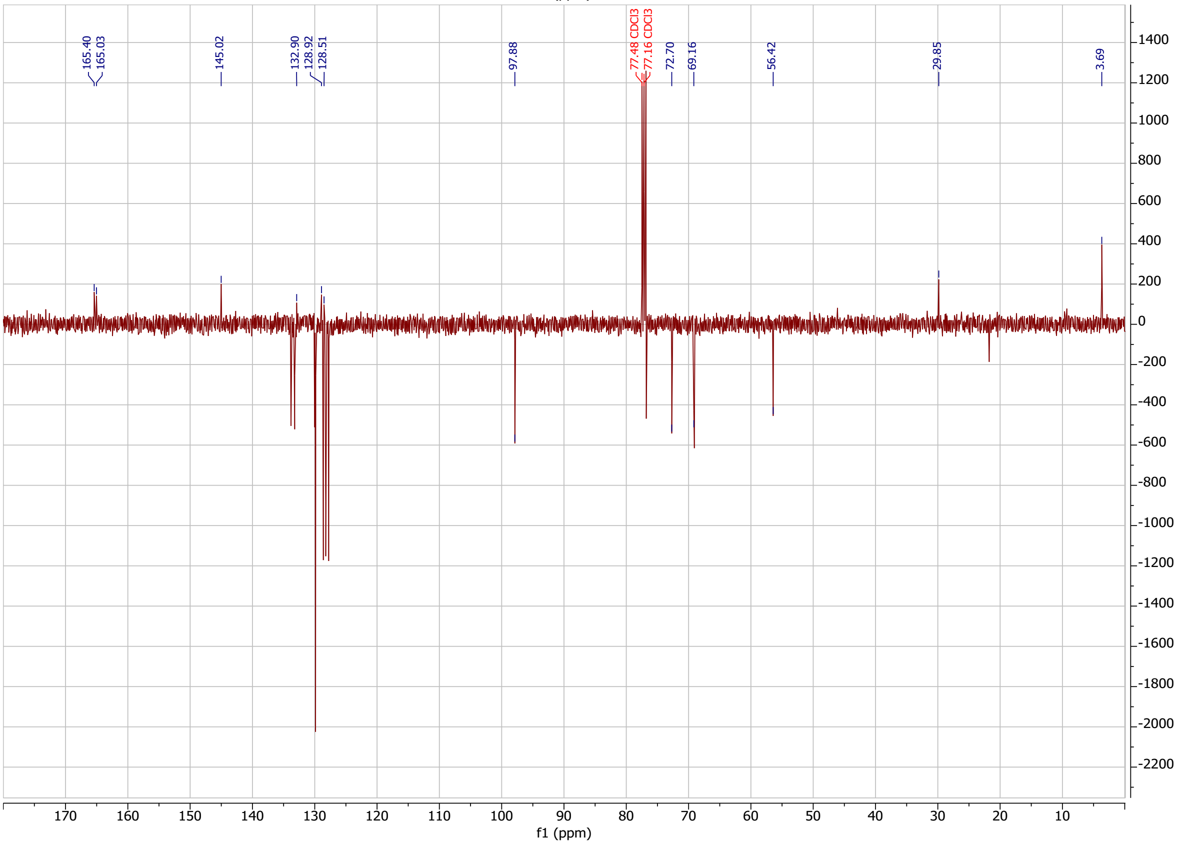

**MH066** in $CDCl_3$

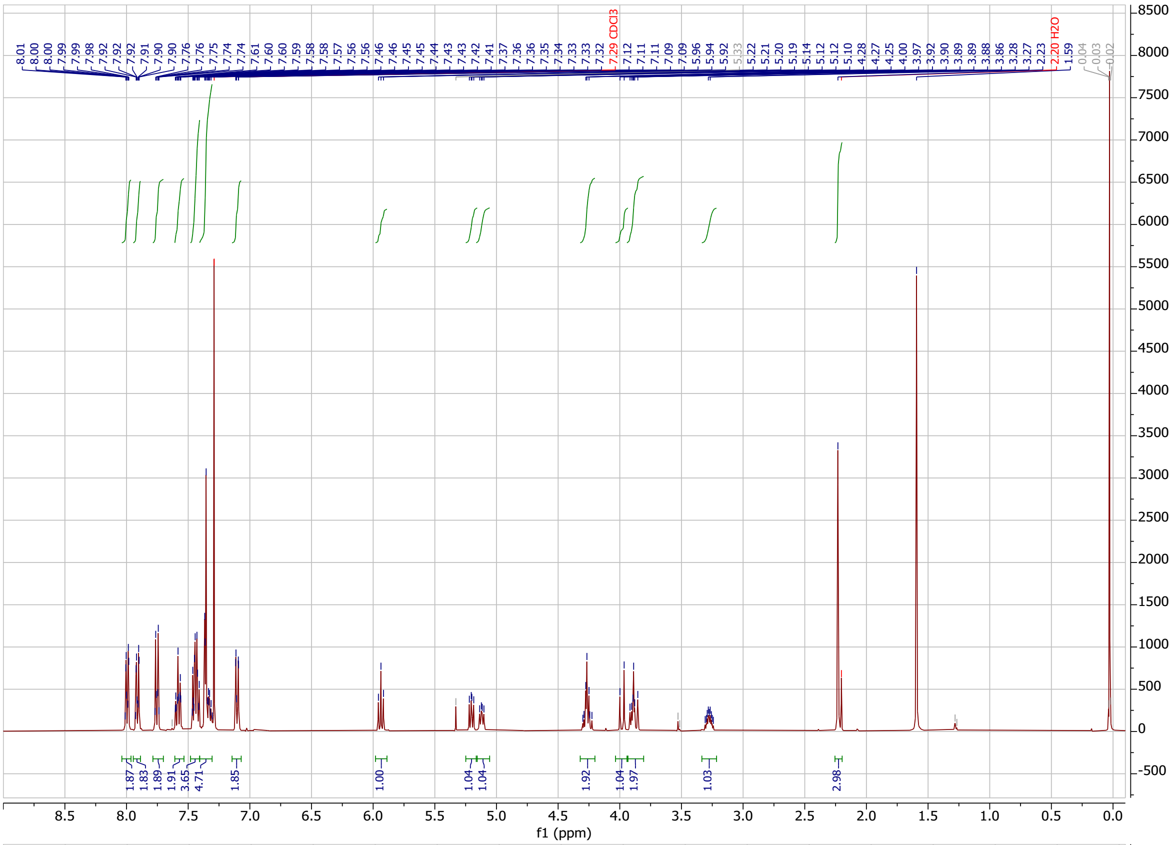

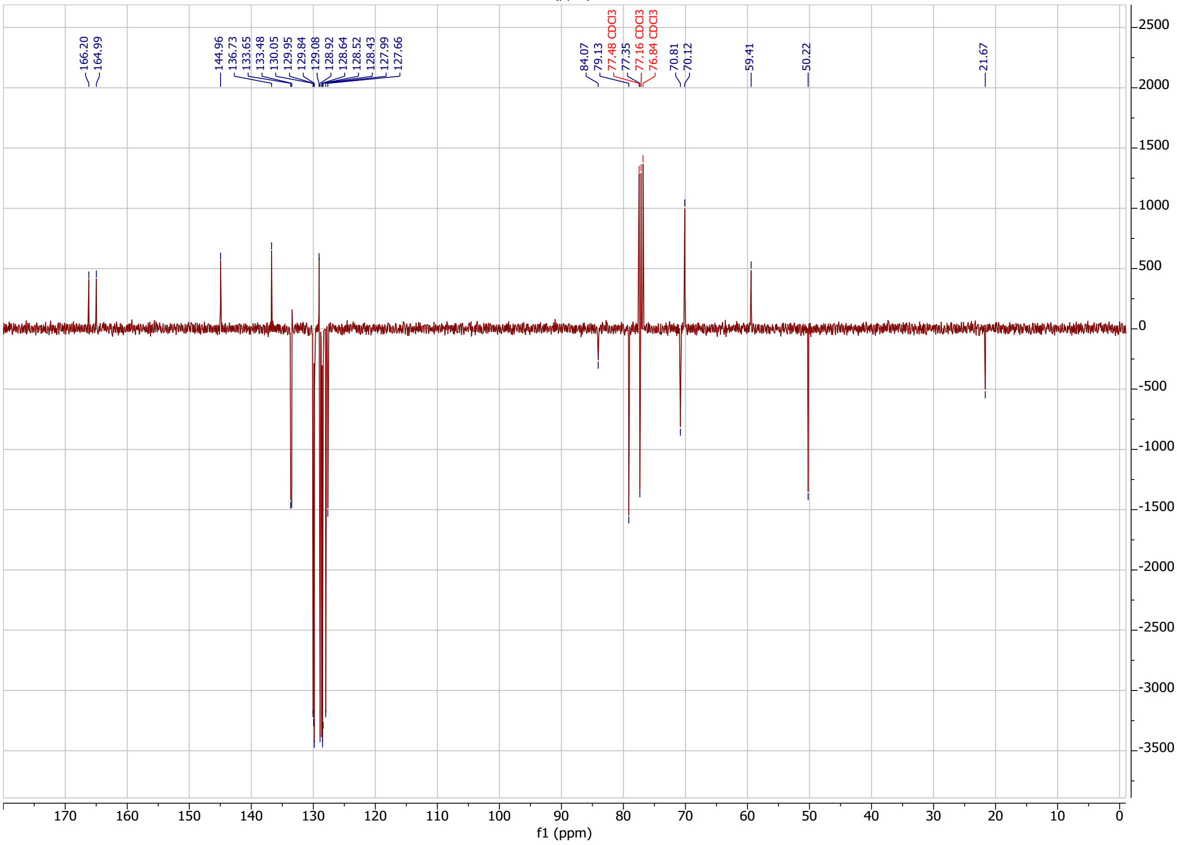

**MH068** in $CDCl_3$

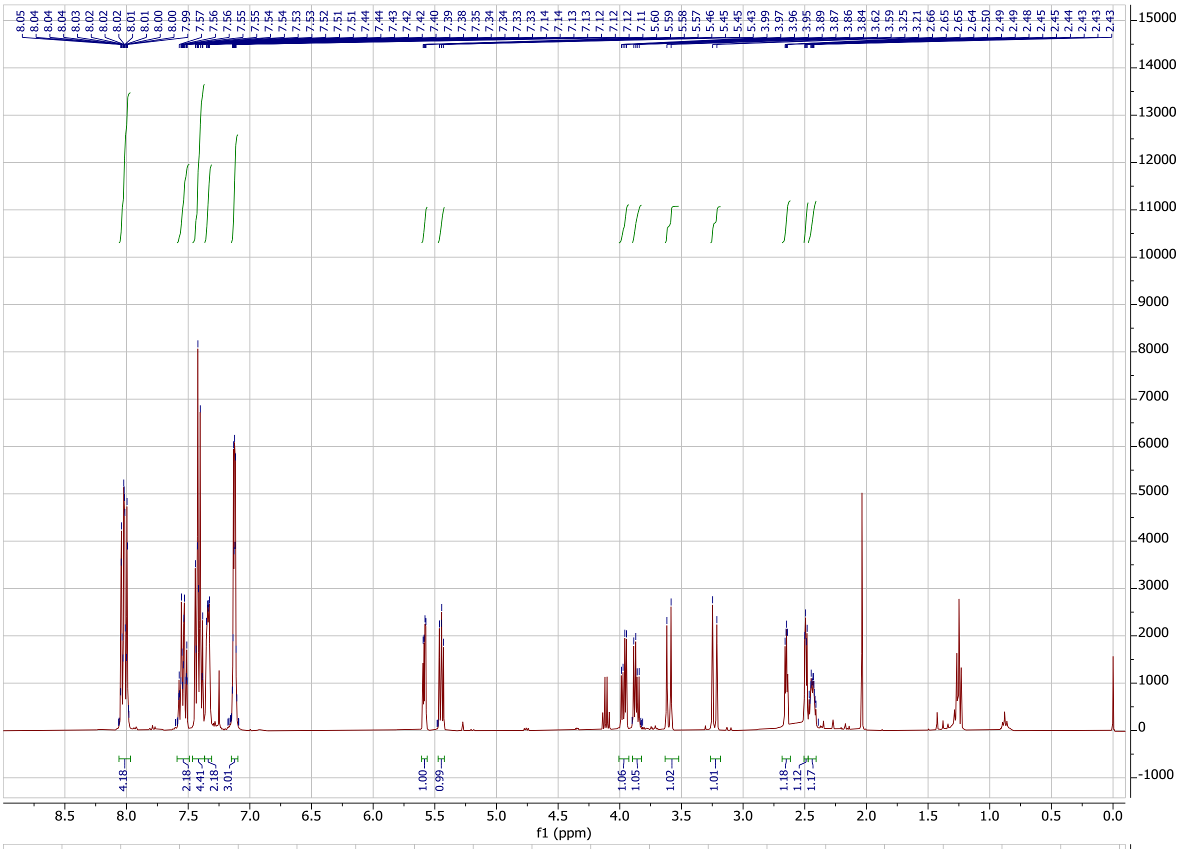

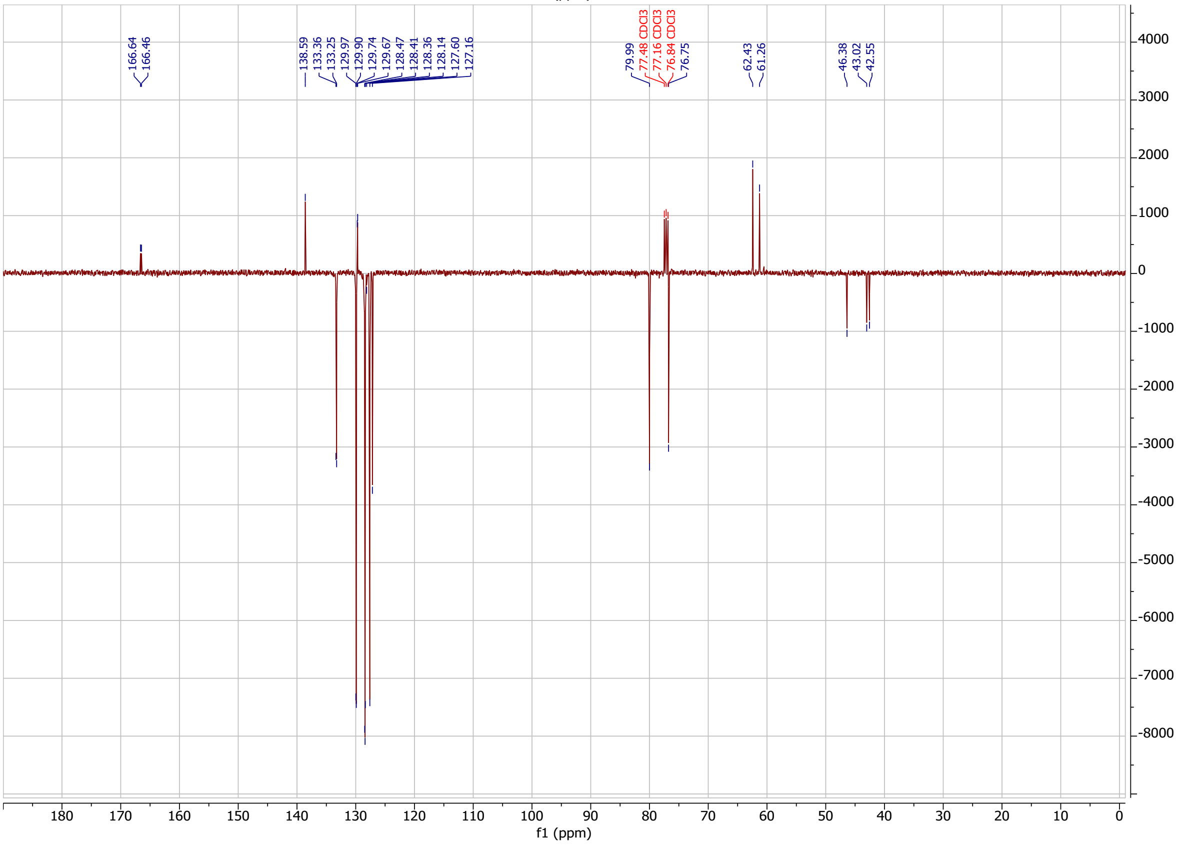

**MH072.1** in $CDCl_3$

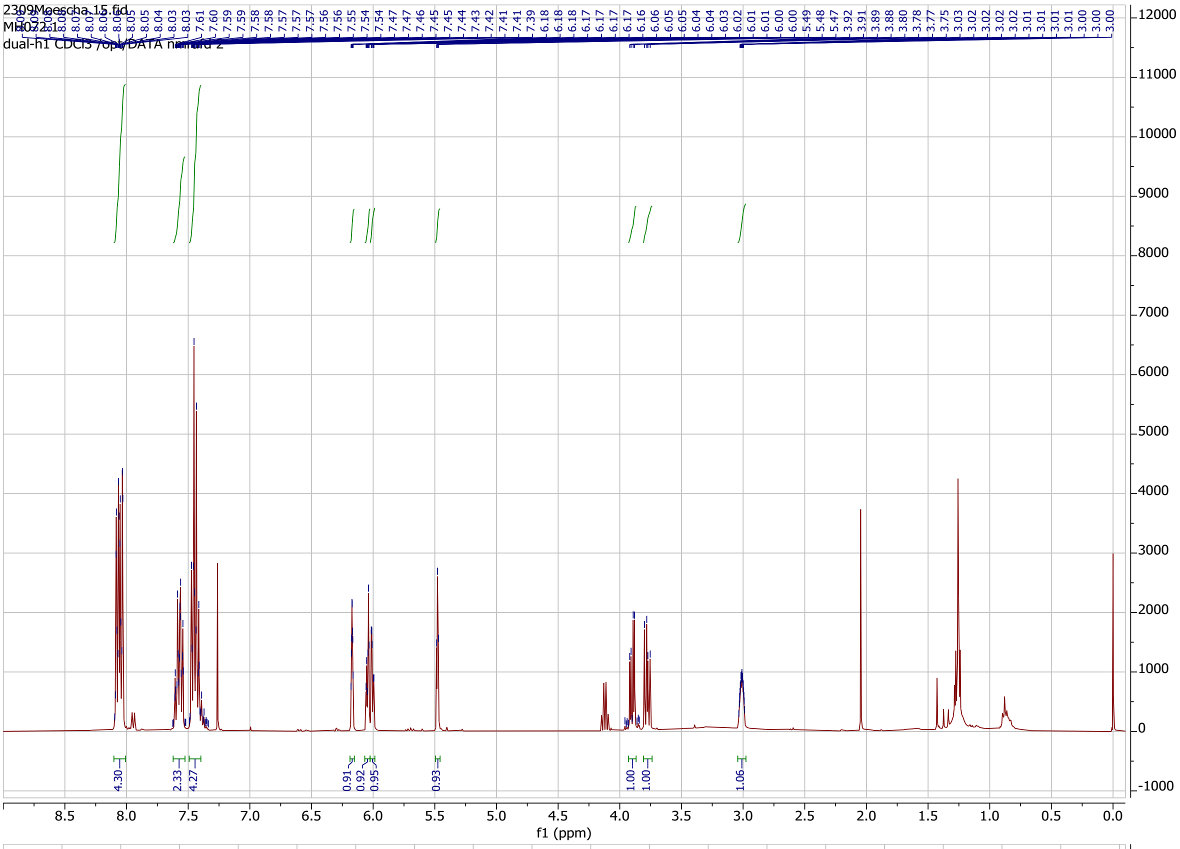

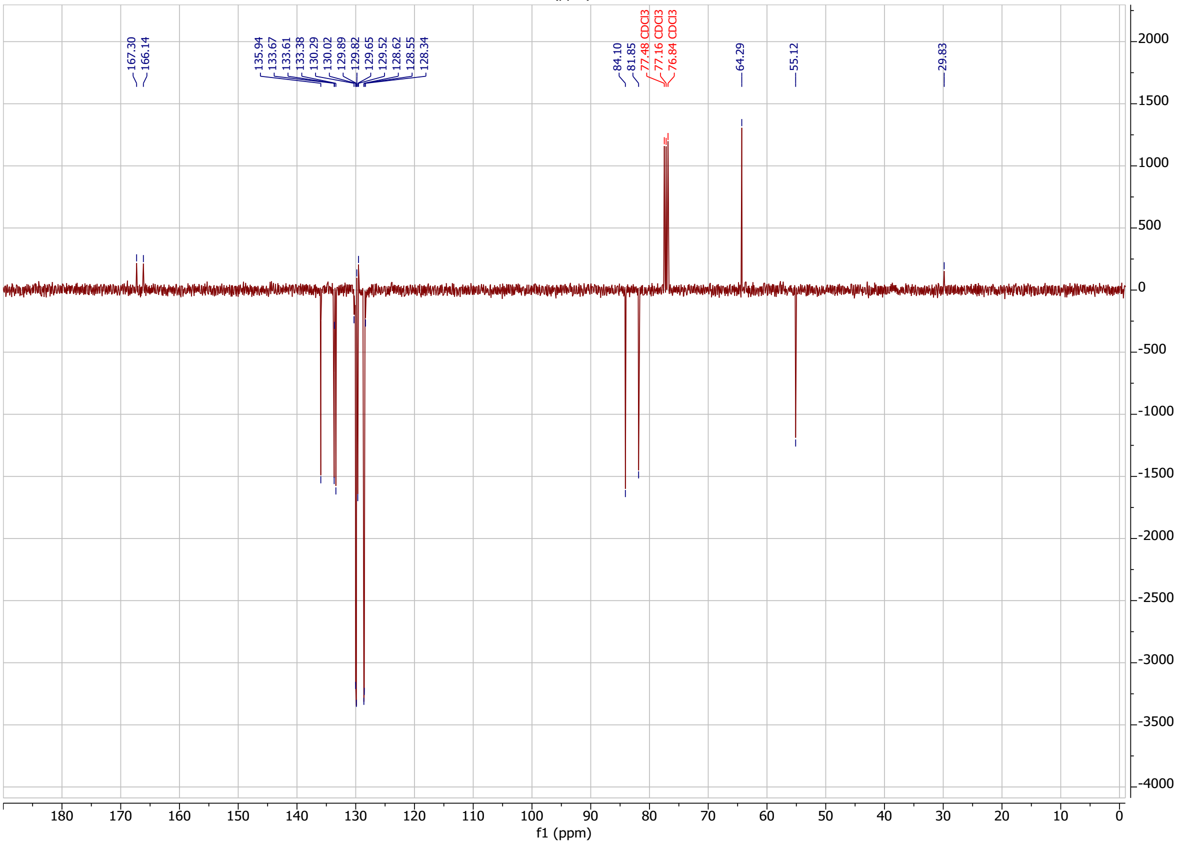

**MH172** in $CDCl_3$

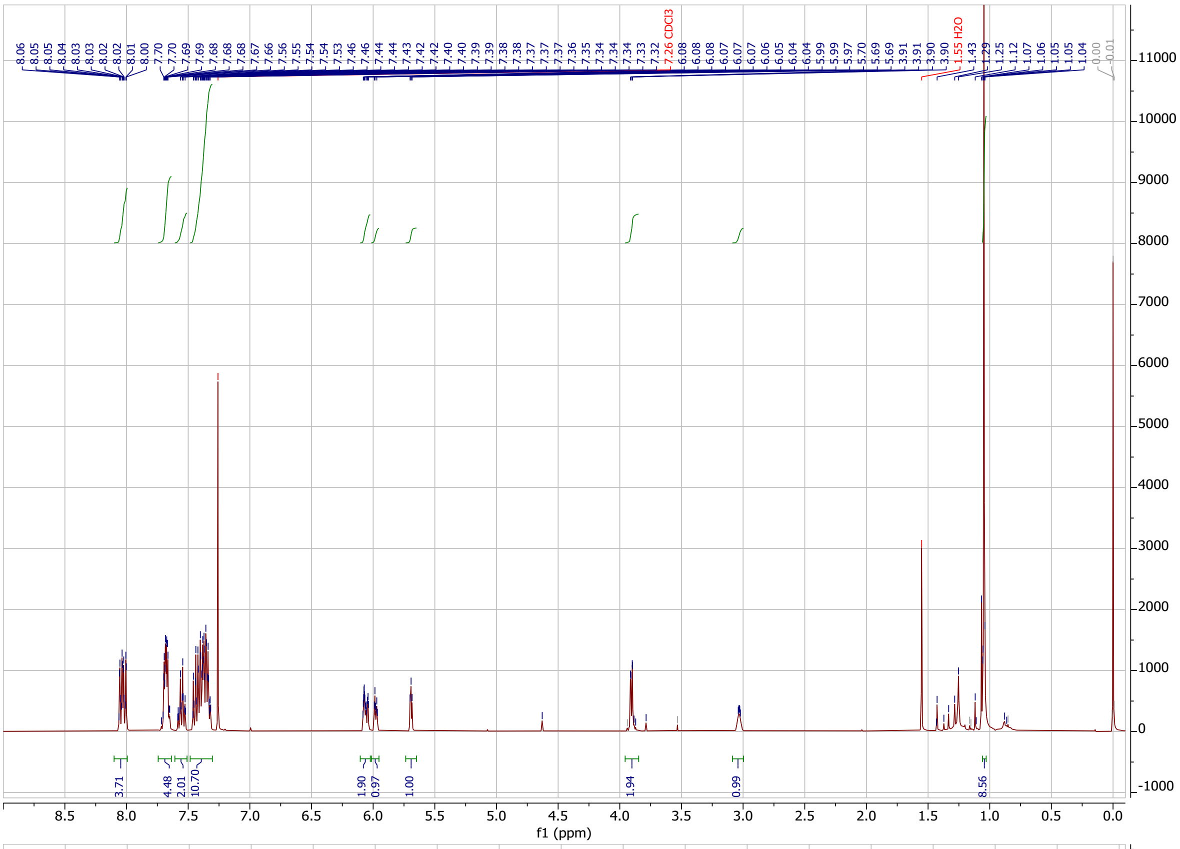

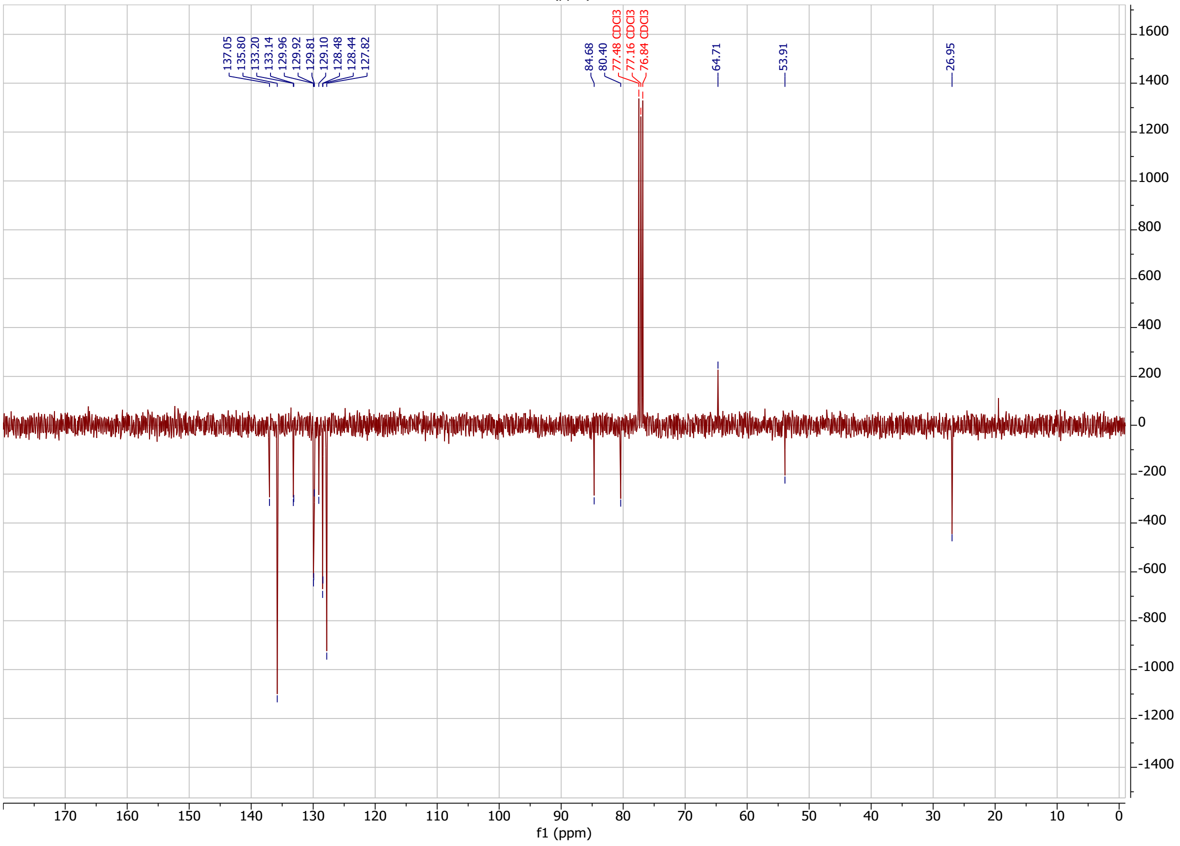

**MH174** in $CDCl_3$

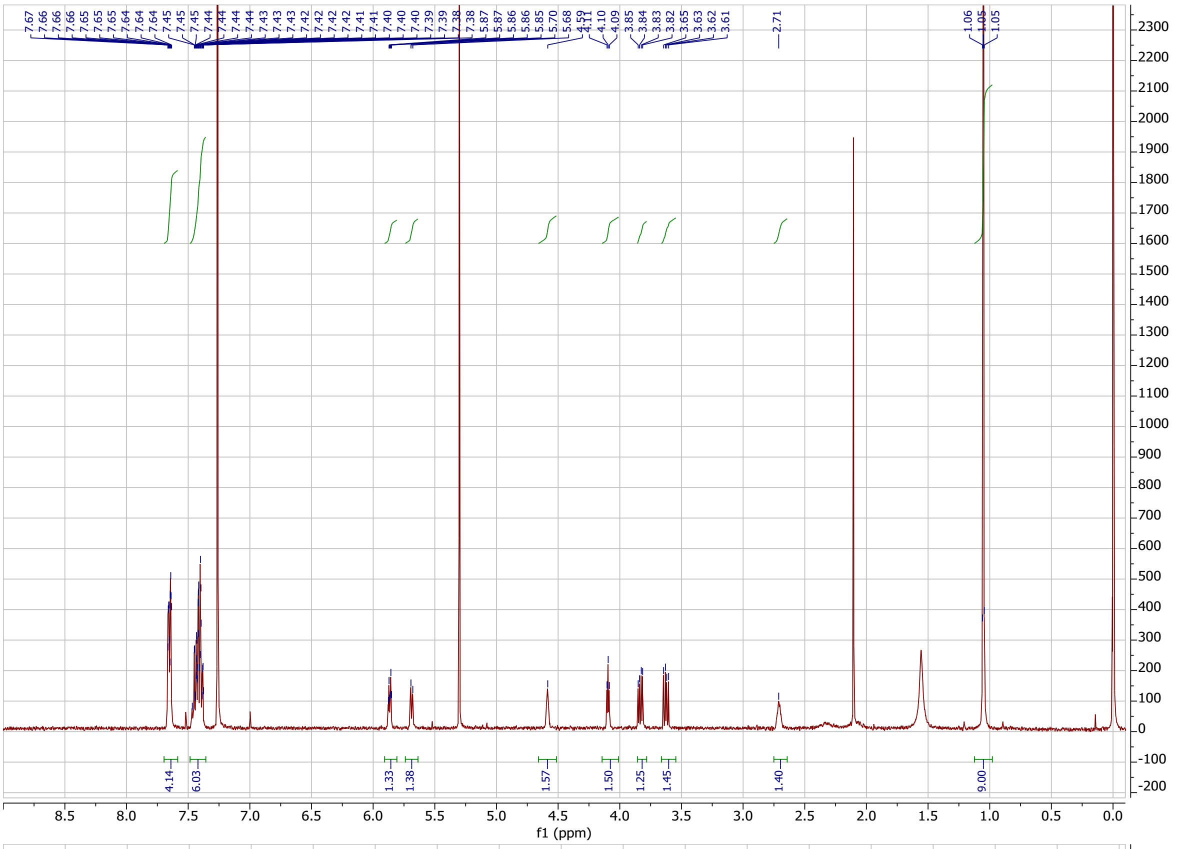

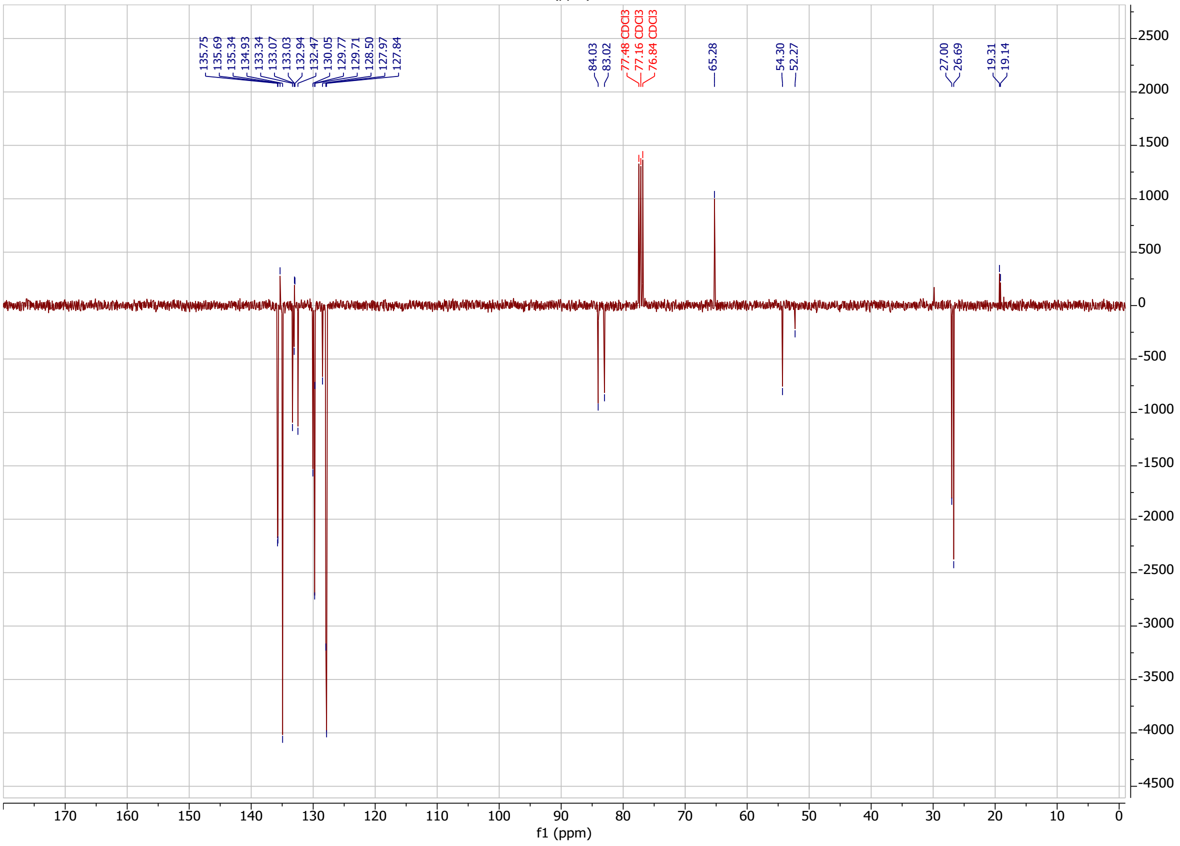

**RvB15** in $CDCl_3$

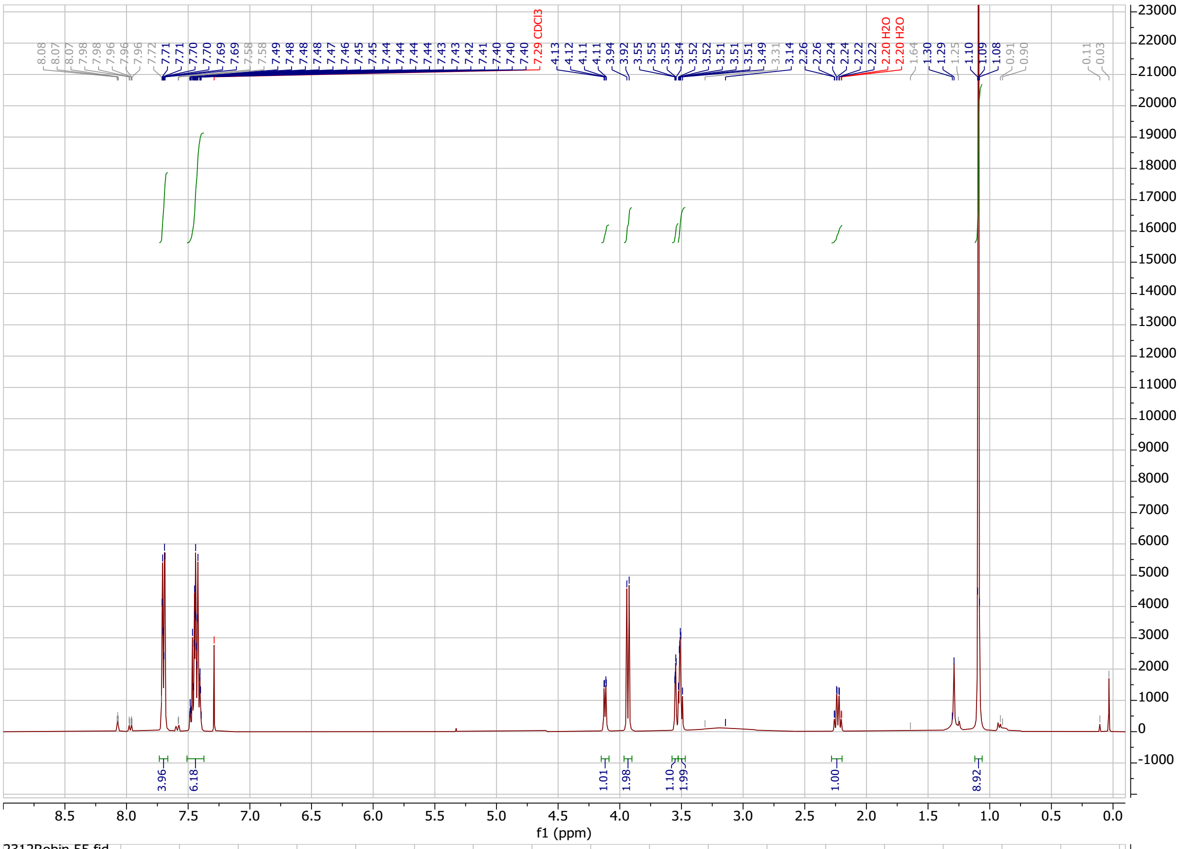

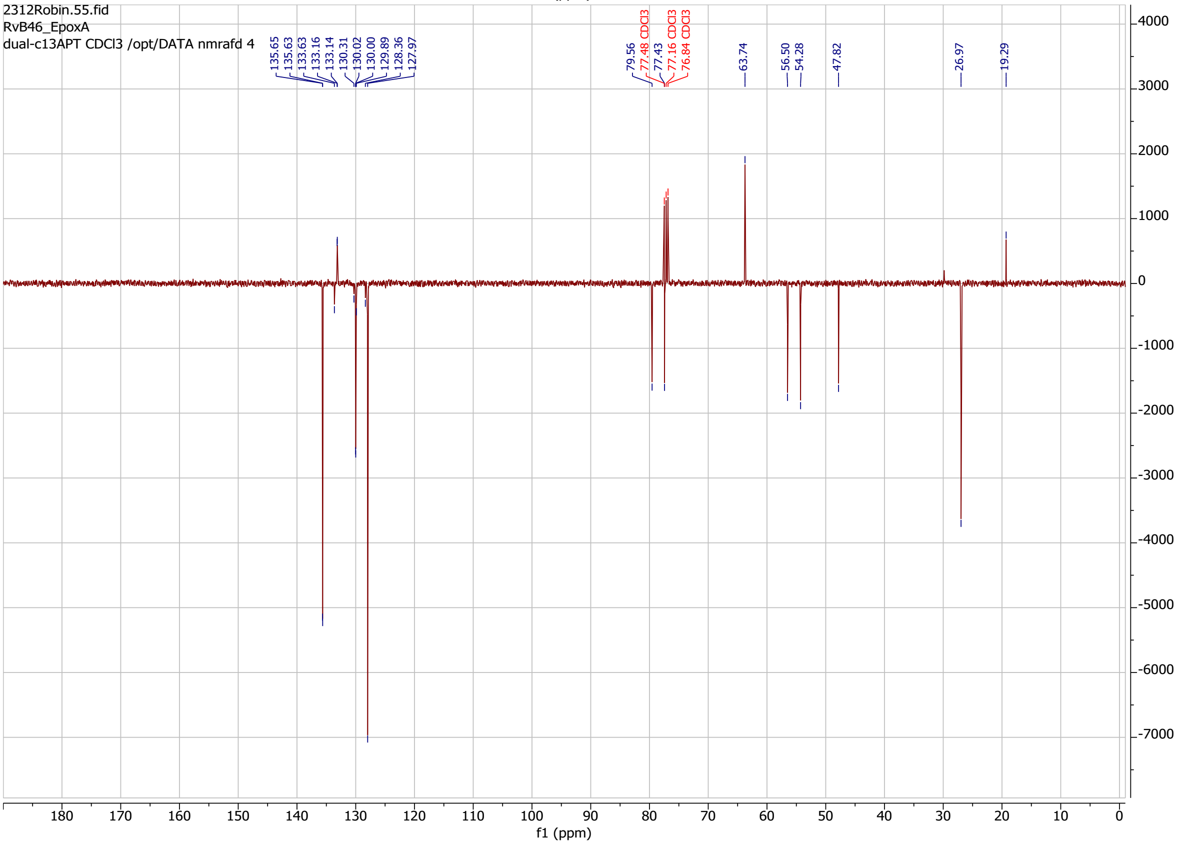

**RvB16** in $CDCl_3$

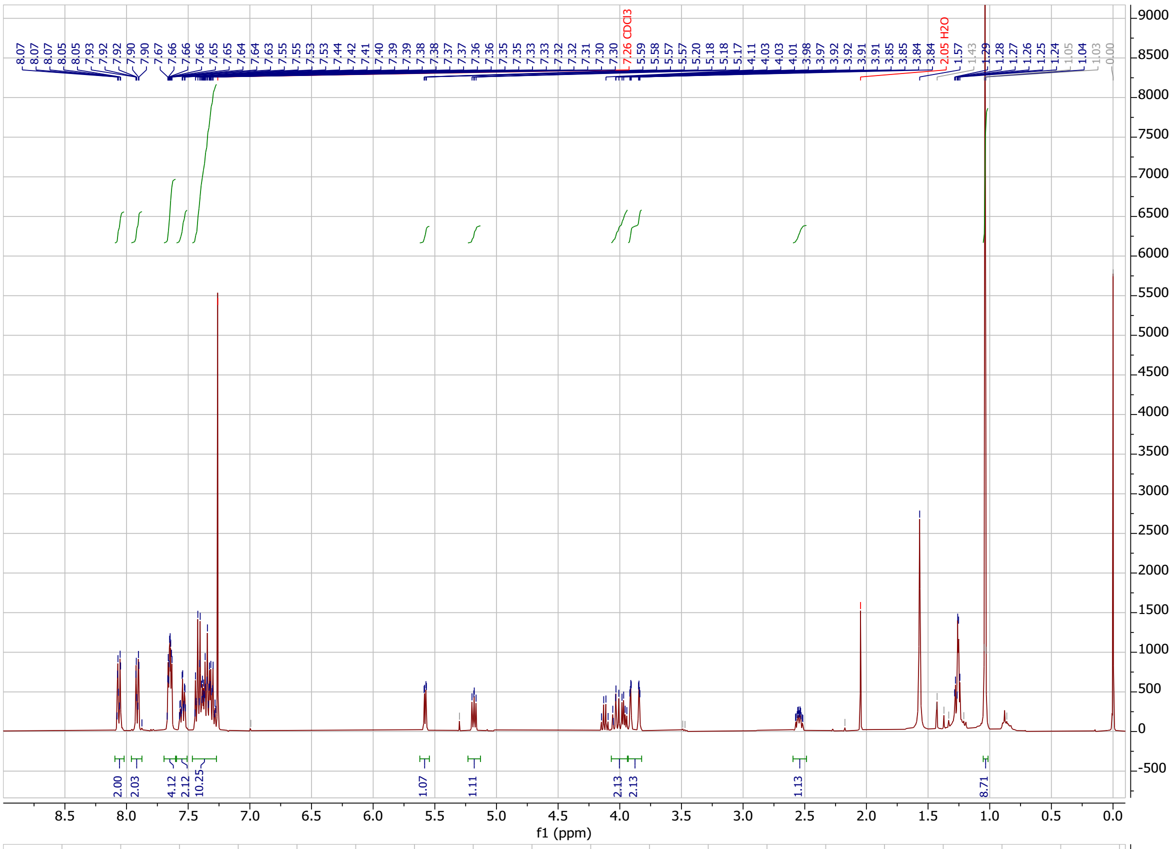

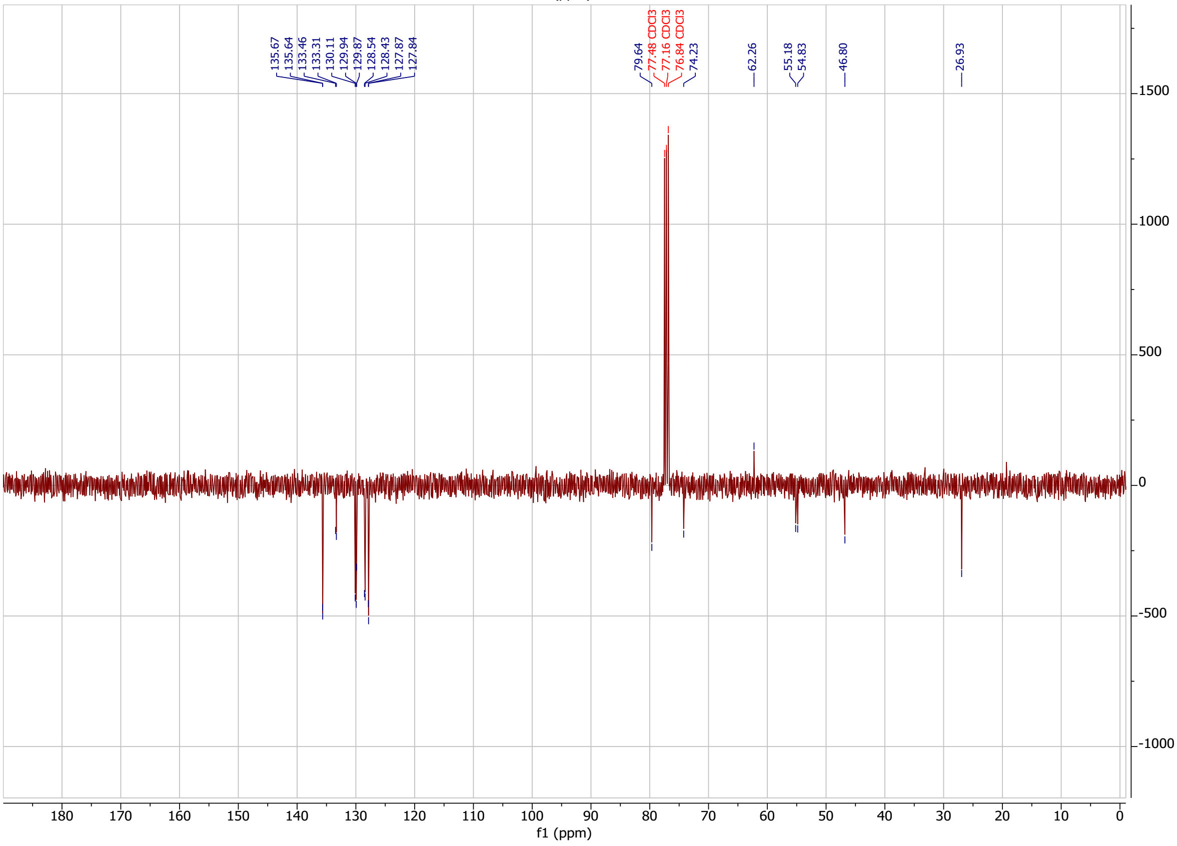

**RvB17 & RvB18** in $CDCl_3$

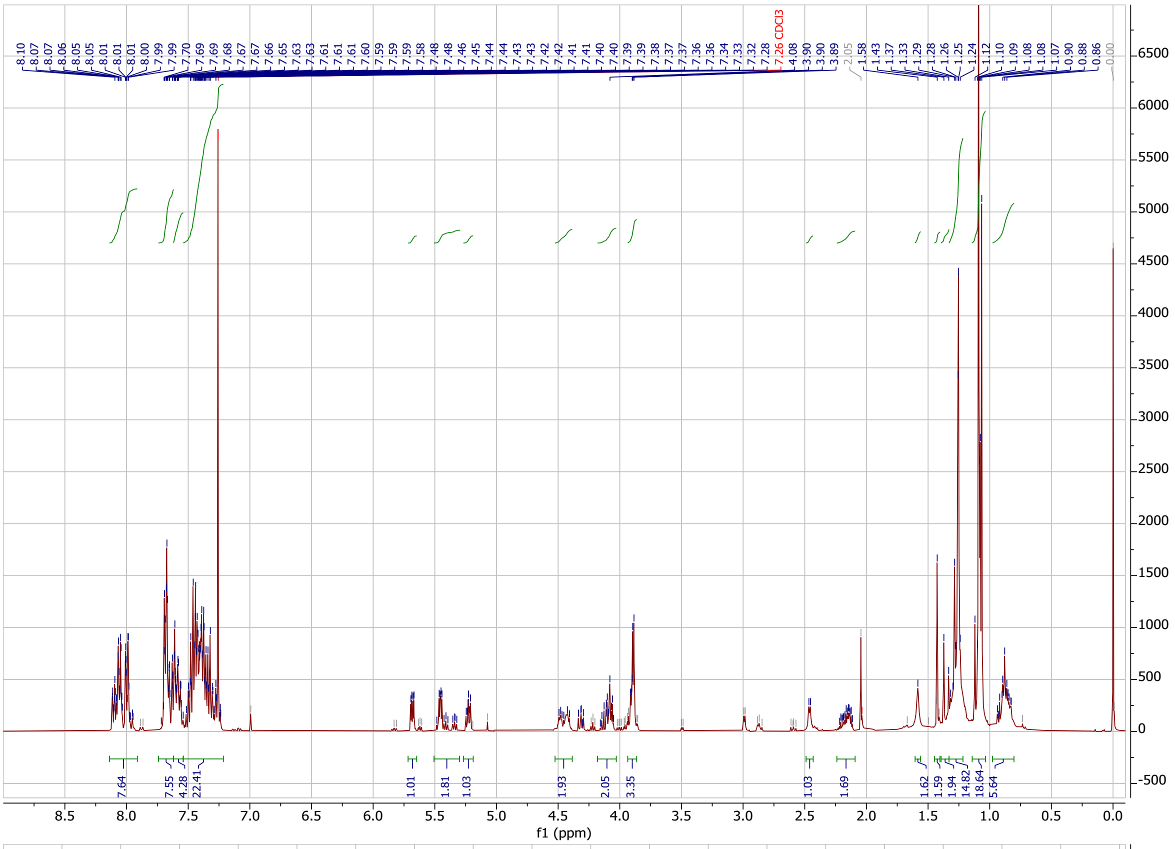

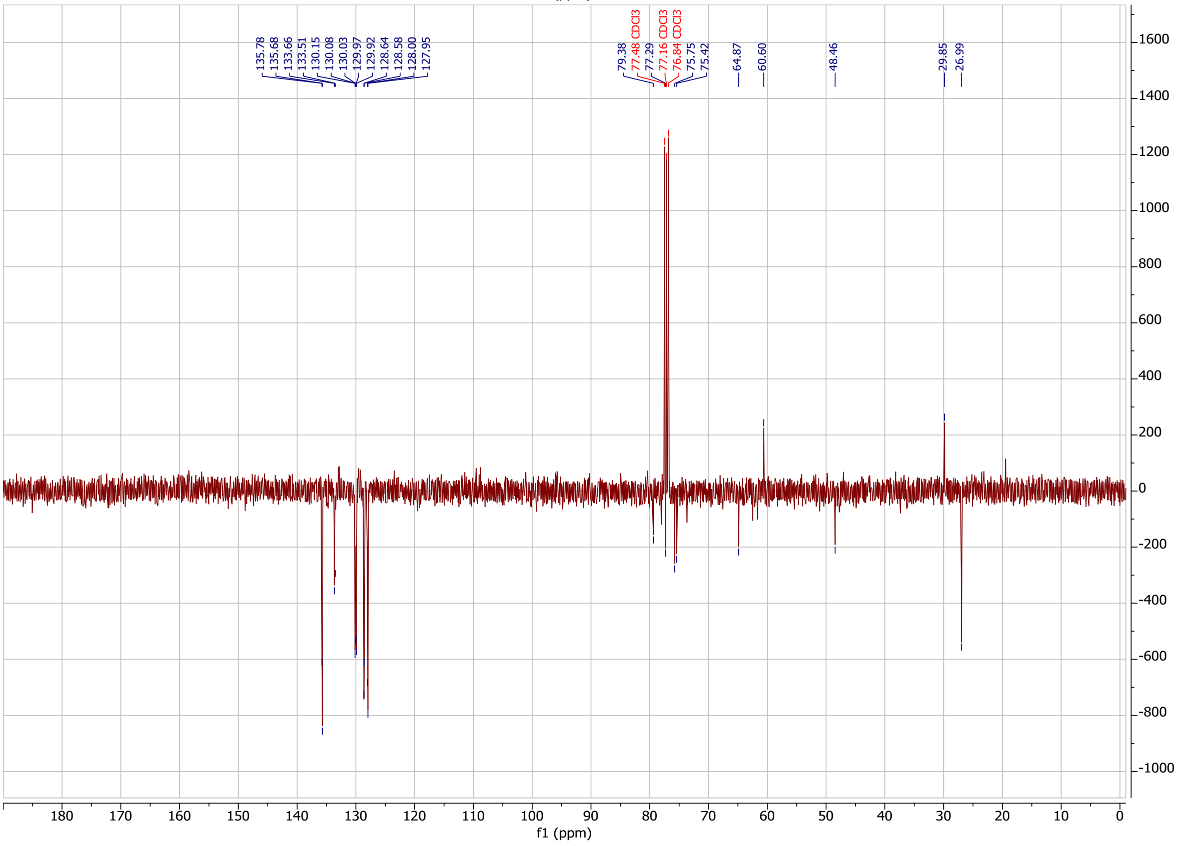

**RvB19 & RvB19** in $CDCl_3$

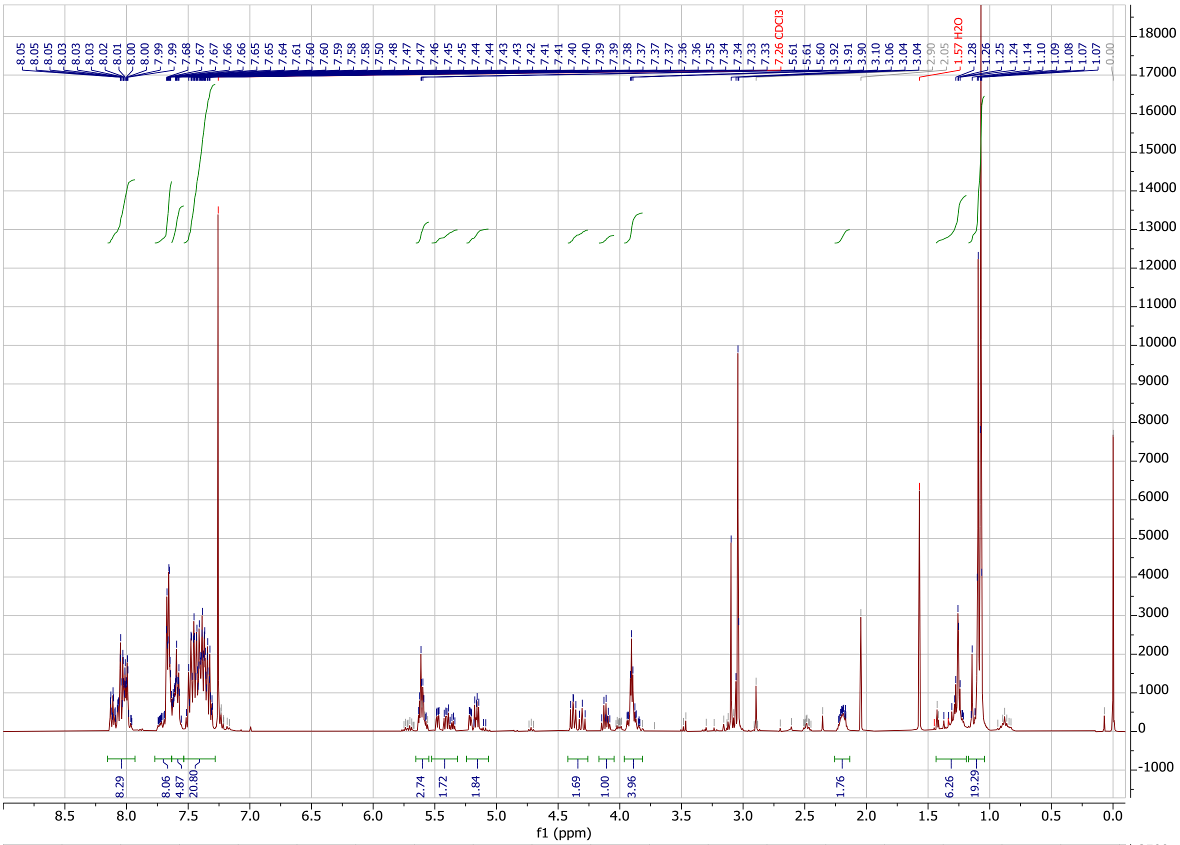

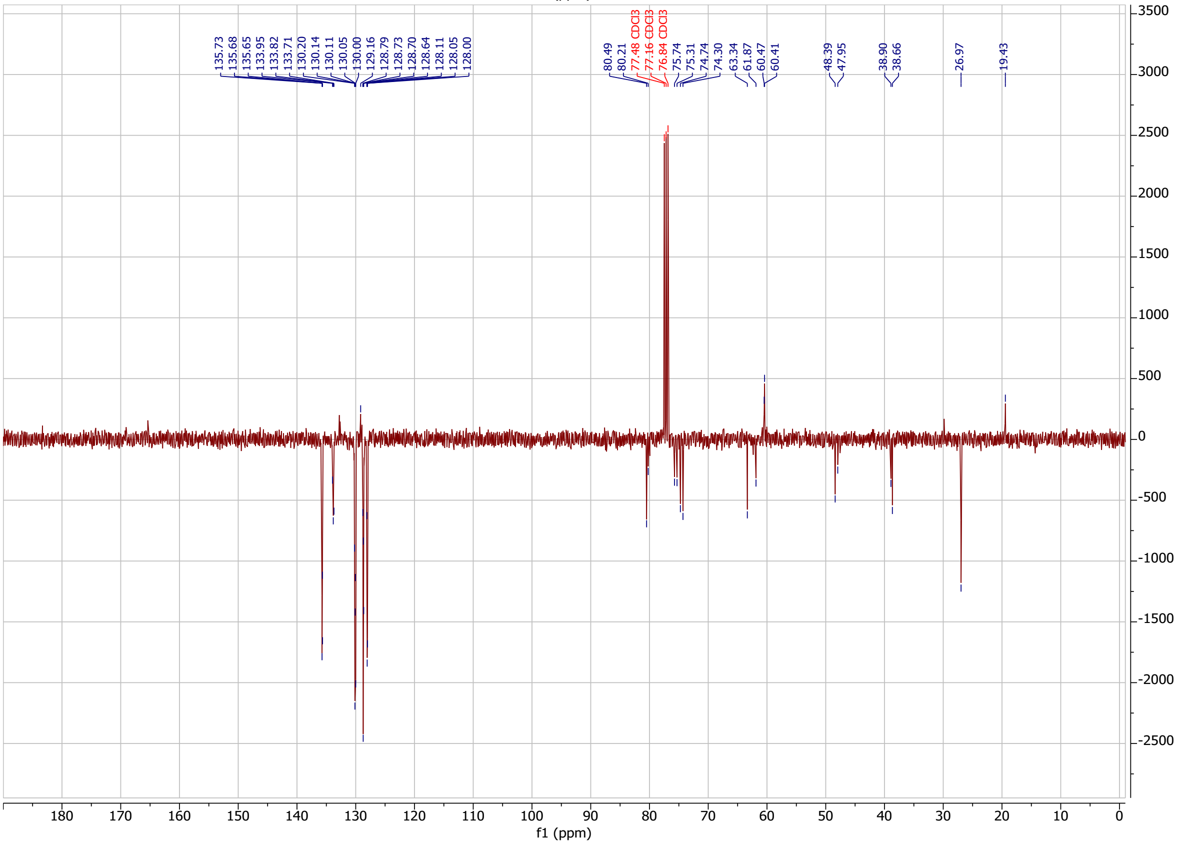

**RvB21** in $CDCl_3$

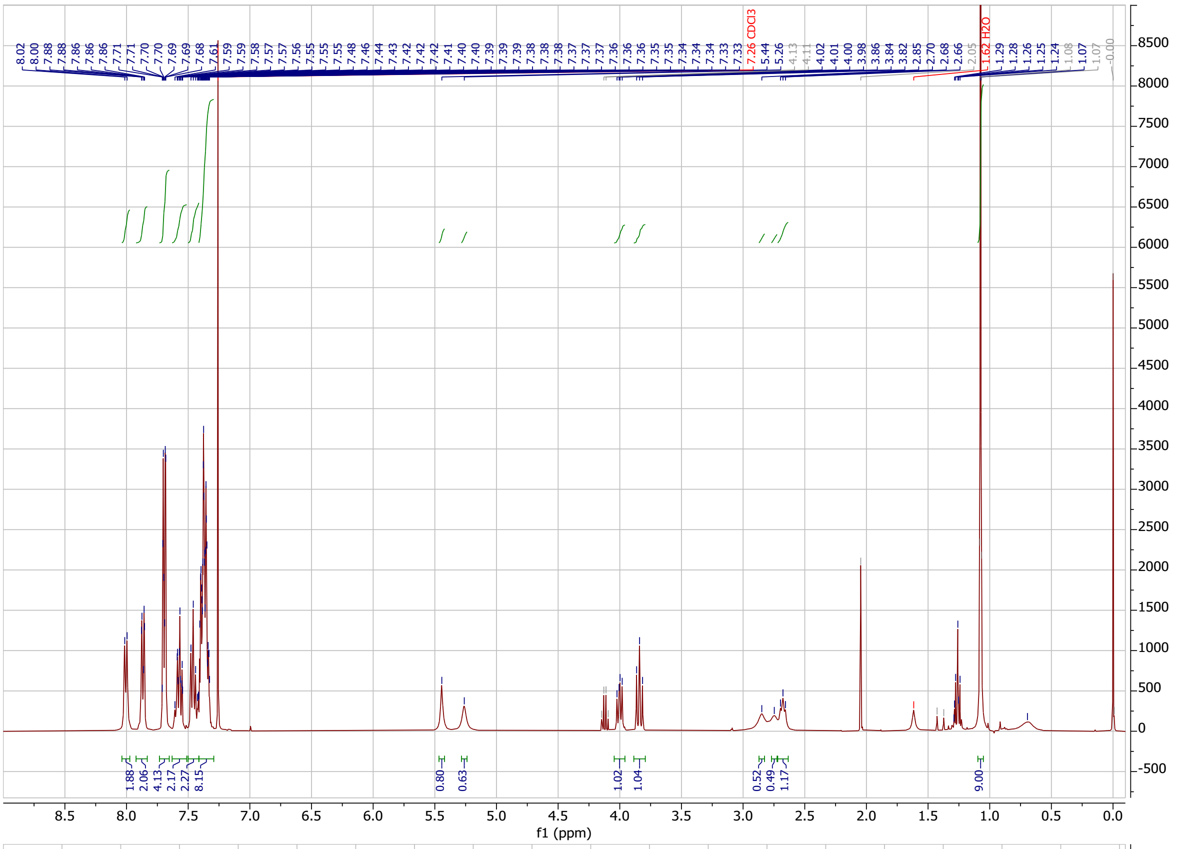

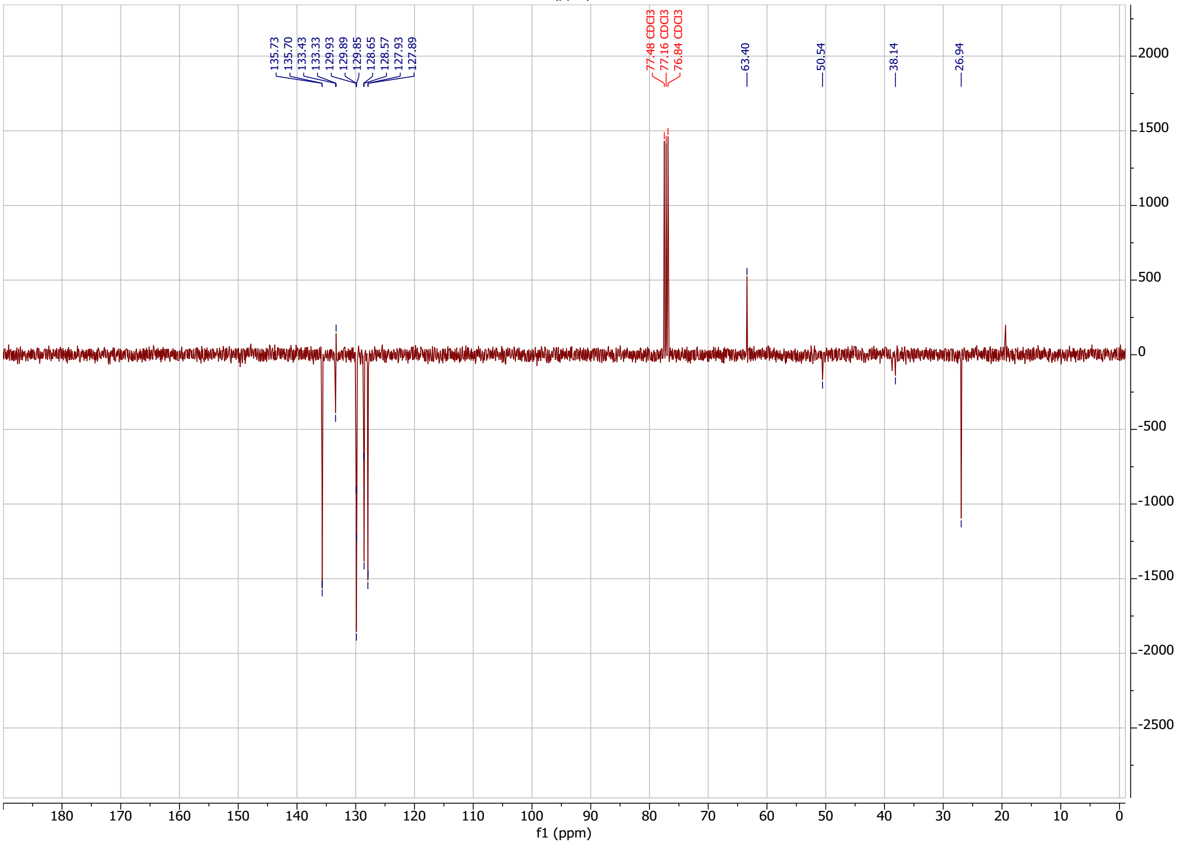

**MH367** in $CDCl_3$

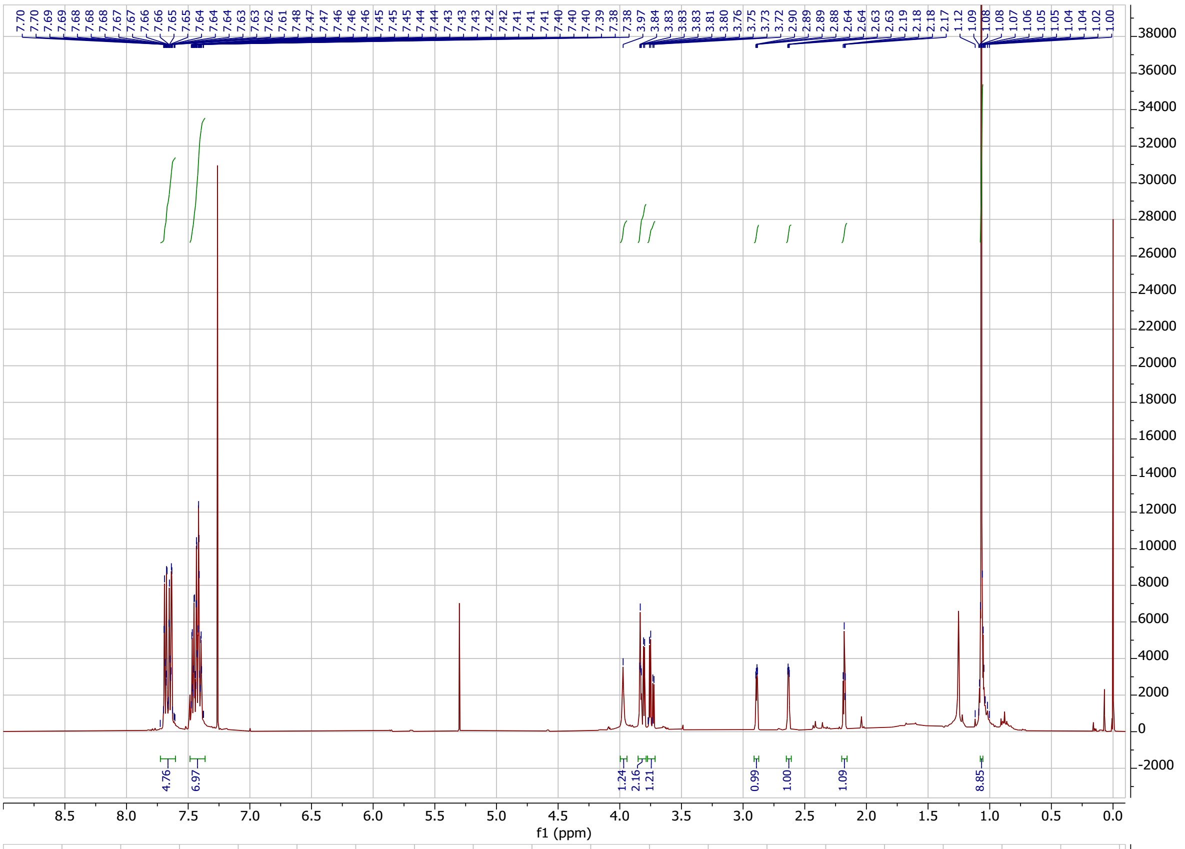

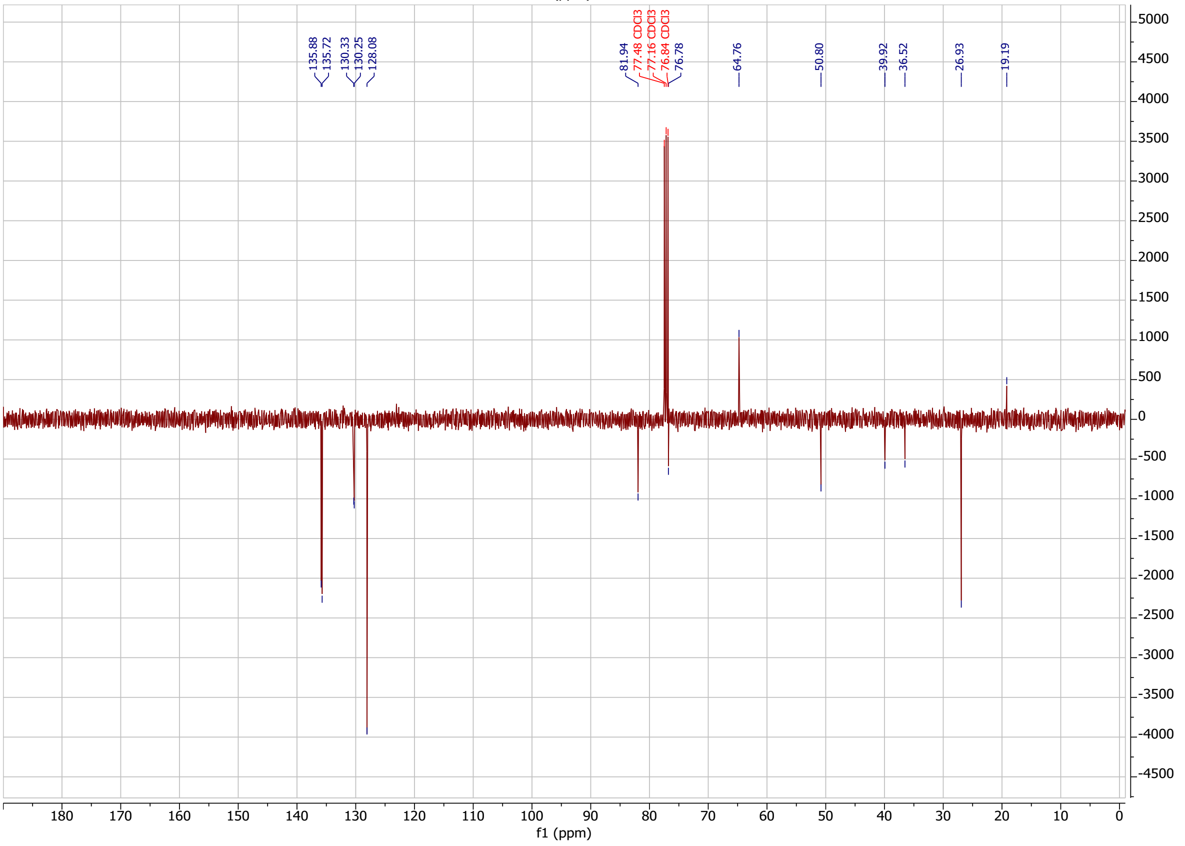

**MH371** in $CDCl_3$

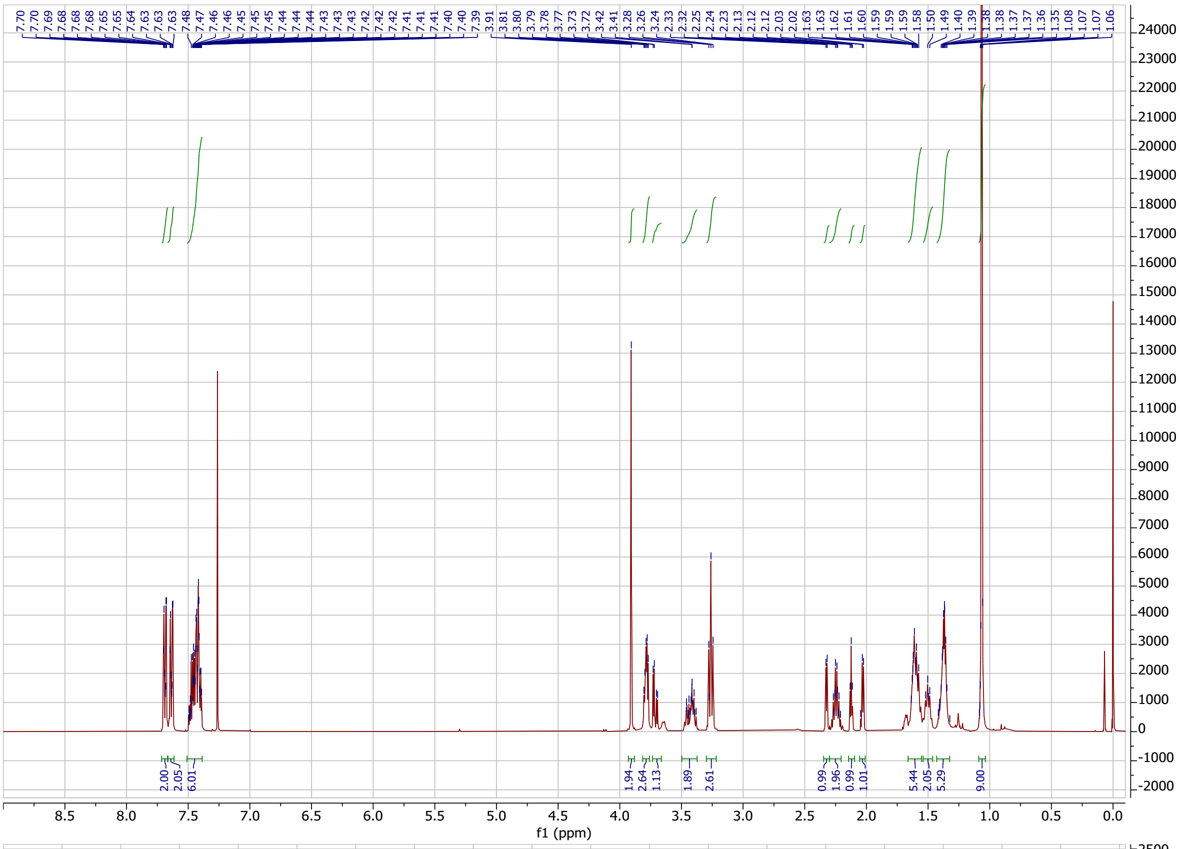

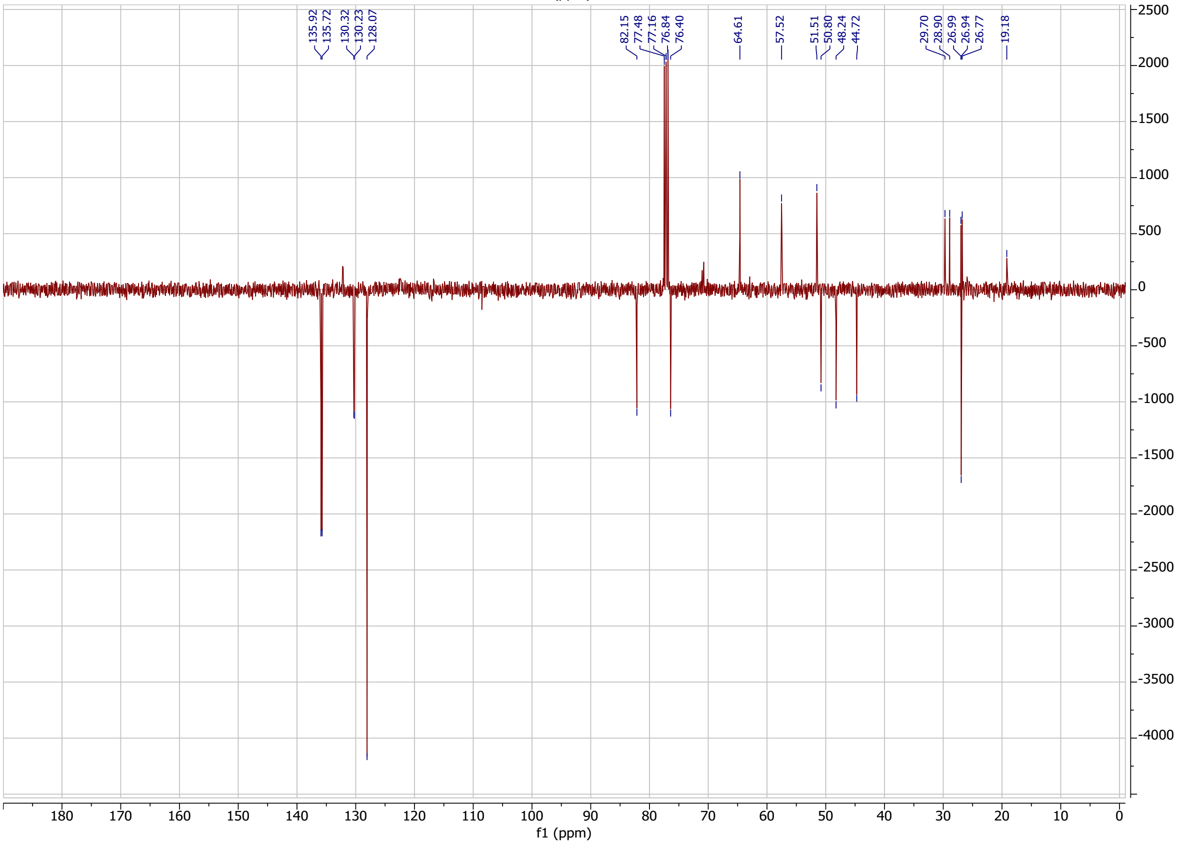

**MH373** in MeOD

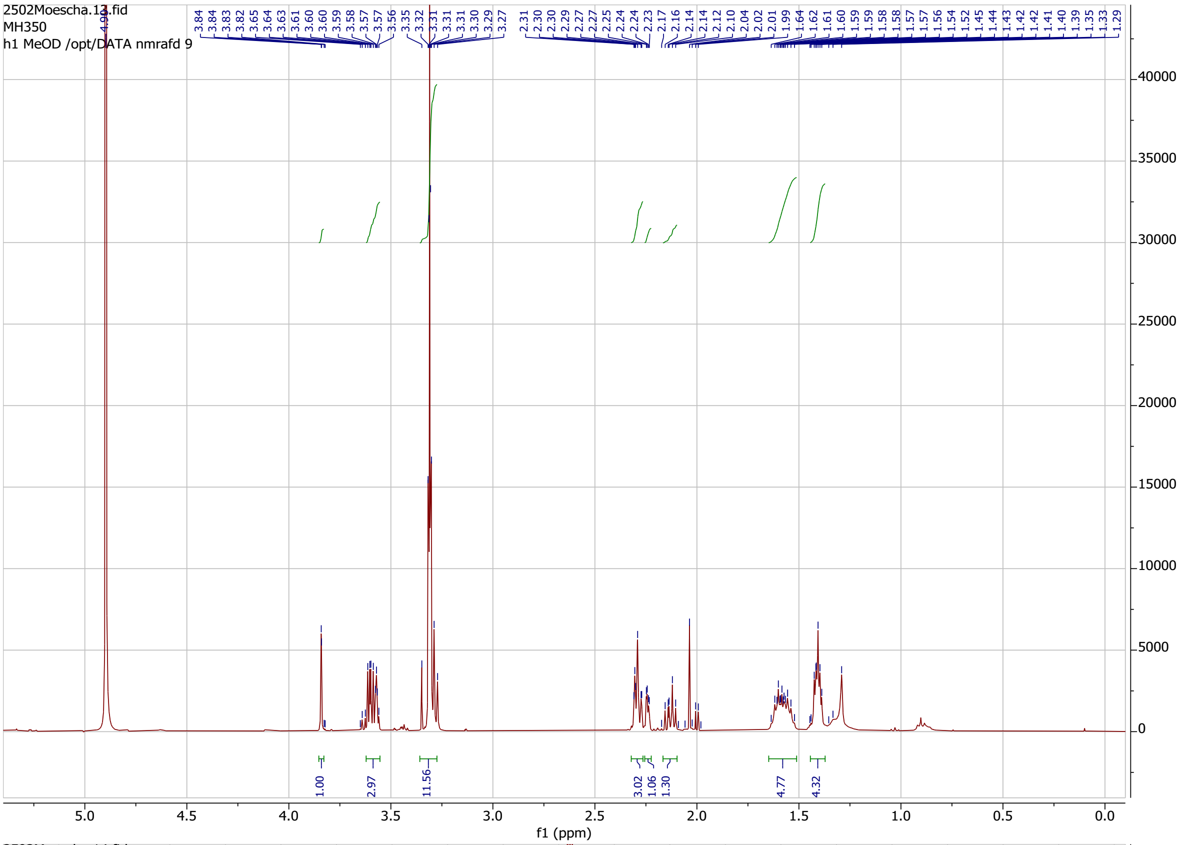

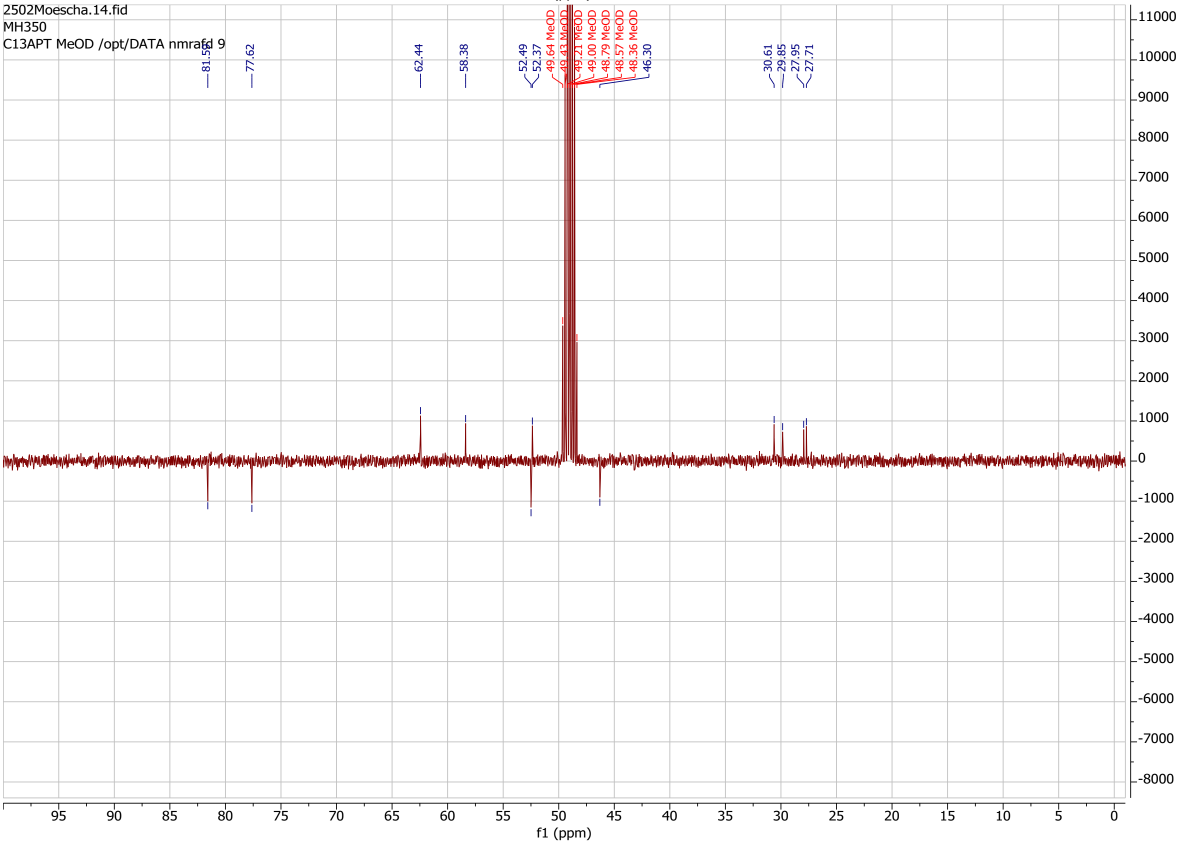

**MH374** in MeOD

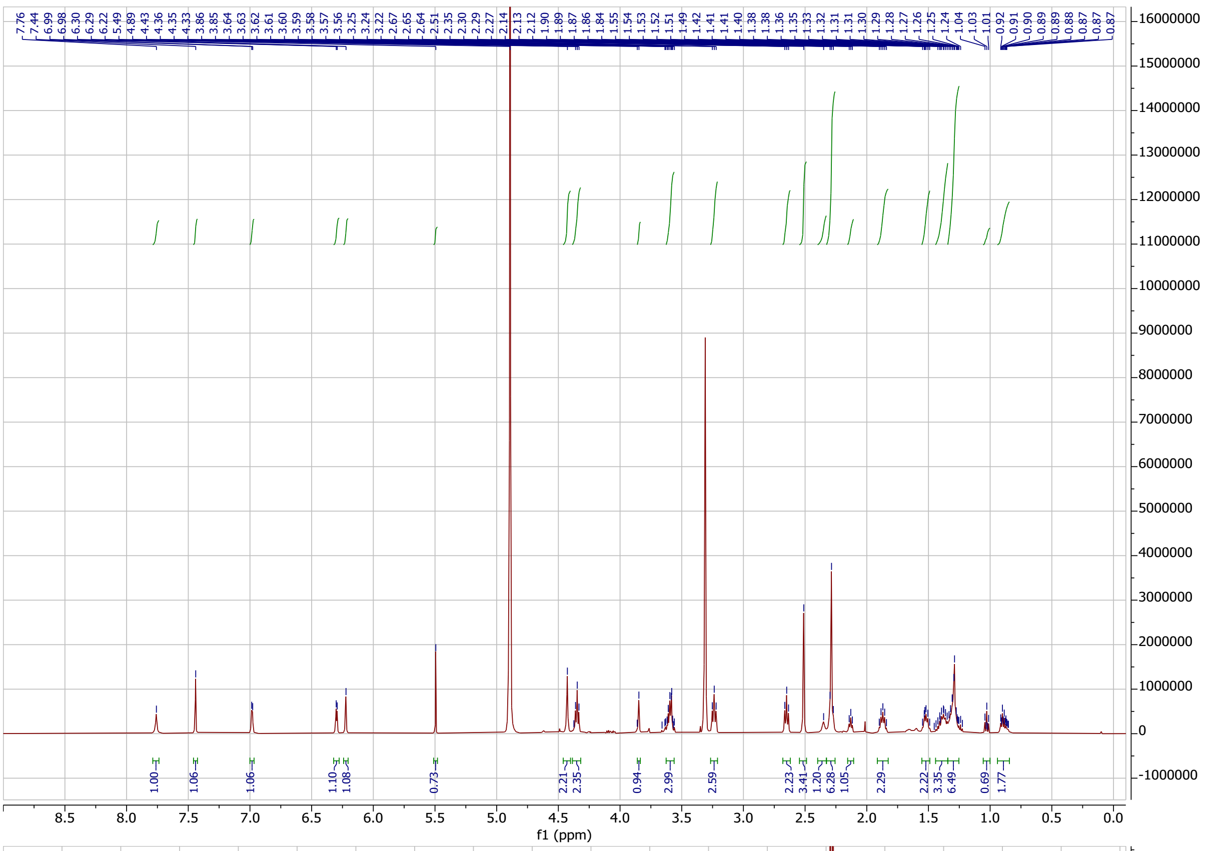

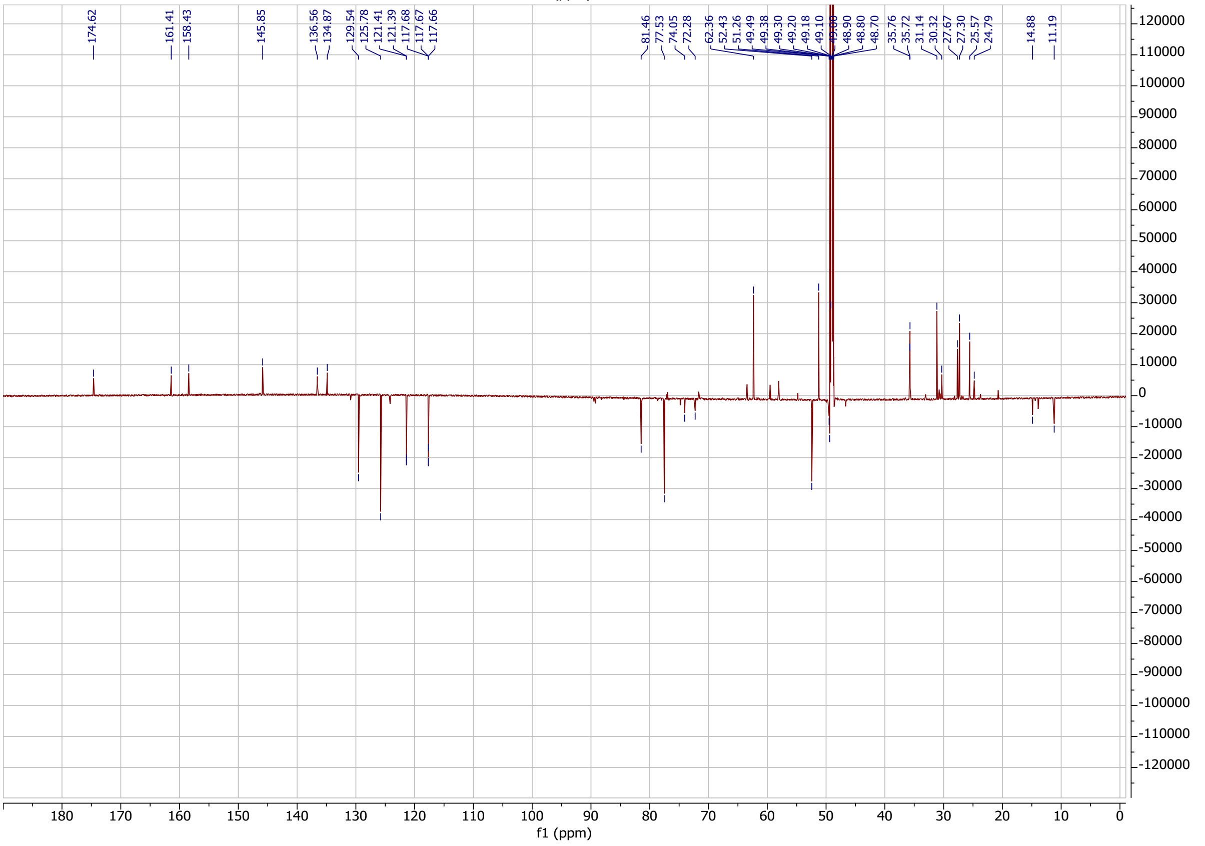

**FK23-037** in $CDCl_3$

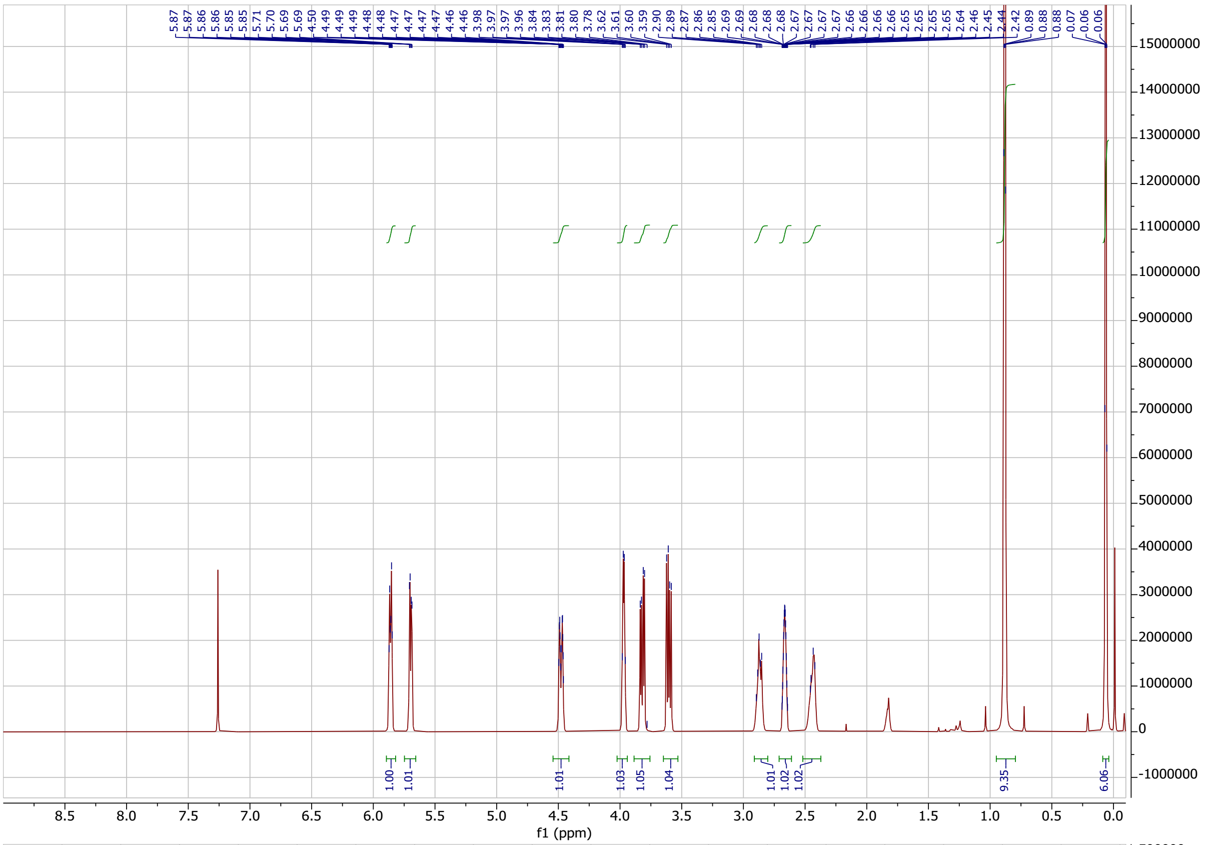

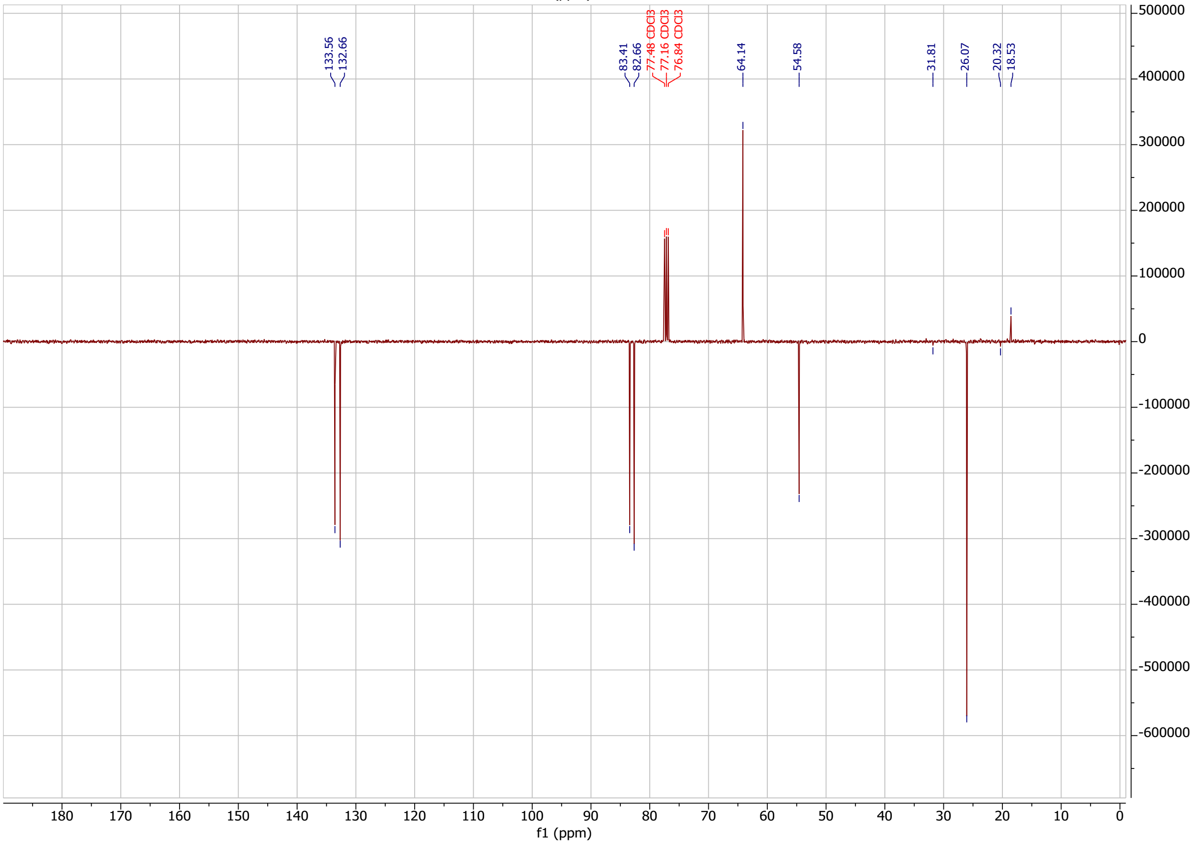

**FK23-043** in $CDCl_3$

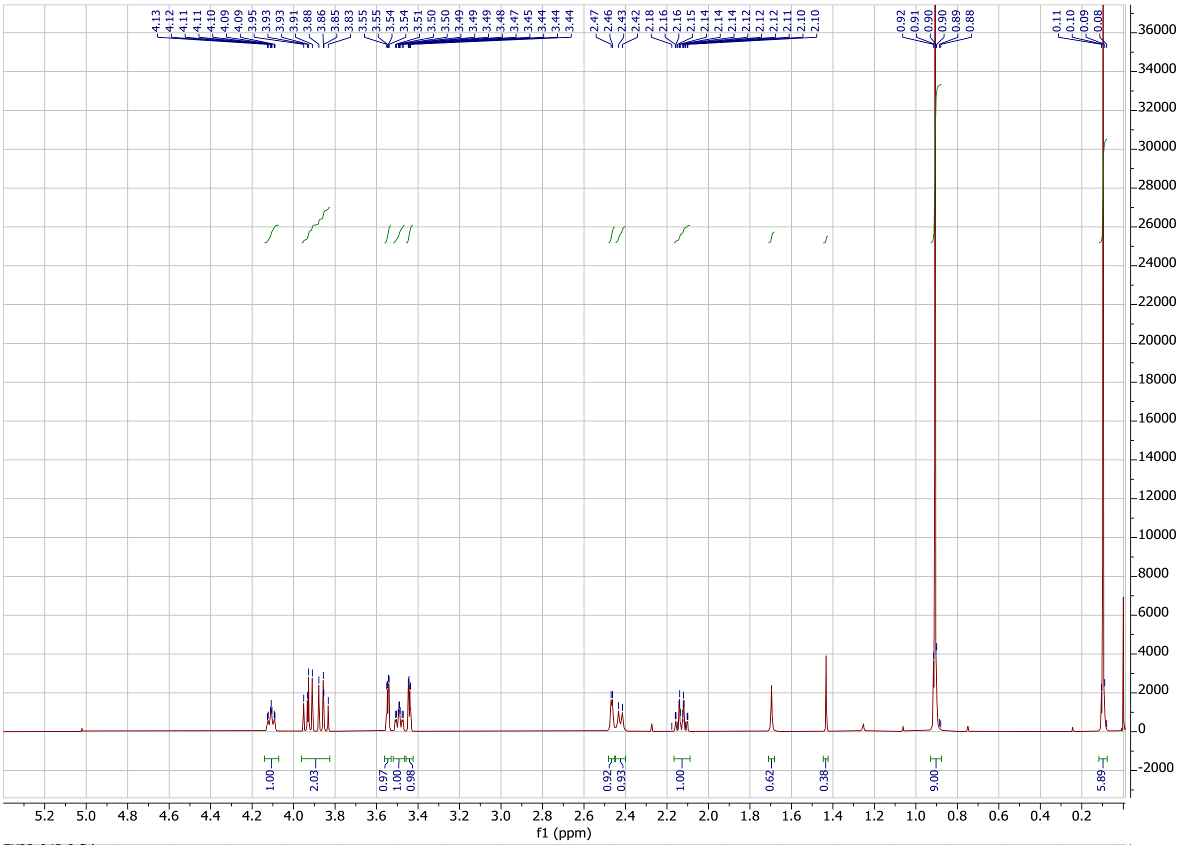

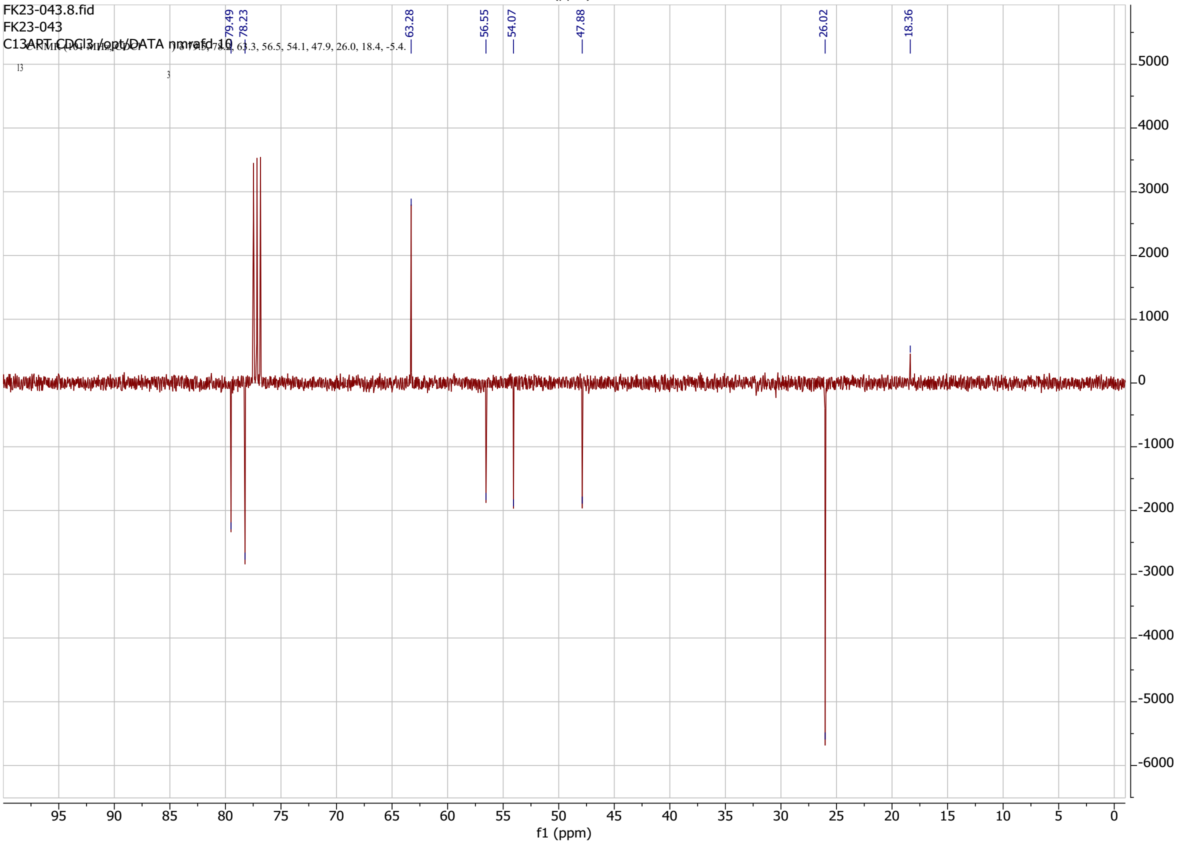

**FK-AH017** in MeOD

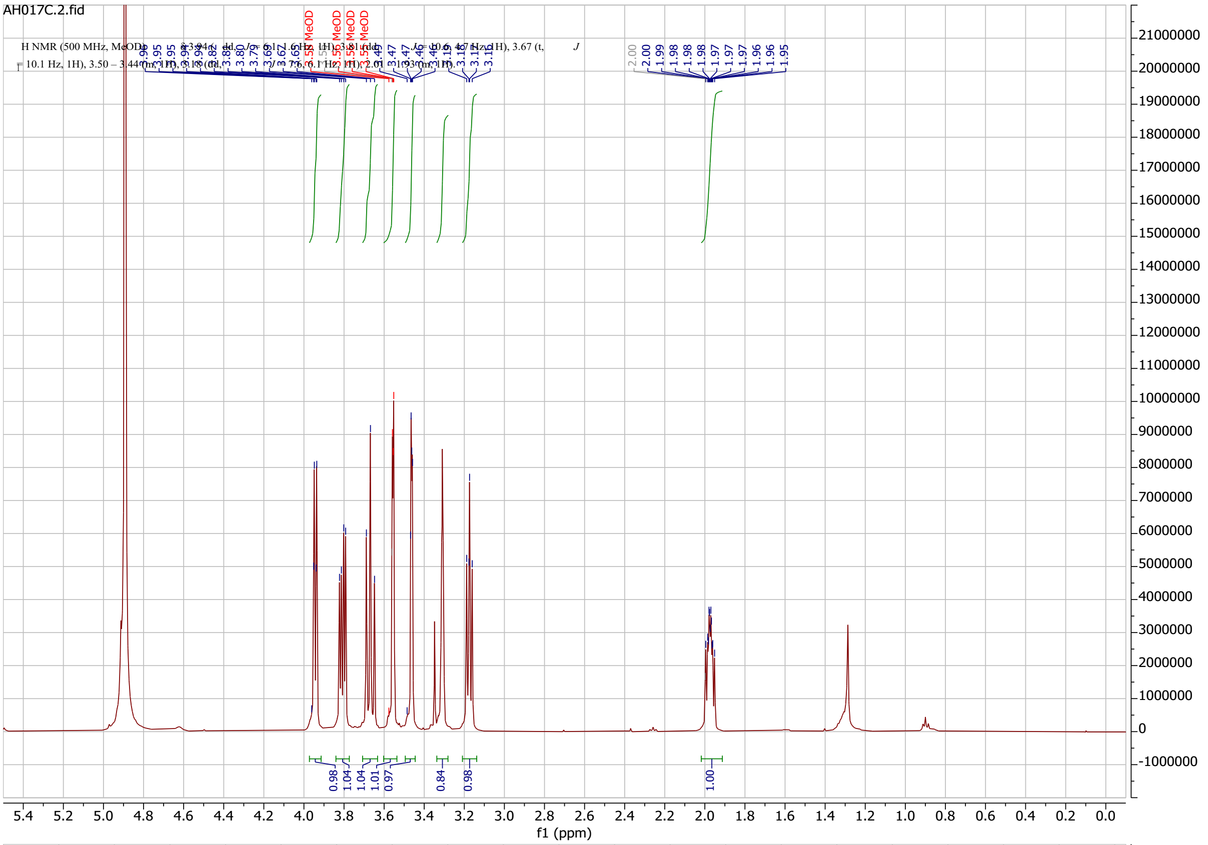

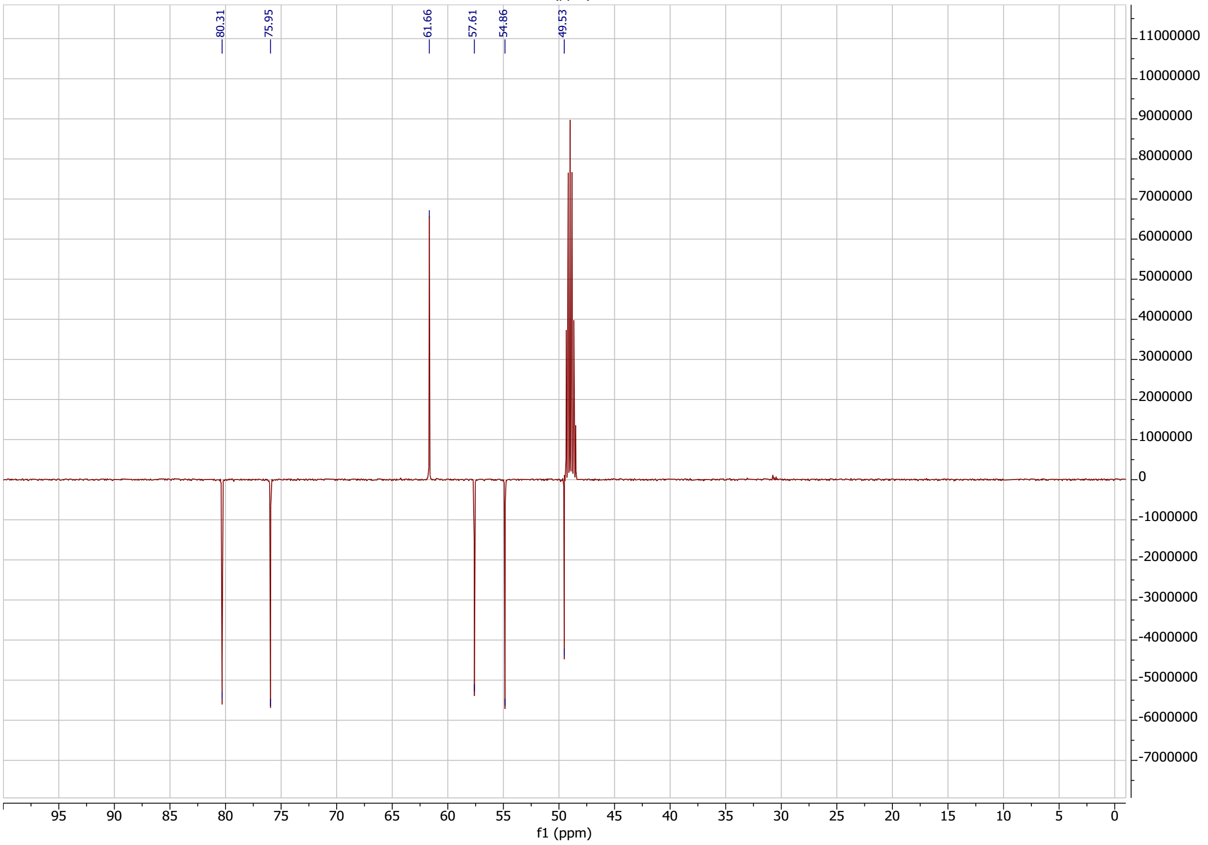

**FK23-045** in $CDCl_3$

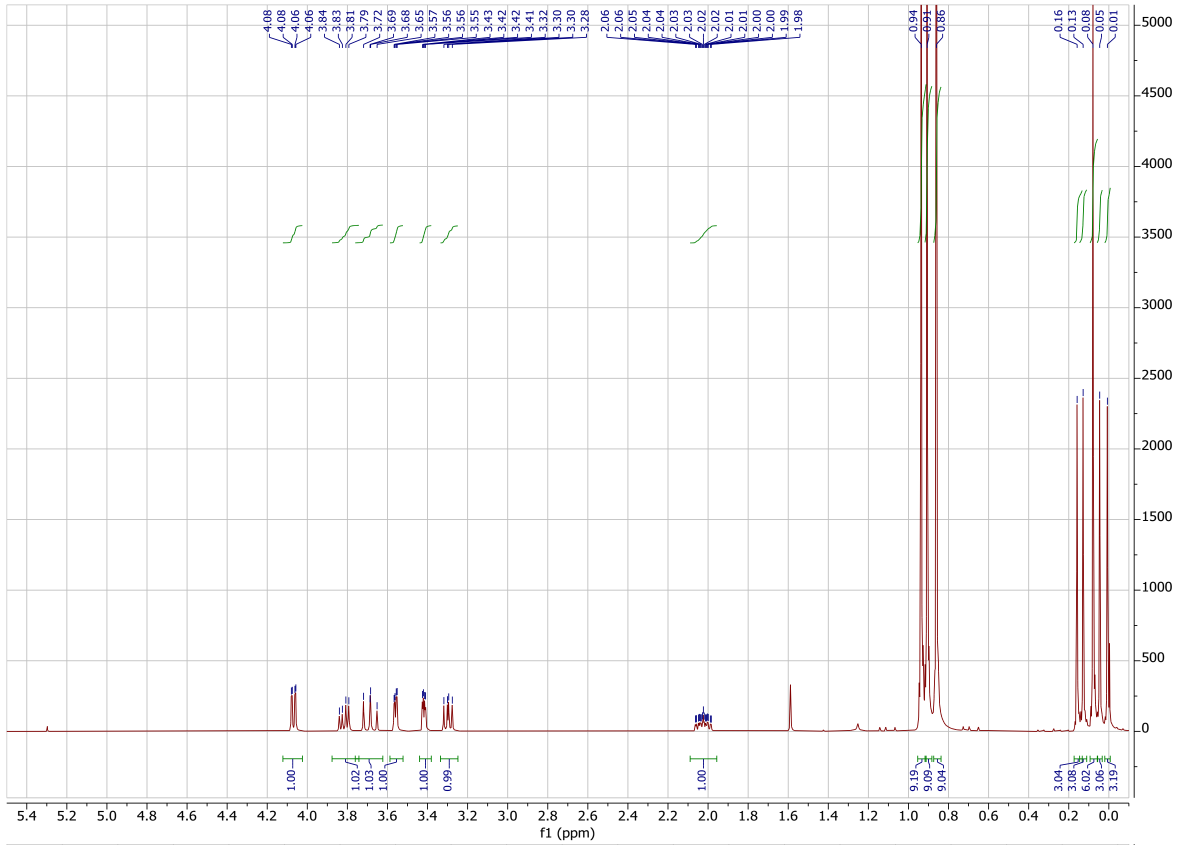

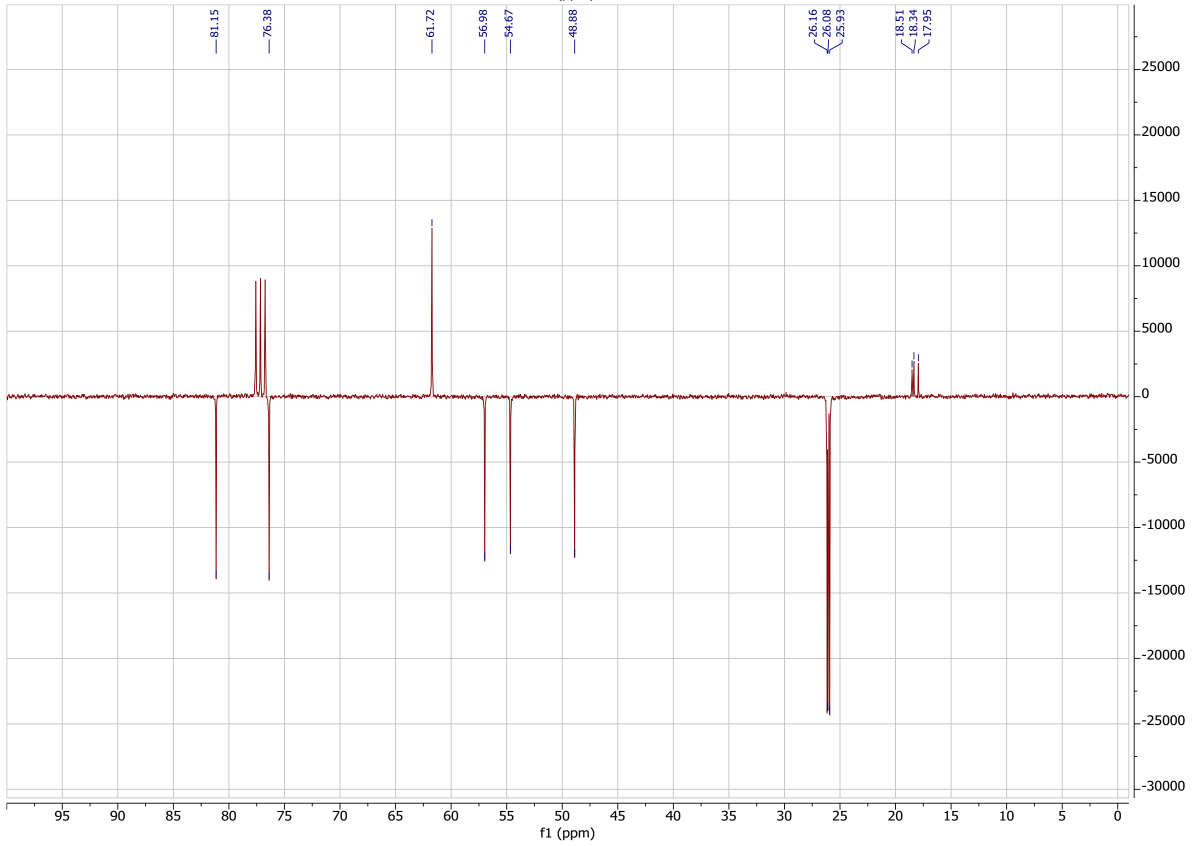

**FK23-111** in $CDCl_3$

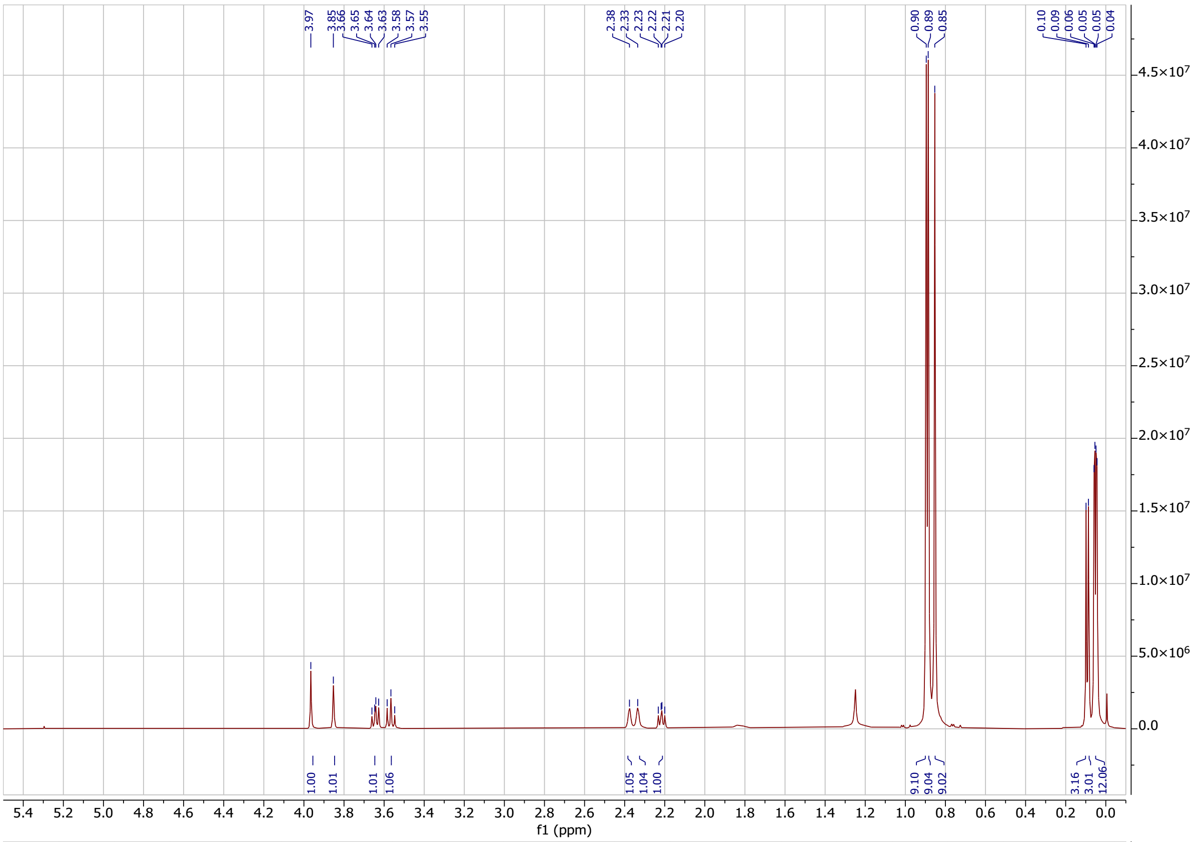

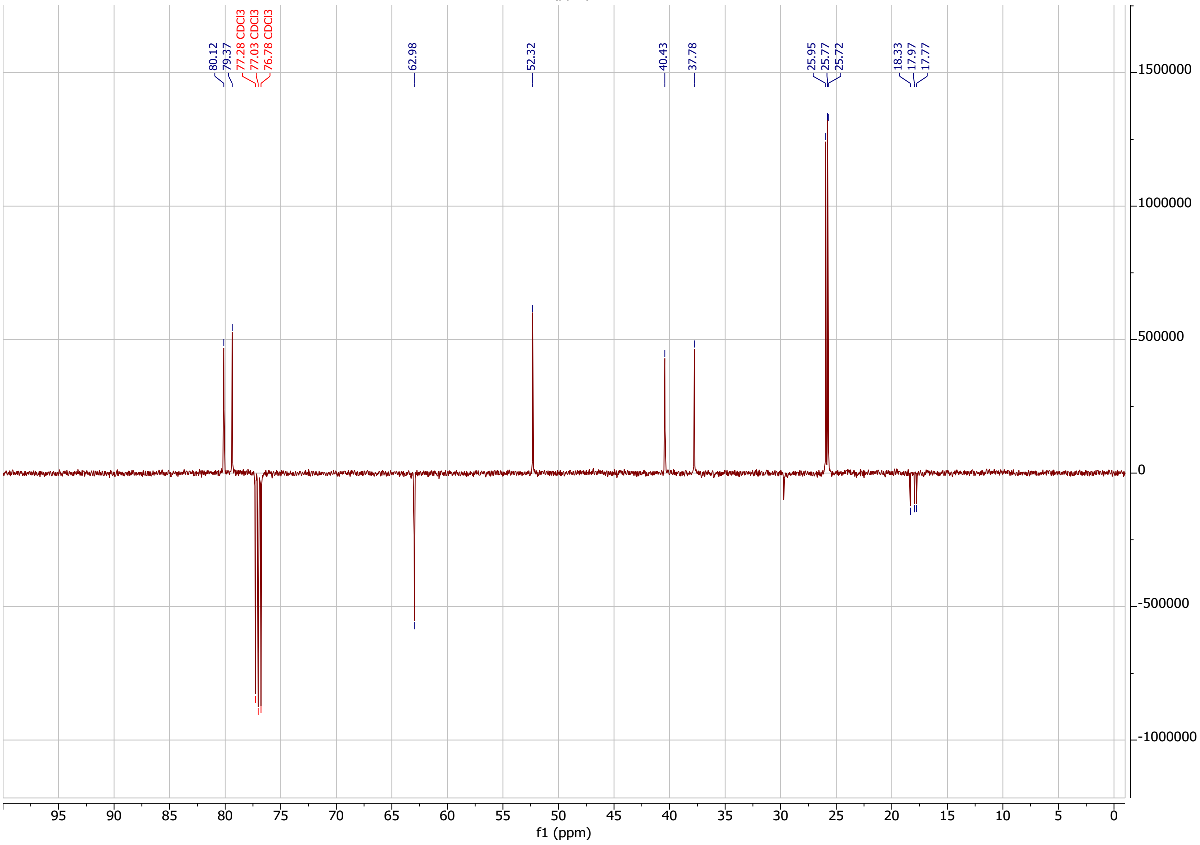

**FK23-115** in $D_2O$

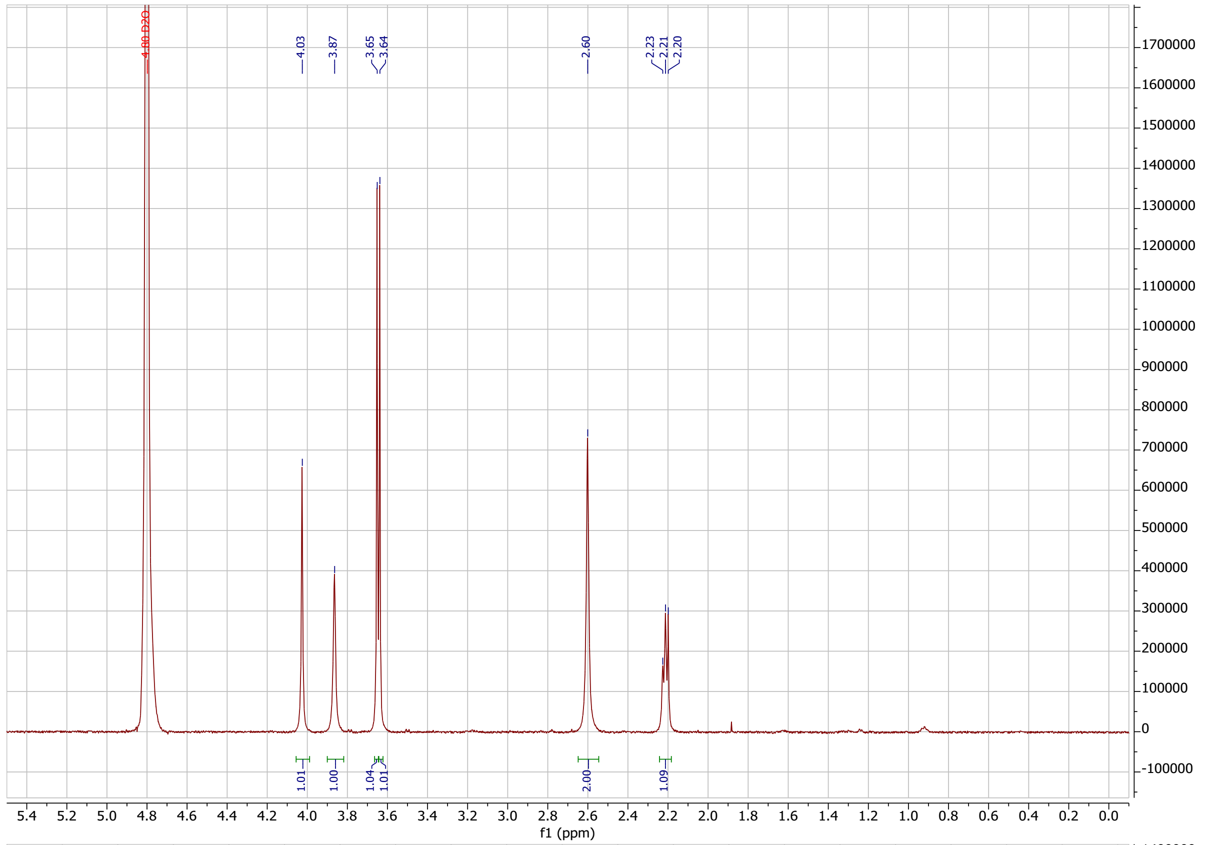

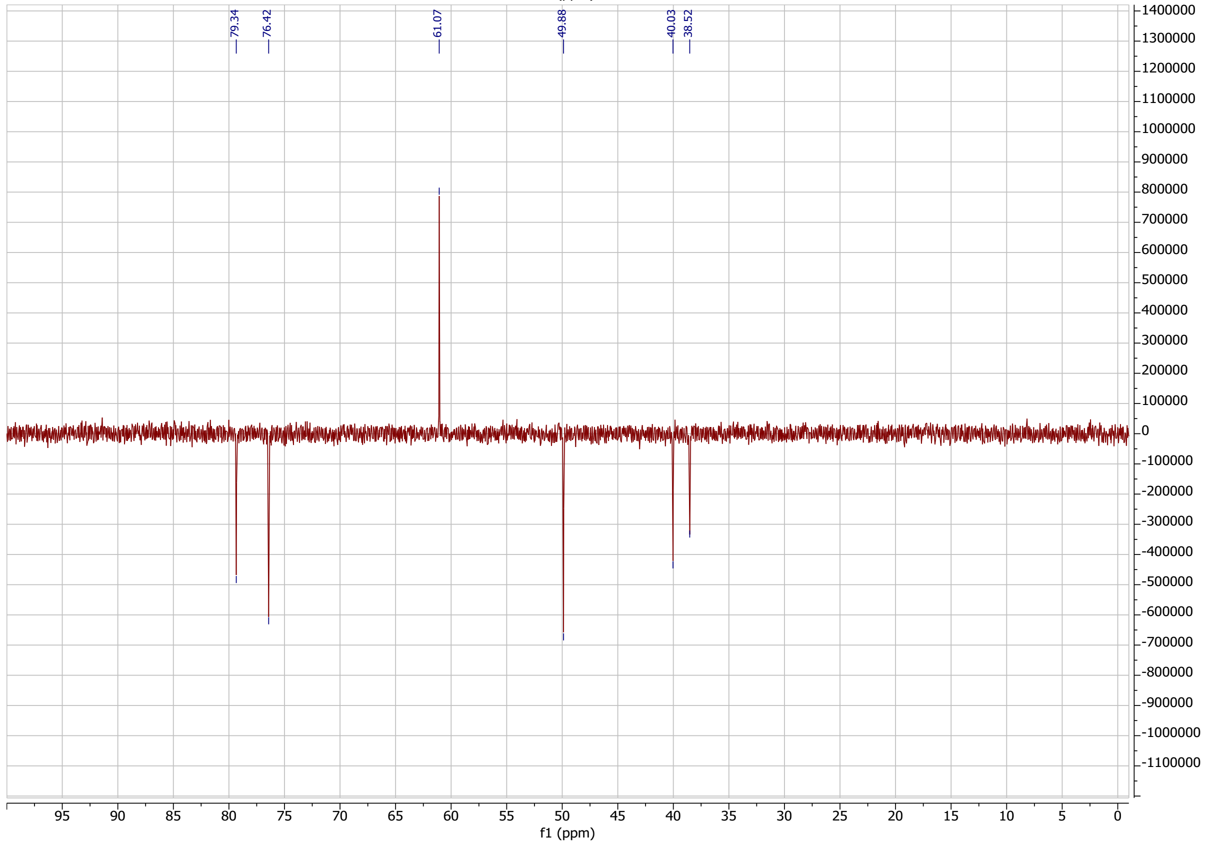
